# *De-Novo*-Designed Translational Repressors for Multi-Input Cellular Logic

**DOI:** 10.1101/501783

**Authors:** Jongmin Kim, Yu Zhou, Paul Carlson, Mario Teichmann, Friedrich C. Simmel, Pamela A. Silver, James J. Collins, Julius B. Lucks, Peng Yin, Alexander A. Green

## Abstract

Synthetic biology aims to apply engineering principles toward the development of novel biological systems for biotechnology and medicine. Despite efforts to expand the set of high-performing parts for genetic circuits, achieving more complex circuit functions has often been limited by the idiosyncratic nature and crosstalk of commonly utilized parts. Here, we present a molecular programming strategy that implements RNA-based repression of translation using *de-novo*-designed RNAs to realize high-performance orthogonal parts with mRNA detection and multi-input logic capabilities. These synthetic post-transcriptional regulators, termed toehold repressors and three-way junction (3WJ) repressors, efficiently suppress translation in response to cognate trigger RNAs with nearly arbitrary sequences using thermodynamically and kinetically favorable linear-linear RNA interactions. Automated *in silico* optimization of thermodynamic parameters yields improved toehold repressors with up to 300-fold repression, while in-cell SHAPE-Seq measurements of 3WJ repressors confirm their designed switching mechanism in living cells. Leveraging the absence of sequence constraints, we identify eight- and 15-component sets of toehold and 3WJ repressors, respectively, that provide high orthogonality. The modularity, wide dynamic range, and low crosstalk of the repressors enable their direct integration into ribocomputing devices that provide universal NAND and NOR logic capabilities and can perform multi-input RNA-based logic. We demonstrate these capabilities by implementing a four-input NAND gate and the expression NOT((A1 AND A2) OR (B1 AND B2)) in *Escherichia coli*. These features make toehold and 3WJ repressors important new classes of translational regulators for biotechnological applications.

## INTRODUCTION

Synthetic biology aims to provide an engineering-driven approach towards building biological entities with complex and novel functionality using well-characterized modular parts. Precise and programmable control of gene expression is becoming a basic requirement for many synthetic biology applications as well as biotechnology in general. For example, the ability to control the expression of multiple genes will aid in optimization of biosynthetic pathways for industrial chemical production while maximizing productivity and minimizing toxicity to host organisms^1^. Moreover, continued advances in gene regulation provide crucial new capabilities for emerging synthetic biology applications in areas such as cell-based therapies^2^, advanced biomaterials^3^, and green chemistry^4^.

Starting with the first demonstrations of toggle switches^5^ and oscillators^6^ in *Escherichia coli*, synthetic biology approaches have yielded novel and increasingly sophisticated gene circuitry over the years including synchronized oscillators^7,8^, logic gates^9-12^, memory devices^13,14^, analog signal processors^15,16^, and state machines^17^. Bacteria enhanced with engineered genetic circuits have found use as memory devices in the mammalian gut environment^18^ and have been deployed against tumors in mouse models^19^. Cancer cell classifiers^20^ have further expanded the utility of synthetic circuitry for biomedical applications. Despite these advances, many of the regulatory elements in previous work overlap or are not compatible with each other, thereby limiting the ability to integrate such diverse components and elementary systems towards more complex circuits for biotechnological applications.

A basic requirement for engineering complex systems is a large repertoire of regulators that are modular, programmable, homogeneous, predictable, and easy to compose. Though there have been numerous reports describing the mining of regulatory elements from natural organisms^13,21^, such mined parts require further scrutiny to assess their dynamic range, compatibility, and crosstalk. The idiosyncratic nature of many protein regulatory elements presents challenges in their use in circuits with complex regulatory behavior. Recently, these challenges were addressed for protein-based regulatory parts by using insulation and computer-aided design strategies to scale up the complexity of combinatorial^22^ and sequential^23^ logic circuits in bacteria. In mammalian cells, the highly specific DNA-modifying activities of recombinases were used to implement over 100 genetic circuits with minimal optimization^24^, while composable viral proteases were used to construct a variety of protein-only circuits^25^.

Due to their simple base-pairing rules and well-characterized thermodynamic parameters, RNA molecules provide an alternative means to construct genetic circuits with advantages in terms of programmability, predictability, and composability compared to most proteins. In a seminal study^26^, RNA base-pairing design rules were used to create multiple orthogonal sense-antisense RNA partners that regulate translation. Building on the natural pT181 transcriptional attenuator system, a set of orthogonal mutants of a natural antisense RNA–mediated transcriptional attenuation system were successfully demonstrated^27,28^. Further, a number of synthetic RNA-based translational repressors were described by exploring orthogonal mutants starting from the well-known IS10 antisense RNA–mediated translation control system^29^. These previous reports described RNA-based repressor designs where the specificity and efficiency of the control systems can be largely explained by the thermodynamics of RNA interactions. However, in a number of instances, the strategy for creating a family of orthogonal devices was based on the design features observed in natural systems or derived from previous work, thereby failing to fully capitalize on the sequence space and design features accessible to *de-novo*-designed riboregulators.

To achieve truly synthetic *de novo* RNA regulatory elements, we have previously reported toehold switches^30^ that relied on two crucial design principles for regulation of prokaryotic translation: first, the accessibility of ribosomal binding site (RBS) and start codon was limited without resorting to direct base-pairing to either of those conserved translational elements; second, linear-linear nucleic acid interactions based on toehold-mediated strand-displacement reactions^31^ were used as the trigger mechanism instead of using loop regions to initiate interactions^27,29^. These strategic changes resulted in a large library of synthetic riboregulator parts that showed wide dynamic range, and little crosstalk, enabling sophisticated translational control *in vivo*^32^. The modular and programmable nature of toehold switches provided a general framework for their integration into multi-input logic processing ribocomputing devices^33^ and enabled their deployment in diagnostic systems^34–37^. In this previous work, however, NOT logic was only achieved by using antisense RNAs, which imposed sequence constraints and precluded efficient implementation of the universal NAND and NOR logic gates. Recent work has highlighted the importance of robust universal logic gates as building blocks for constructing complex genetic circuitry^22,23,38^. A crucial factor in the development of these systems has been the generation of large libraries of repressors that carry out NOT logic by suppressing gene expression^39,40^. Accordingly, the development of analogous libraries of high-performance RNA-based repressors could enable more efficient and complex forms of biomolecular logic.

Here we describe two new types of *de*-*novo*-designed RNA-based repressors termed toehold and three-way junction (3WJ) repressors that build on the design strategies of toehold switches to achieve effective translational inhibition. Both repressors employ toehold-mediated interactions to strongly repress translation in response to trigger RNAs with nearly arbitrary sequences, including full-length mRNAs, and can decrease gene expression in excess of 100-fold, a substantial improvement over previous RNA-based translational repressors^29^. Using an automated forward-engineering approach, we successfully employ thermodynamic considerations alone to improve the performance of the toehold repressors in a second-generation library. Furthermore, in-cell SHAPE-Seq measurements are used to directly confirm the formation of three-way junction structures in the 3WJ repressors following binding of the trigger RNAs. Validated high-performance repressors with low crosstalk are integrated into ribocomputing devices to achieve NOR and NAND logic with up to four sequence-independent input RNAs, providing universal building blocks for logical computation. Overall, this study reports a valuable set of new high-performance riboregulators for achieving complex gene regulation and demonstrates that computer-driven design and ribocomputing strategies can be successfully applied towards achieving robust translational repression.

## RESULTS

### Design of Synthetic Translational Repressors

Previously, we developed toehold switches that repress translation using a hairpin secondary structure that sequesters the RBS and start codon within a hairpin loop and stem, respectively (Figure 1a). A single-stranded toehold domain **a\*** at the 5′ end of the switch RNA hairpin provides the initial binding site for a single-stranded trigger RNA strand, which has a complementary domain **a**. Upon binding of the cognate trigger molecule to the hairpin and completion of a toehold-mediated branch migration, the RBS and start codon are freely available for ribosome binding and translation of the downstream gene. The lack of sequence constraints in designing trigger RNA molecules greatly expanded the set of orthogonal toehold switches and the use of thermodynamically and kinetically favorable toehold-mediated interactions provided wide dynamic range^32^.

**Figure 1.**
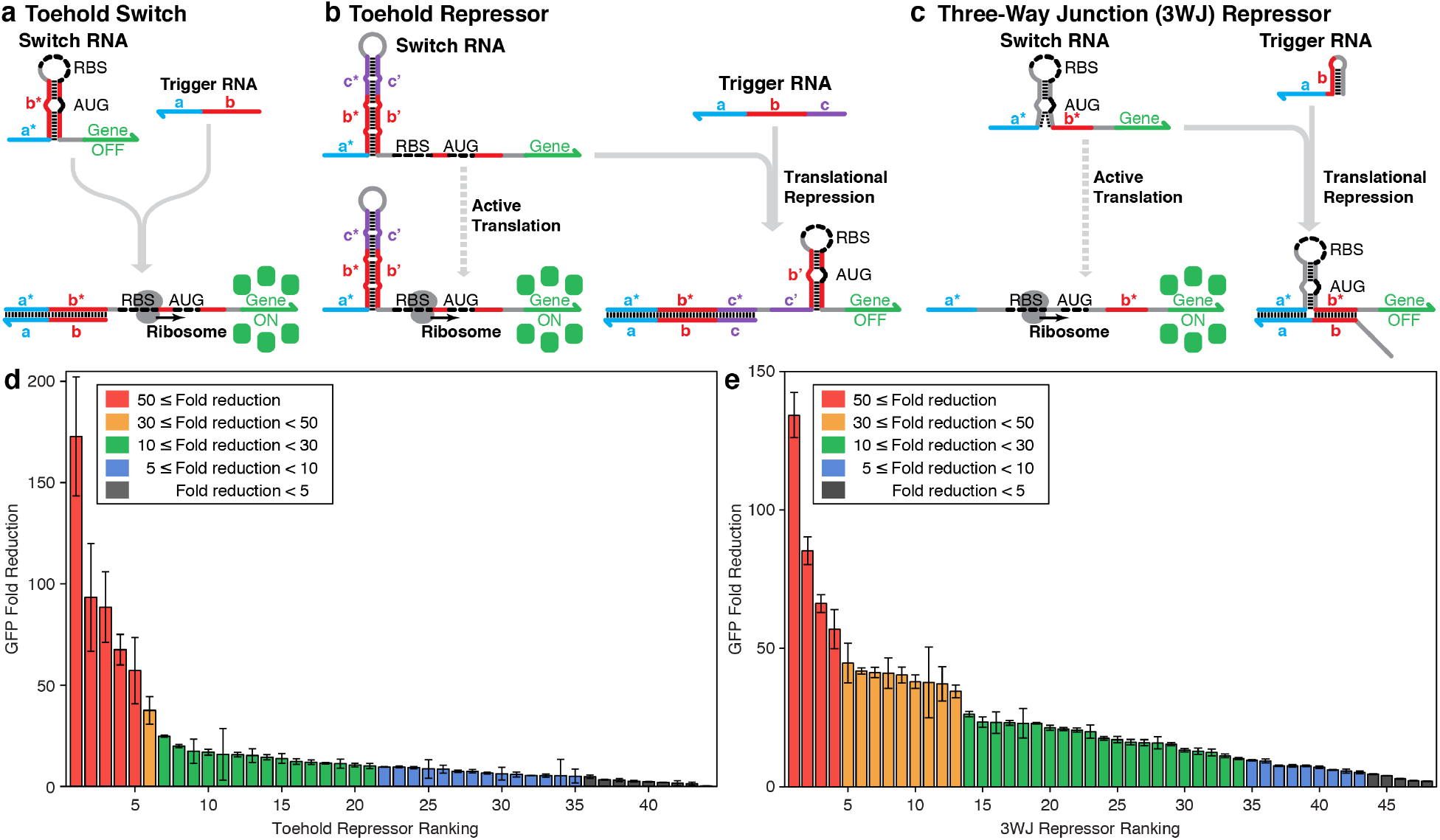
Operating mechanisms of *de*-*novo*-designed repressors and *in vivo* library characterization. **a**, Toehold switches repress translation through programmed base pairs before and after the start codon (AUG), leaving the ribosomal binding site (RBS) and start codon regions completely unpaired. RNA-RNA interactions are initiated via a single-stranded interaction domain called a toehold. The toehold domain **a\*** of switch RNA binds to a complementary **a** domain on the trigger RNA, initiating branch migration through domain **b** to open up the RBS and start codon of the switch RNA. **b**, Toehold repressors harbor a programmed strong hairpin structure upstream of an exposed RBS and start codon, allowing translation of a downstream gene. The toehold domain **a\*** of switch RNA binds to a complementary **a** domain on the trigger RNA, initiating branch migration through domains **b** and **c** to open the strong hairpin stem. The newly freed domains are used to form a downstream hairpin structure that represses translation by sequestering the RBS and start codon. **c**, The switch RNA of the three-way junction (3WJ) repressors contains an unstable hairpin structure that allows ribosomal access to the RBS and start codon and provides two single-stranded domains **a\*** and **b\*** for trigger binding. The trigger RNA has a hairpin structure and employs the toehold domain **a** to bind to the switch RNA whereupon branch migration through domain **b** forms a three-way junction structure that prevents ribosomal access to the RBS and start codon. **d**, Fold reduction of GFP fluorescence levels obtained 3 hr after induction for 44 first-generation toehold repressors. **e**, Fold reduction of GFP fluorescence levels obtained 3 hr after induction for a library of 48 3WJ repressors. Relative errors for the ON and OFF translation states are from three biological replicates. Relative errors for GFP fold reduction were obtained by adding the relative errors of the repressor ON and OFF state fluorescence measurements in quadrature.

We sought to obtain a library of programmable, wide dynamic range translational repressors analogous to the toehold switches and devised two types of repressors inspired by the design principles of these earlier riboregulators. The first type of repressor employs a switch RNA with a 5’ toehold domain and is thus referred to as a toehold repressor (Figure 1b, see Supplementary Information and Supplementary Figure S1a for details). The 15-nt toehold domain of the switch RNA is followed by an extended hairpin structure and a single-stranded expression region containing an RBS, start codon, and the coding sequence of the output gene. In the absence of the trigger RNA, the exposed RBS and start codon enable active translation of the output gene. The trigger RNA of the toehold repressors is a 45-nt single-stranded RNA sequence that is complementary to the toehold and stem of the switch RNA. After binding of the trigger to the switch RNA toehold, the ensuing branch migration process unwinds the hairpin stem and releases the domains **b’** and **c’**. Domain **b’** is complementary to the sequences upstream and downstream of the start codon, and thus forms a hairpin structure with these domains. This newly formed hairpin recapitulates the repressed structure of the toehold switch with the RBS sequestered within a 12-nt loop and the start codon concealed within the stem. As a result, translation is repressed upon trigger binding. The toehold repressor trigger sequence does not possess complementary bases either to the RBS or the start codon, which allows arbitrary choice of potential trigger sequences. If a trigger RNA sequence leads to in-frame stop codons in the expression region, bulges can be introduced or shifted in the **b’** domain of the switch RNA hairpin to compensate.

The second type of repressor adopts a three-way junction structure to suppress translation and is thus referred to as a three-way junction (3WJ) repressor (Figure 1c, see Supplementary Information and Supplementary Figure S1b for details). The switch RNA in these systems makes use of an unstable hairpin secondary structure that contains an RBS in the loop region and a start codon in the stem region. Despite its high secondary structure, this unstable hairpin was previously demonstrated to be translationally active in toehold switch mRNA sensors^35^. On either side of the unstable hairpin are two single-stranded domains **a\*** and **b\***. We hypothesized that transient formation of the bottom stem domain of the hairpin would co-localize these two domains to provide an effective binding site for a complementary trigger RNA. To take advantage of this design feature and improve repressor orthogonality, we designed cognate triggers where domain **b** is mostly contained in a hairpin secondary structure and a toehold domain **a** and part of domain **b** is located at the 3’ end. When the trigger RNA is expressed, the toehold initially binds to the **a\*** domain and part of the **b\*** domain of the switch RNA. The switch RNA **b\*** domain then completes a branch migration to unwind the trigger RNA stem. The resulting trigger-switch complex has a stable three-way junction structure that effectively sequesters the RBS and start codon within the loop and stem of the switch RNA, respectively, and strongly represses translation. Despite the use of a trigger with a hairpin structure to improve device orthogonality, the 3WJ repressors can also detect nearly arbitrary trigger RNAs provided that the trigger RNA sequence does not lead to an in-frame stop codon in domain **b\***.

### *In Silico* Design of Repressor Libraries and *In Vivo* Validation

We generated libraries of both translational repressors *de novo* using the NUPACK nucleic acid sequence design package^41^. A total of 44 toehold repressors and 48 3WJ repressors were designed and validated *in vivo* (see Supplementary Tables S1–S3 for sequence information). Members of the toehold repressor library were selected to reduce the potential for a non-cognate trigger RNA to disrupt the switch RNA stem. Members of the 3WJ repressor library were selected to minimize the potential for the non-cognate trigger RNAs to interact with the switch RNA. The *E. coli* BL21 Star DE3 strain with an IPTG-inducible genomic T7 RNA polymerase and decreased RNase activity was used for repressor characterization. A medium-copy plasmid containing the switch RNA regulating GFP and a high-copy plasmid encoding trigger RNA were transformed into *E. coli*. For measurements in the absence of a cognate trigger RNA, a non-cognate RNA strand with high secondary structure was transcribed from the high-copy plasmid.

Figure 1d shows the fold reduction of GFP fluorescence observed for the toehold repressor library after three hours of IPTG induction. The GFP fold reduction was measured from the geometric mean fluorescence of GFP obtained from flow cytometry for cells in the ON state expressing the non-cognate trigger RNA and in the repressed OFF state expressing a cognate trigger RNA (see Supplementary Figure S2a for ON and OFF state GFP expression levels). Cell autofluorescence was not subtracted from either the ON or OFF state fluorescence for determination of the GFP fold reduction. Although the toehold repressor devices show wide variations in performance, 48% or 21 out of 44 provide at least 10-fold change in gene expression upon detection of the trigger RNA. Furthermore, five devices or 11% exhibit GFP fold reduction of at least 50-fold, corresponding to over 98% repression of GFP signal. The 3WJ repressors overall provided improved performance compared to the toehold repressors (Figure 1e). A substantially higher fraction of these devices at 71% or 34 out of 48 provided at least 10-fold reduction of GFP expression, while a smaller fraction (8% or 4 out of 48) yielded exceptionally high 50-fold reduction in GFP (see Supplementary Figure S2b for ON and OFF state GFP expression levels). Although BL21 Star DE3 is an RNase-deficient strain, we found that the both types of repressors provided greater than 20-fold reduction of GFP in *E. coli* BL21 DE3 cells with wild-type RNase levels (see Supplementary Figure S3).

### Automated Forward Engineering of Second-Generation Toehold Repressors

To generate toehold repressors with higher performance, we implemented an automated strategy for ranking putative riboregulator devices (Figure 2a). We first compiled a set of 114 different thermodynamic parameters that could be computed rapidly from the sequence information of the trigger and switch RNAs (see Supplementary Information and Supplementary Figure S4 for details). This set of thermodynamic parameters and experimental GFP fluorescence data from the toehold repressor library were then used as inputs for the ranking algorithm. The algorithm first computed all three-parameter linear regressions between the thermodynamic parameters and the GFP fold reduction of each toehold repressor. Figure 2b displays the best three-parameter linear regression obtained from experimental measurements of GFP fold reduction, which provided a coefficient of determination R^2^ of 0.422. From the set of over 200,000 regressions, a scoring function was generated from the top ten regressions providing the highest R^2^ values.

**Figure 2.**
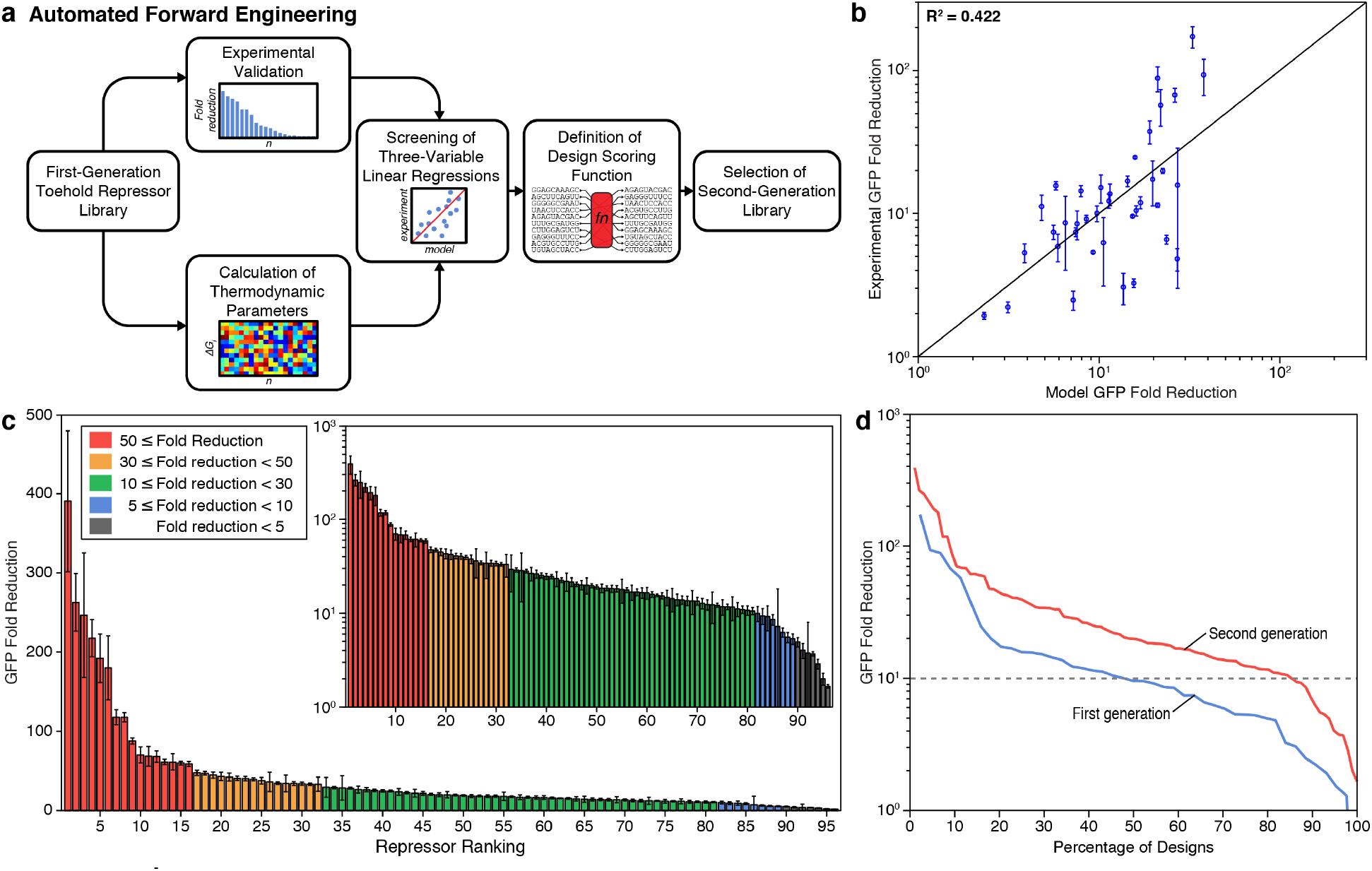
Thermodynamic analysis and automated forward engineering of toehold repressors. **a**, Automated forward engineering was carried out by screening three-variable linear regressions based on 114 different thermodynamic parameters. The top 10 three-variable regressions were used to generate a scoring function, which was then used to select second-generation toehold repressor designs. **b**, Correlation between predicted performance of toehold repressors using a three-parameter linear regression model and experimentally observed performance of the repressors. **c**, Fold reduction of GFP fluorescence levels obtained 3 hr after induction for 96 second-generation toehold repressors. Relative errors for ON and OFF translation states are from three biological replicates. Relative errors for GFP fold reduction were obtained by adding the relative errors of the repressor ON and OFF state fluorescence measurements in quadrature. **d**, Percentage of first-generation and second-generation library components that had GFP fold reduction that exceeded the value defined on y-axis. The GFP fold reduction of 10 is marked by gray dashed line.

NUPACK was used to generate an additional set of 265 toehold repressor sequences using identical secondary structures and design parameters as the first-generation library. The automatically generated scoring function was then used to rank each of the devices to select the top 96 for a second-generation library for validation (see Supplementary Table S4 for sequence information). Figure 2c presents the fold reduction of GFP fluorescence for the second-generation toehold repressors after 3 hours of induction (see Supplementary Figure S2c for ON and OFF state GFP expression levels). There is a dramatic increase in GFP fold reduction for the devices in general, with 8 switches exhibiting a dynamic range greater than 100 and 81 switches exhibiting a dynamic range greater than 10. The second-generation systems exhibit an average GFP fold reduction of 40 compared to 20 for the first-generation library. High-performing toehold repressors exhibit fold changes rivaling the dynamic range of protein-based regulators without requiring any *in vitro* evolution or large-scale screening experiments. We quantified the effectiveness of our selection criteria by calculating the percentage of toehold repressors with GFP fold reductions exceeding a given minimal level (Figure 2d). The yield of high-performance switches is higher for the second-generation devices across all fold reductions. For example, more than 84% of second-generation designs show at least 10-fold reduction as compared to 48% of first-generation designs.

### SHAPE-Seq Measurements of 3WJ Repressor Structure

To better understand the operating mechanism of the synthetic repressors, we performed in-cell SHAPE-Seq^42^ on several devices with varying repression efficiencies. The strong secondary structure of the toehold repressors, however, prevented interrogation by SHAPE-Seq. Fortunately, the weaker secondary structures of the 3WJ repressors enabled SHAPE-Seq studies for multiple trigger-switch interaction lengths. In the SHAPE-Seq experiment, 1-methyl-7-nitroisatoic anhydride (1M7) is introduced into the cell culture. 1M7 covalently modifies cellular RNAs in a structure-dependent manner, preferentially at positions that are unstructured and unconstrained by interactions. These modifications can be detected by reverse transcription stops coupled to high-throughput sequencing, and the mapped modification positions can then be used to calculate a reactivity value (b) at each nucleotide. Higher reactivities correspond to flexible or unstructured nucleotides, and lower reactivities indicate constrained interactions such as base-pairing or stacking effects. Simultaneous measurements of GFP expression using the same cell cultures used for structural probing allow direct links to be drawn between the performance of repressor variants and their structures.

We first studied a single 3WJ repressor switch RNA and three trigger variants, with interaction lengths ranging from 18 to 25 nts (Figure 3a, see Supplementary Table S5 for sequence information). Functional characterization demonstrated active translation from the switch RNA and strong translational repression upon trigger expression (Figure 3b). SHAPE-Seq reactivity measurements of these variants showed remarkable agreement with the proposed *in silico* design strategy. When the trigger RNA was not expressed, we observed a trend of high reactivities across the switch RNA sequence (Figure 3c). This reactivity signature supports the design hypothesis that the switch hairpin is sufficiently weak to facilitate structural disruption by ribosomes, leading to active translation. A striking difference is seen when a trigger RNA is expressed (Figure 3d). Sharp drops in reactivity are observed precisely at the predicted binding sites of each trigger (**a-a\*** in blue and **b-b\*** in red), providing structural evidence of trigger binding across the junction. Moreover, drops in reactivity also occur within the stem of the switch RNA hairpin at regions predicted to form the hairpin structure, providing direct evidence that trigger binding leads to the formation of a stable, translationally inaccessible 3WJ structure. Interestingly, higher reactivities are observed at several positions around the base of the hairpin when the triggers are present (specifically U16-U19), suggesting slight fraying or flexibility at the base of the trigger-switch three-way junction. We also studied a second 3WJ repressor with different triggers. These experiments also showed formation of the intended three-way junction structure in the repressed state (see Supplementary Information for details and Supplementary Figure S5). To the best of our knowledge, these results represent the first structural confirmation of the regulatory mechanism of a completely *de-novo*-designed riboregulator.

**Figure 3.**
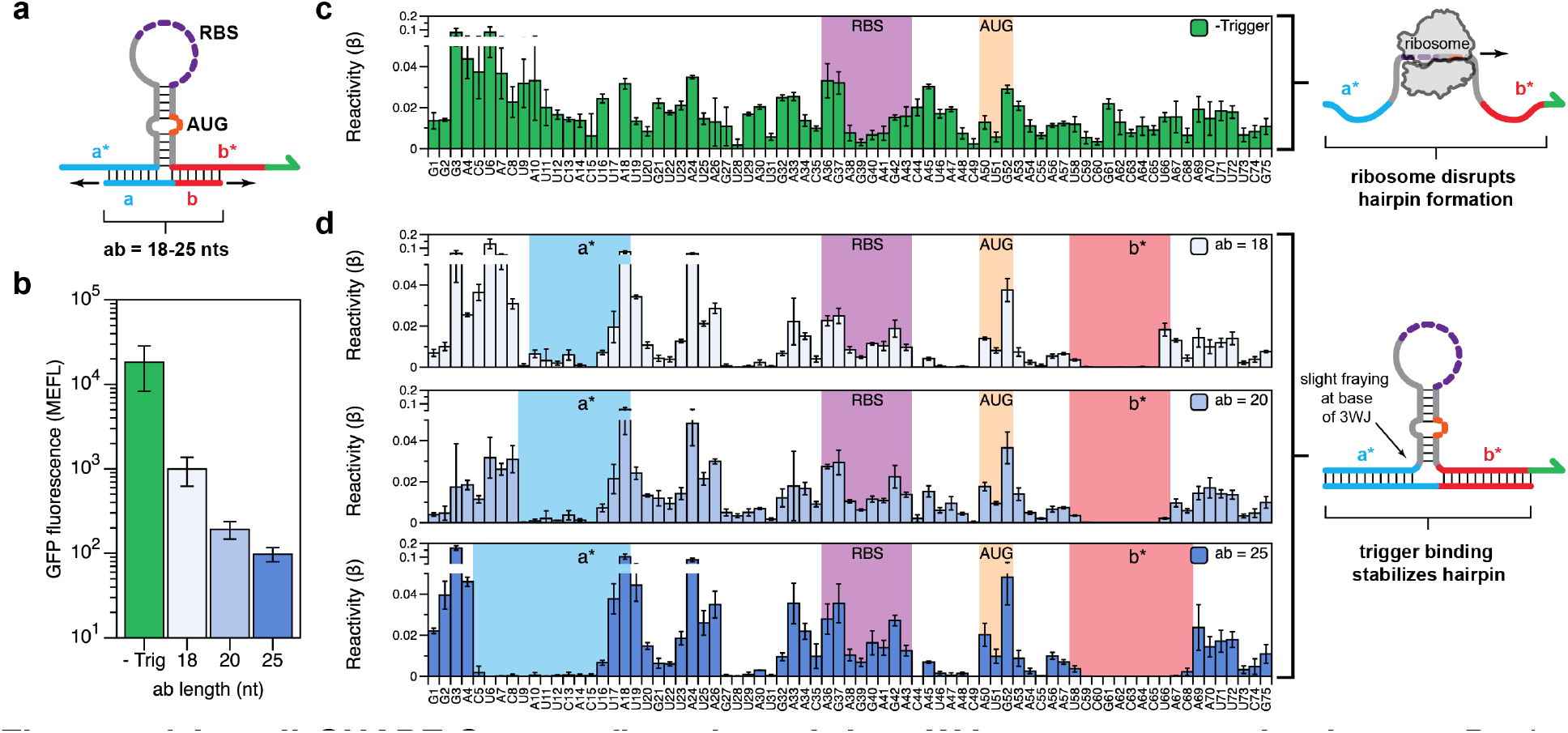
In-cell SHAPE-Seq confirmation of the 3WJ repressor mechanism. **a**, Design schematic for testing 3WJ repressor variants. A 3WJ repressor switch was characterized using in-cell SHAPE-Seq, either expressed alone or co-expressed with a trigger RNA. Several triggers were tested, varying in their designed binding length (**ab**) to either side of the switch hairpin. **b**, Functional characterization of switch plasmid expressed without trigger (green) and with triggers of increasing interaction length (blue). Strong repression is observed upon trigger binding, with longer triggers showing increased repression efficiency. **c**, In-cell SHAPE-Seq reactivity profile of the switch RNA expressed alone. A trend of high reactivities is observed across the molecule, consistent with the design hypothesis that the switch hairpin can be disrupted by ribosome binding, leading to active translation. **d**, In-cell SHAPE-Seq reactivity profiles of the switch co-expressed with trigger RNAs. Sharp drops in reactivity are observed at the predicted trigger binding sites (**a-a\*** and **b-b\***) and within the switch hairpin, suggesting formation of a stable 3WJ structure when the trigger is bound. The RBS and start codon (AUG) positions are indicated.

### Evaluation of Repressor Orthogonality

One of the prerequisites for higher-order logic processing is the orthogonality of regulatory components with respect to one another. We thus measured *in vivo* the interactions between pairwise combinations of different repressor trigger and switch RNAs. For the second-generation toehold repressors, we first performed *in silico* screening to isolate a subset of 16 devices that provided more than 10-fold GFP reduction and also displayed low levels of predicted crosstalk with non-cognate triggers. Flow cytometry was used to quantify GFP output in *E. coli* for all 256 trigger-switch interactions after three hours of IPTG induction (Figure 4a). Crosstalk in terms of GFP fold reduction was calculated by dividing the GFP fluorescence obtained from a non-cognate trigger and a given switch RNA by the fluorescence of the switch in its triggered state. Typically, non-cognate trigger-switch pairs showed significantly higher GFP output as compared to cognate trigger-switch pairs as shown in Figure 4a. For instance, the first column shows that the GFP expression for toehold repressor switch 3 was at least 4-fold higher for all non-cognate RNAs compared to its cognate trigger RNA 3. However, the crosstalk level was high in many instances such that the set of orthogonal devices that maintained at least 12-fold dynamic range was reduced to four devices as indicated by the trigger-switch combinations in the red boxes in Figure 4a. For the less stringent orthogonality condition of at least 7-fold dynamic range, the toehold repressor library yielded a set of eight independent riboregulators.

**Figure 4.**
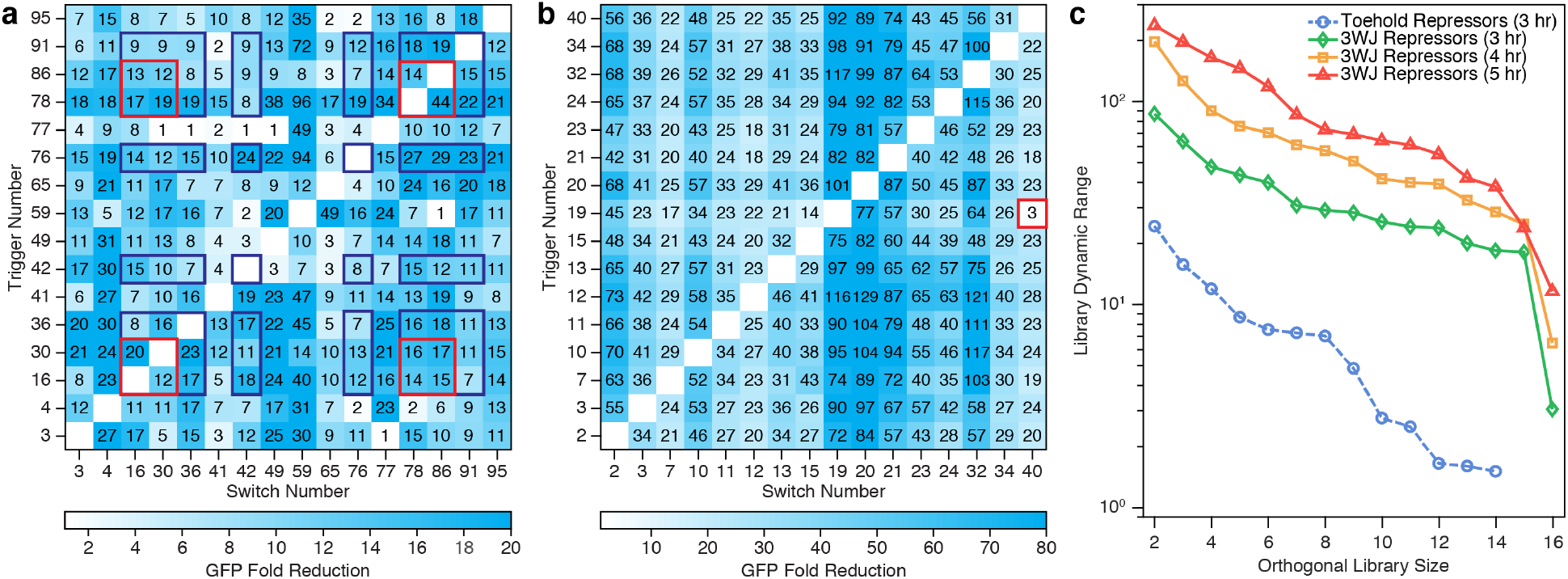
Assessment of toehold and 3WJ repressor orthogonality. **a**, Toehold repressor crosstalk measured by flow cytometry for 256 trigger-switch combinations 3 hours after induction. Red boxes designate a subset of four repressors that exhibit sufficiently low crosstalk to provide at least 12-fold GFP reduction. Blue boxes designate a subset of eight repressors that provide at least 7-fold GFP reduction. **b**, Three-way junction repressor crosstalk measured by flow cytometry for 256 trigger-switch combinations 3 hours after induction. All but one of the 240 non-cognate combinations provides at least 14-fold higher GFP expression than cognate pairs. **c**, Comparison of overall library dynamic range and orthogonal library size for the toehold repressors and 3WJ repressors. Three-way junction repressor orthogonal library size and dynamic range increase over the 3- to 5-hour IPTG induction time.

Based on the shorter exposed single-stranded regions of the 3WJ repressor trigger RNAs, which reduced the potential for interaction with switch RNAs, we anticipated that the 3WJ repressor devices would show improved orthogonality compared to the toehold repressors. We measured the pairwise trigger-switch interactions for 16 of the top devices using the same methods applied to the toehold repressors (Figure 4b). The 3WJ repressor library showed substantially reduced crosstalk while maintaining strong repression of cognate trigger-switch pairs (see Supplementary Figure S6 for GFP expression levels). In fact, we found that a set of 15 out of the 16 3WJ repressors tested provided at least 17-fold reductions in GFP expression in the presence of the cognate trigger compared to any of the other 14 noncognate triggers in the orthogonal set. Moreover, we only observed significant crosstalk in a single pairwise interaction, which occurred between switch 40 and trigger 19 (red box in Figure 4b).

To quantify the orthogonality of the devices, we determined the maximum number of repressors that could be used to provide a given minimum level of overall dynamic range (Figure 4c, see Supplementary Table S6 for the sets of orthogonal repressors). This analysis showed large improvements in orthogonal library size and dynamic range for the 3WJ repressors compared to the toehold repressors. For example, the most orthogonal eight-device toehold repressor set provided an overall dynamic range of at least 7-fold, while the corresponding eight-device 3WJ repressor set yielded an overall dynamic range of 29-fold. We also applied this analysis to measurements of the 3WJ repressors induced over longer four-hour and five-hour induction times. As the induction time increased, the overall fold reduction of GFP in the cells generally increased, leading to parallel improvements in device orthogonality. For instance, an eight-device orthogonal 3WJ repressor library provided a dynamic range that rose to 73-fold after five hours of induction. Moreover, a set of six 3WJ repressors provided a remarkable library dynamic range of 118 at the five-hour time point.

### mRNA-Sensing Repressors

The ability to detect nearly arbitrary trigger RNA molecules enables synthetic repressors to respond to intracellular mRNAs and has potential diagnostic applications^34–37^. We implemented mRNA-sensing toehold repressors by extending their toehold domains to 30 nts to compensate for the increased secondary structure of target mRNAs. *In silico* screening was then used to identify fragments along the target mRNA that provided the lowest secondary structure to facilitate repressor binding (Figure 5a, Supplementary Table S7). For the 3WJ repressors, we left the riboregulator design unchanged and selected target mRNA binding sites by determining 27-nt regions having low secondary structure.

**Figure 5.**
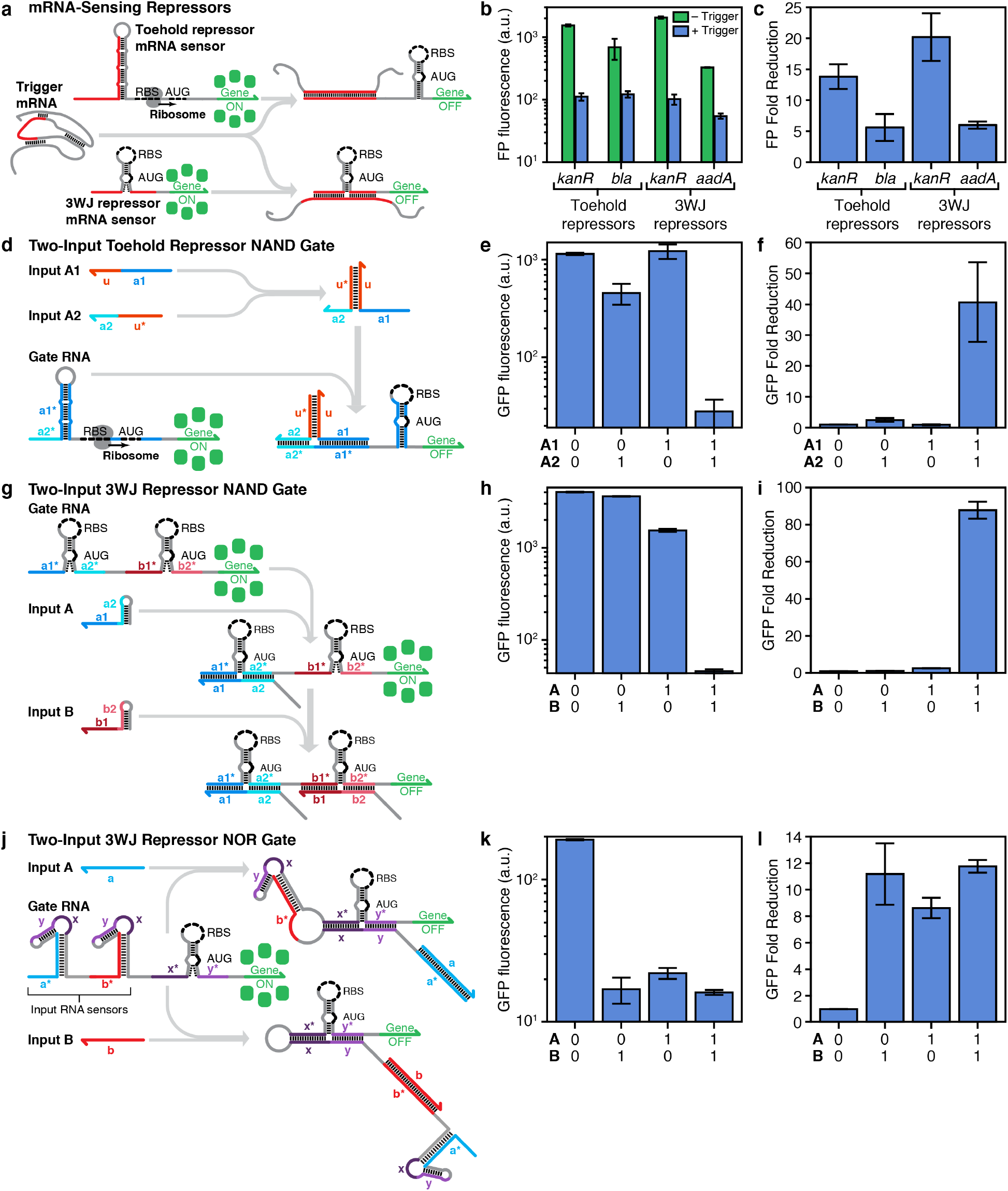
mRNA sensing and two-input logic operations using repressor-based devices. **a**, Design schematic of the mRNA-sensing toehold and 3WJ repressors. The region within the mRNA sequence used to trigger repression is emphasized in red. **b**, Fluorescent protein (FP) fluorescence observed for toehold and 3WJ repressors targeting two different pairs of antibiotic resistance mRNAs. **c**, Fold reduction of FP for the toehold and 3WJ repressor mRNA sensors. **d**, Design schematic of a two-input NAND gate based on toehold repressors. Two input RNAs hybridize through domains **u** and **u\*** to form a complete trigger sequence to repress the gate RNA. **e, f**, GFP fluorescence (e) and fold reduction (f) measurements of multiple input combinations for the two-input NAND gate based on toehold repressors. **g**, Design schematic of a two-input NAND gate constructed from 3WJ repressors. In the gate RNA, two switch hairpin modules are inserted in-frame and upstream of the reporter gene. Both input RNAs are required to bind to the gate RNA to prevent gene expression. **h, i**, GFP fluorescence (h) and fold reduction (i) measurements of multiple input combinations for the two-input NAND gate based on 3WJ repressors. **j**, Design schematic of a two-input NOR gate based on 3WJ repressors. The gate RNA contains a single 3WJ repressor hairpin to regulate the output gene and a pair of trigger modules sequestered within the loops of strong hairpin secondary structures that sense the cognate input RNAs. Toehold-mediated binding of either input RNA causes the corresponding hairpin to unwind, which releases the trigger to bind to the 3WJ repressor module and inhibit gene expression. **k, l**, GFP fluorescence (k) and fold reduction (l) measurements of multiple input combinations for the two-input NOR gate based on 3WJ repressors. The FP fold reductions of devices are calculated by dividing the fluorescence from the gate RNA obtained for the null input case by the fluorescence for each input combination. Relative errors for different states are from the SD of three biological replicates. Relative errors for the mRNA sensors and ribocomputing device fold reductions were obtained by adding the relative fluorescence errors in quadrature.

The mRNA-sensing repressors were validated against several mRNAs encoding antibiotic resistance genes: the kanamycin resistance protein (*kanR)*, beta-lactamase (*bla*) conferring resistance to the antibiotic ampicillin, and *aadA* conferring resistance to spectinomycin. For the toehold repressors, we constructed a *bla* sensor to repress the translation of GFP output after binding to the cognate *bla* mRNA transcripts and a *kanR* sensor to repress mCherry output upon binding to *kanR* mRNA. For the 3WJ repressors, we constructed *kanR* and *aadA* sensors that both regulated GFP output. The mRNA-sensing repressors were then tested using procedures employed for library validation and expression induced for 5 hours using IPTG. Figure 5b displays the fluorescence intensity of the fluorescent proteins (FPs) GFP and mCherry produced by the repressors with and without expression of the target mRNAs. For all four mRNA sensors, protein expression is significantly reduced in the presence of the cognate mRNA. Mean fluorescence intensities were then used to determine the fold reduction of each FP for the mRNA sensors (Figure 5c). Both the toehold and 3WJ repressors provided greater than 10-fold reductions in mCherry and GFP fluorescence, respectively, upon inhibition by the *kanR* mRNA. Sensors for *bla* and *aadA* yielded 5-fold reductions in GFP expression in response to these mRNAs. These results show that the programmable repressors can be used to strongly inhibit output protein expression in response to functional mRNAs.

### Repressor-Based Logic Circuitry

The modular and programmable nature of the toehold and 3WJ repressors makes them ideal candidates for integration into ribocomputing devices for implementing sophisticated genetic programs. We have previously demonstrated that toehold switch riboregulators can be incorporated into such RNA-based computing systems for multi-input intracellular computation using RNA input signals and protein signals^33,43^. These devices employ RNA-RNA interactions for all signal processing tasks and they co-localize signal processing functions by using gate RNA transcripts containing one or more riboregulator modules upstream of the output gene. We thus applied the ribocomputing strategy to the repressors to enable efficient computation of NAND and NOR logic functions in living cells.

We first studied two-input NAND gates based on toehold repressors optimized for these logic operations (see Supplementary Information for circuit design details and Supplementary Table S8 for circuit sequence information). The trigger RNA sequence was divided into two input RNAs A1 and A2 and complementary bridging domains **u** and **u\*** were appended to each input (Figure 5d and Supplementary Figure S7a). Since neither input has the complete trigger sequence, they are unable to activate the toehold repressor gate, which consists of a single switch RNA hairpin module upstream of a GFP output gene. However, when both inputs A1 and A2 are present, they hybridize to one another through the **u**-**u\*** interaction and bring both trigger halves into close proximity for binding to the gate RNA. Similar associative toehold mechanisms have been demonstrated *in vitro*^44,45^ and have been used for AND logic in ribocomputing devices^33^. Figure 5e shows the mean GFP fluorescence intensity obtained from the two-input NAND circuit after 6 hours of induction by IPTG (see Supplementary Figure S8 for GFP population histograms of all ribocomputing circuits). The RNA inputs and the gate RNA were expressed from separate plasmids through the T7 promoter in BL21 Star DE3 cells. In cases where an input RNA was not present, a non-cognate RNA was expressed in its place. For the logical TRUE input conditions with neither input or only one input present, GFP output from the gate RNA remains high. However, when the logical FALSE condition occurs with two input RNAs expressed, the NAND gate provides strongly reduced GFP expression. Mean GFP fluorescence levels obtained for the null input condition with no cognate input RNAs expressed were divided by the mean GFP fluorescence obtained from each of the input conditions to compute the fold reductions for the circuit (Figure 5f). This analysis shows that GFP levels were reduced by 40-fold in the logical FALSE state. We also observed a noticeable decrease of 2.5-fold upon expression of input A2 alone. We attribute this leakage effect to the input binding causing a partial disruption of the gate RNA stem and potentially to cross interactions between the **u\*** bridging domain with exposed single-stranded regions of the gate.

We also implemented repressor-based gate RNAs integrating multiple repressor hairpin modules upstream of the output gene. Attempts using toehold repressor hairpins, however, proved unsuccessful since their strong hairpin secondary structure and long target RNA binding sites prevented efficient translation of the downstream gene. In contrast, the comparatively weak secondary structure of the 3WJ repressors and their short trigger RNA binding sites were ideal for incorporation into gate RNAs. We implemented a two-input NAND gate RNA composed of two orthogonal 3WJ repressor hairpins separated by a 17-nt single-stranded spacer domain (Figure 5g). In the presence of only one input RNA, the input can bind to repress translation from its corresponding hairpin module. However, translation of the output gene will continue from the unrepressed hairpin module, since the ribosome can translate through weak hairpin secondary structures and duplexes formed by input and gate RNAs. As a result, only upon binding of both input RNAs at the same time to the gate RNA will output gene expression be fully inhibited. We evaluated the resulting two-input NAND gate and found that GFP expression remained strong except for the logical FALSE case with both inputs expressed after 6 hours of induction by IPTG (Figure 5h). Small decreases in GFP expression were observed when only one of the input RNAs was present, likely as a result of inhibition of one of the two translation initiation sites from the gate RNA. The GFP fold reductions of the circuit show a large 88-fold decrease in expression in response to the two input RNAs compared to null input case (Figure 5i). This reduction was at least a factor of 33 higher than any of the changes in expression observed for single-input cases.

To implement NOR ribocomputing devices responsive to two sequence-independent input RNAs, we developed a new gate RNA architecture that exploited co-localized intramolecular interactions (Figure 5j and Supplementary Figure S9a). These NOR gate RNAs contain multiple sequestered trigger RNA sequences upstream of a 3WJ repressor module regulating the output gene. The trigger RNA domains **x** and **y**, which are complementary to the downstream repressor hairpin, are confined within the loops of strong hairpin secondary structures. These hairpins function as input RNA sensors that provide toehold domains for binding to complementary input RNA sequences. When an input RNA is expressed, binding to the input sensor leads to a branch migration that unwinds the sensor stem. This interaction releases the trigger RNA domain and enables the trigger to repress the downstream 3WJ repressor domain through an efficient gate RNA intramolecular interaction. We constructed the two-input NOR gate RNA using a validated 3WJ repressor and two input sensor hairpins, resulting in a gate RNA regulatory region of 312 nts. Measurements of GFP fluorescence from this gate RNA showed a substantial reduction in fluorescence upon expression of any of the cognate input RNAs after 6 hours of IPTG induction (Figure 5k). We observed lower GFP output from the NOR gate RNA for the null input case compared to other 3WJ repressor gate RNAs. It is likely that there is some repression leakage from incomplete sequestration of the trigger RNA domains. Despite the lower overall expression level of the gate RNA, analysis of the GFP fold reductions from the circuit show between 8- to 12-fold decrease in GFP output in response to one or two input RNAs (Figure 5l) and confirm the successful operation of the new gate RNA design.

### Three- and Four-Input Repressor-Based Logic Circuitry

To further evaluate the capabilities of the repressor-based ribocomputing devices, we implemented several multi-input logic circuits employing more than two input RNAs. NAND gate RNAs based on 3WJ repressors were extended to three- and four-input operation by adding additional repressor modules upstream of the output gene. A three-input gate RNA for NOT (A AND B AND C) computations was constructed using three orthogonal 3WJ repressor hairpins separated by 11-nt single-stranded spacer domains (Figure 6a). This ribocomputing device showed high GFP expression for all logical TRUE conditions lacking at least one of the input RNAs (Figure 6b), while providing low expression for the logical FALSE condition with all inputs. The GFP fold reduction for this NAND circuit was 163-fold over the null input case and provided at least 33-fold lower GFP expression than the other input RNA combinations (Figure 6c). We constructed a four-input device to evaluate the expression NOT (A AND B AND C AND D) using a gate RNA with four orthogonal 3WJ repressor modules separated by 17-nt single-stranded spacers, resulting in a gate RNA regulatory region of 365 nts (Figure 6d and Supplementary Figure S9b). The GFP fluorescence for this device remained high for all logical TRUE conditions and decreased substantially when all input RNAs were expressed (Figure 6e). Calculation of the GFP fold changes for this circuit showed a 6-fold reduction in GFP expression in the sole logical FALSE state (Figure 6f). The overall dynamic range of the four-input NAND gate was substantially lower than that of the three-input NAND gate. We attribute the weaker performance of this circuit to the difficulty in forming the complete five-RNA molecular complex required for complete repression of gene expression. Nevertheless, the four-input NAND ribocomputing device provided at least a 3.7-fold reduction in GFP compared to all logical TRUE states.

**Figure 6.**
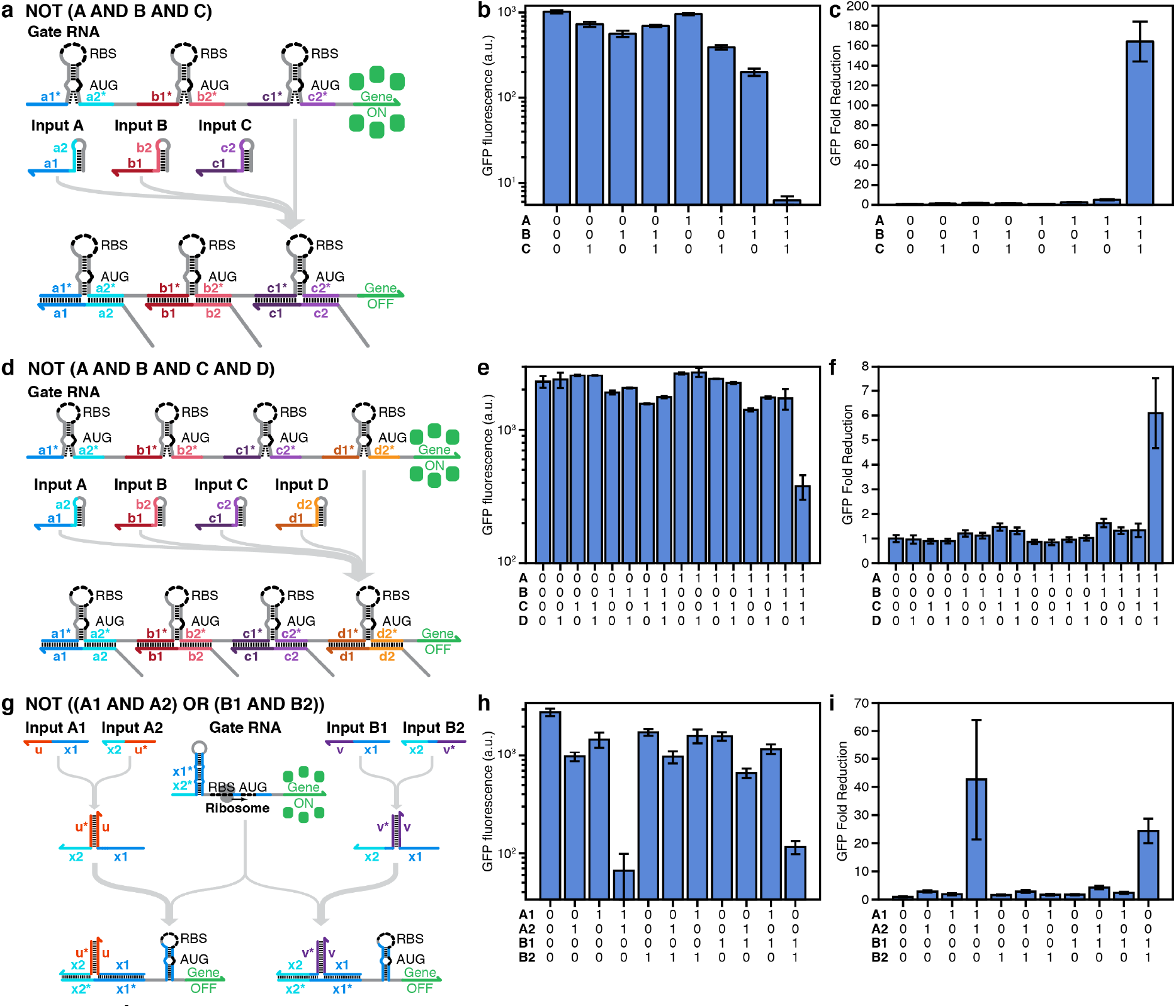
Multi-input ribocomputing devices employing toehold and 3WJ repressors. **a**, Design schematic for a three-input NAND gate based on 3WJ repressors. The gate RNA contains three orthogonal 3WJ repressor hairpins in-frame and upstream of the output gene. **b, c**, GFP fluorescence (b) and fold reduction (c) measurements of multiple input combinations for the three-input NAND device based on 3WJ repressors. **d**, Design schematic for a four-input NAND gate based on 3WJ repressors. The gate RNA contains four orthogonal 3WJ repressor hairpins in-frame and upstream of the output gene. **e, f**, GFP fluorescence (e) and fold reduction (f) measurements of multiple input combinations for the four-input NAND device based on 3WJ repressors. **g**, Design schematic for a NOT ((A1 AND A2) OR (B1 AND B2)) logic circuit. Independent two-input NAND gate behavior is enabled by two partial-trigger domains **x1** and **x2** coupled with bridging domains **u** and **u\*** for inputs A1 and A2 or with orthogonal bridging domains **v** and **v\*** and a shifted junction site for inputs B1 and B2. The two partial-triggers together present the full-length trigger for the toehold repressor. **h, i**, GFP fluorescence (h) and fold reduction (i) measurements of multiple input combinations for the four-input ribocomputing device based on toehold repressors. The GFP fold reductions of devices are calculated by dividing the GFP fluorescence from gate RNA obtained for the null input case by the GFP fluorescence for given input combinations. Relative errors for ON and OFF states are from the SD of three biological replicates. Relative errors for the ribocomputing device fold reductions were obtained by adding the relative errors of the sensor ON and OFF state fluorescence measurements in quadrature.

A second four-input logic system was implemented based on the toehold repressor NAND system (Figure 5d-f). Observing the domains in inputs A1 and A2, we recognized that there are no sequence constraints in choosing the bridge sequences appended to the two partial-trigger domains. We thus designed a second set of bridge domains **v** and **v\*** and shifted the trigger splitting point by 4 nts to generate two new NAND inputs B1 and B2 (Figure 6g and Supplementary Figure S7b). The resulting ribocomputing device, which performed the computation NOT ((A1 AND A2) OR (B1 AND B2)), was tested using multiple combinations of the four input RNAs (Figure 6h-i). As expected, we observed substantial reductions in GFP expression only when A1 and A2 or B1 and B2 were expressed simultaneously. Further, only weak crosstalk was observed when the non-interacting A triggers and B triggers were tested in pairs. The crosstalk observed for the trigger A2 and B1 combination was at least 5-fold less than the cognate pair of triggers.

## DISCUSSION

We have developed two new types of high-performance translational repressors using *de novo* RNA sequence design. Toehold repressors, optimized using an automated forward-engineering procedure, exploit strong RNA secondary structures to provide a very wide dynamic range of gene expression and are thus best suited for applications requiring tight translational control. Three-way junction repressors exhibit very low device crosstalk while using weaker RNA secondary structures and are optimal for multiplexed sensing and multi-input logic systems. In-cell SHAPE-Seq measurements of the 3WJ repressor system provided for the first time direct structural confirmation of the operating mechanism of a *de*-*novo-* designed riboregulator. We harnessed the modular and programmable nature of the synthetic repressors to detect full-length mRNA molecules and to integrate them into ribocomputing device architectures for evaluating multi-input logic expressions in living cells.

Compared to previous translational repressors, toehold and 3WJ repressors exhibited substantially improved dynamic range, with both libraries having multiple devices with reductions in gene expression well in excess of 50-fold. Forward engineering further resulted in toehold repressors with up to 300-fold reduction in GFP expression. Previously reported translational repressors based on the IS10 system successfully reduced GFP expression by ~10-fold and formed an orthogonal library of six devices with under 20% crosstalk^29^. This library size is similar to the orthogonal library of eight toehold repressors providing 7-fold dynamic range. In contrast, the 3WJ repressors, which featured designs better optimized to reduce crosstalk, provided substantially improved orthogonality with 15 repressors yielding 17-fold dynamic range.

Like toehold switches that activate gene expression, toehold and 3WJ repressors can detect trigger RNAs with nearly arbitrary sequences and exploit favorable linear-linear interactions for trigger-switch binding. Despite these design similarities, there are several noteworthy performance differences between the three types of translational riboregulators. First, we found that both repressor systems provided lower dynamic range than the toehold switch activators, which can often provide over 400-fold increases in GFP expression after forward engineering^32^ or design tuning^33^. These differences in dynamic range are primarily caused by translation from the switch RNA that occurs prior to binding of the trigger RNA to repress translation. Toehold activators, whose switch RNAs are initially transcribed in a repressed state, are not subject to this form of leakage. Second, despite the large available sequence space, we found that the toehold repressors provided substantially smaller orthogonal libraries compared to toehold switch activators and 3WJ repressors. The toehold activators and 3WJ repressors both yielded libraries of over a dozen devices providing dynamic range over 20-fold^32^, while toehold repressors were limited to sets of two or three devices with similar overall dynamic range. We attribute this reduced orthogonality to the extended single-stranded regions of the toehold repressor switch and trigger RNAs. Although non-cognate switch and trigger RNAs were selected to reduce the potential for hybridization, their *de*-*novo*-designed single-stranded regions, with lengths of 30 and 45 nts, respectively, make it very challenging to exclude all non-specific binding interactions, which can occur across small domains of complementarity. In particular, binding of non-cognate trigger molecules to portions of the exposed RBS and start codon regions in the switch RNA likely led to significant reductions in translational activity. In contrast, the 3WJ repressors partially conceal the RBS and start codon within a weak stem, which allows ribosomal access yet discourages interactions in *trans* with non-cognate trigger RNAs. For sensing mRNAs, non-specific interactions with the toehold repressors are not as deleterious since the increased secondary structure of natural RNAs discourages interactions in *trans* with switch molecules. Third, the 3WJ repressors showed weaker repression strength than the forward-engineered second-generation toehold repressors. Finally, while this work was being performed, an independent study uncovered similar designs for RNA-based translational repressors (Carlson et al., submitted). These designs enabled targeting of endogenous mRNA transcripts, but exhibited weaker repression efficiency than the systems presented here. Thus, depending on system requirements, one can choose a device architecture optimized for stronger repression, improved orthogonality, or silencing of targeted endogenous mRNAs.

The high performance and modularity of the toehold and 3WJ repressors enabled them to be incorporated into genetically compact ribocomputing devices that effectively computed NOT-related logic expressions with up to four different input RNAs. Previous implementations of ribocomputing NOT logic have been limited by sequence constraints due to a reliance on antisense RNA interactions. Using a split trigger design where co-localization of two partial triggers is required to deactivate the toehold repressor, we demonstrated a functional two-input NAND gate and extended this concept to obtain combined four-input NAND/NOR functionality. These trigger design strategies require partial sequence complementarity between the input RNAs used. Fortunately, the 3WJ repressor designs were amenable to integration into long gate RNAs that simultaneously detect multiple sequence-independent trigger RNAs. The gate RNA with multiple 3WJ repressors showed good performance in a NAND circuit using up to four input RNAs. Alternatively, NOR gates can be constructed by regulating a 3WJ switch module using trigger RNAs sequestered within upstream hairpin domains. Since translation of the NOR gate output protein begins at the single 3WJ switch module, this gate RNA architecture does not require translation through downstream hairpins nor does it append long N-terminal sequences to the output protein, which both occur for previously reported OR gate RNAs. We thus expect that this NOR gate design strategy can also be employed using toehold activators to improve the performance of OR gate ribocomputing systems.

We also successfully applied in-cell SHAPE-Seq^42^, which combines in-cell probing of RNA structure with gene expression measurements, to simultaneously characterize RNA structure and function in high-throughput for the 3WJ repressors. Analysis of 3WJ repressors yielded direct evidence to support our mechanistic model and also revealed potential pit-falls in our design strategies. These results highlight how SHAPE-Seq can be used to confirm design principles and understand potential failure modes of synthetic riboregulators. Moreover, the ability of RNA structure probing methods to detect interactions such as long-range tertiary structures and RNA-protein binding^46^ will make them even more important as the complexity of *de-novo*-designed riboregulators continues to increase. Indeed, we expect that future application of SHAPE-Seq to ribocomputing gate RNA designs will help uncover relationships between sequence, structure, and function. Such efforts, in concert with forward engineering strategies driven by thermodynamic models, RNA sequence design, and automated design methodologies^47^, have the potential to provide sizeable improvements in *de-novo*-designed riboregulator performance and RNA-based computing architectures.

Overall, the toehold and 3WJ repressors represent a versatile new set of components to add to the rapidly expanding RNA synthetic biology toolkit. The development of these NOT, NAND, and NOR logic devices coupled with advances in RNA-guided CRISPR/Cas systems^48,49^, RNA-based transcriptional regulators^27,50,51^, and systems that merge these capabilities^52^ point to increasingly sophisticated forms of RNA-enabled genetic circuits that exploit regulation at the transcriptional, translational, and post-transcriptional levels to achieve more dynamic and programmable cellular functions. We further expect that the ability of toehold and 3WJ repressors to detect nearly arbitrary RNAs will prove useful in cell-free diagnostic systems^34–37^ for identification of specific pathogen nucleic acids and their use of universal RNA-RNA interactions could enable their application in other prokaryotic hosts.

## METHODS

### Strains and growth conditions

These *E. coli* strains were used in this study: BL21 Star DE3 (F^-^ *ompT hsdS*_B_ (r_B_^-^ m_B_^-^) *gal dcm rne131* (DE3); Invitrogen), BL21 DE3 (F^-^ *ompT hsdS*_B_ (r_B_^-^ m_B_^-^) *gal dcm* (DE3); Invitrogen), and DH5α (*endA1 recA1 gyrA96 thi-1 glnV44 relA1 hsdR17* (r_K_^-^ m_K_^+^) λ^-^). All strains were grown in LB medium at 37°C with appropriate antibiotics: ampicillin (50 μg mL^-1^), spectinomycin (25 μg mL^-1^), and kanamycin (30 μg mL^-1^).

### Toehold and 3WJ repressor library construction

Plasmids were constructed using PCR and Gibson assembly. DNA templates for toehold repressor switch and trigger RNA expression were assembled from single-stranded DNAs purchased from Integrated DNA Technologies. The synthetic DNA strands were amplified via PCR and then inserted into plasmid backbones using 30-bp homology domains via Gibson assembly^53^. All plasmids were cloned in the *E. coli* DH5α strain and validated through DNA sequencing. Backbones for the plasmids were taken from the commercial vectors pET15b (ampicillin resistance, ColE1 origin), pCOLADuet (kanamycin resistance, ColA origin), and pCDFDuet (spectinomycin resistance, CDF origin) from EMD Millipore. GFPmut3b-ASV, GFPmut3b with an ASV degradation tag^54^, was used as the reporter for the toehold repressor switch plasmids except for the *kanR* mRNA sensor that used mCherry as the reporter. Sequences of elements commonly used in the plasmids are provided in Supplementary Tables.

### Toehold and 3WJ repressor expression

Toehold and 3WJ repressor switch and trigger RNAs were expressed using T7 RNA polymerase in BL21 Star DE3, an RNase-deficient strain, with the T7 RNA polymerase induced with the addition of isopropyl β–D-1-thiogalactopyranoside (IPTG). Selected sets of toehold and 3WJ repressor switch and trigger RNAs were also tested in BL21 DE3 strain with the T7 RNA polymerase induced with the addition of IPTG. For both strains, cells were grown overnight in 96-well plates with shaking at 900 rpm and 37°C. Overnight cultures were then diluted by 100-fold into fresh LB media with antibiotics and returned to shaker (900 rpm, 37C). After 80 minutes, both strains were induced with 0.1 mM IPTG and cells were returned to shaker (900 rpm, 37°C) until the flow cytometry measurements at specified times post-induction.

### Flow cytometry measurements and analysis

Flow cytometry measurements of toehold repressor libraries and ribocomputing devices were performed using a BD LSRFortessa cell analyzer with a high-throughput sampler. Prior to loading to flow cytometer, cells were diluted by a factor of ~65 into phosphate-buffered saline. Cells were detected using a forward scatter (FSC) trigger and at least 30,000 cells were recorded for each measurement. Flow cytometry measurements of 3WJ repressor libraries and ribocomputing devices were performed using a Stratedigm S1300EXi cell analyzer equipped with a A600 high-throughput autosampler. Cells with the 3WJ repressor systems were diluted by a factor of ~17 into phosphate-buffered saline and detected as described above with 40,000 cells recorded for each measurement. Cell populations were gated according to their FSC and side scatter (SSC) distributions as described previously^32^, and the GFP or mCherry fluorescence levels of these gated cells were used to measure circuit output. GFP or mCherry fluorescence histograms yielded unimodal population distributions and the geometric mean was employed to extract the average fluorescence across the approximately log-normal fluorescence distribution from at least three biological replicates. Fold reductions of GFP or mCherry fluorescence levels were then evaluated by taking the average fluorescence output of toehold or 3WJ repressor switch with a non-cognate trigger and dividing it by the fluorescence output with a cognate trigger. Cellular autofluorescence was not subtracted prior to determining the fold reduction and percent repression.

### SHAPE-Seq measurements and analysis

In-cell SHAPE-Seq measurements were carried out as described by Watters *et al*.^42^. Briefly, 3WJ repressor variants were transformed into BL21 Star DE3 as in the functional characterization experiments. Overnight cultures were diluted by a factor of 100 into 1.2 mL of fresh LB with antibiotics. Following IPTG induction and 5 h additional subculture, 100 µL cells were removed and diluted by a factor of ~100 for functional characterization, which was performed using a BD Accuri cell analyzer with a high-throughput sampler. 500 µL of the remaining culture was then added to 13.3 µL of 250 mM 1M7 or 13.3 µL of DMSO (control solvent). Cells were returned to shaking for 3 minutes to allow 1M7 to react, then cellular RNAs were Trizol extracted and reverse transcribed using a custom reverse transcription primer specific for GFPmut3b (5’-CAACAAGAATTGGGACAACTCCAGTG-3’). Additional 5’ and 3’ sequencing adapters were then added. Following 2 × 35 bp paired-end Illumina sequencing, ß reactivities were calculated as described by Aviran *et al*.^55^. Error bars represent the standard deviation of three samples, each probed from a separate transformation on a separate day. Replicate samples were only processed in parallel during final sequencing.

## Acknowledgements

This work was supported by an NIH Director’s New Innovator Award (1DP2GM126892), an Alfred P. Sloan Research Fellowship (FG-2017–9108), Gates Foundation funds (OPP1160667), an Arizona Biomedical Research Commission New Investigator Award (ADHS16–162400), a DARPA Young Faculty Award (D17AP00026), Gordon and Betty Moore Foundation funds (#6984), and Arizona State University funds to A.A.G; an NIH Director’s Pioneer Award (1DP1GM133052–01), Office of Naval Research funds (N00014–16-1–2410), and NSF funds (CCF-1317291, MCB-1540214) to P.Y.; an NSF CAREER award to J. B.L. (1452441); Defense Threat Reduction Agency funds (HDTRA1–14-1–0006), Air Force Office of Scientific Research funds (FA9550–14-1–0060), and Paul G. Allen Frontiers Group funds to J.J.C; and BMBF funds (Erasynbio project UNACS – 031L0011) and DFG funds (SFB 10327TPA2) to F.C.S. J.K. acknowledges a Wyss Institute Director’s Cross-Platform Fellowship. The views, opinions, and/or findings contained in this article are those of the authors and should not be interpreted as representing the official views or policies, either expressed or implied, of DARPA or the Department of Defense.

## AUTHOR CONTRIBUTIONS

J.K. and A.A.G. designed the toehold repressors and ribocomputing devices. J.K., M.T., and A.A.G. performed experiments for the toehold repressors and ribocomputing devices. Y.Z. and A.A.G. designed the 3WJ repressors and ribocomputing devices. Y.Z. performed experiments for the 3WJ repressors and their ribocomputing devices. P.D.C. performed SHAPE-Seq measurements. J.K., Y.Z., A.A.G., and P.D.C. analyzed the data. J.K., Y.Z., P.D.C. and A.A.G. wrote the manuscript. J.K., A.A.G., P.D.C., J.B.L., and P.Y. edited the manuscript. A.A.G., P.Y., J.B.L., J.J.C., P.A.S., and F.C.S. supervised the research.

## COMPETING FINANCIAL INTERESTS

US provisional patents have been filed by J.K., A.A.G., & P.Y. and Y.Z. & A.A.G. based on this work. P.Y. is the co-founder of Ultivue Inc. and NuProbe Global.

## REFERENCES

1. M. J. Smanski, H. Zhou, J. Claesen, B. Shen, M. A. Fischbach & C. A. Voigt, “Synthetic biology to access and expand nature’s chemical diversity,” Nature Reviews Microbiology 14, 135 (2016).

2. T. Kitada, B. DiAndreth, B. Teague & R. Weiss, “Programming gene and engineered-cell therapies with synthetic biology,” Science 359, eaad1067 (2018).

3. R. A. Le Feuvre & N. S. Scrutton, “A living foundry for Synthetic Biological Materials: A synthetic biology roadmap to new advanced materials,” Synthetic and Systems Biotechnology 3, 105–112 (2018).

4. D. R. Nielsen & T. S. Moon, “From promise to practice: The role of synthetic biology in green chemistry,” EMBO reports 14, 1034–1038 (2013).

5. T. S. Gardner, C. R. Cantor & J. J. Collins, “Construction of a genetic toggle switch in Escherichia coli,” Nature 403, 339–342 (2000).

6. M. B. Elowitz & S. Leibler, “A synthetic oscillatory network of transcriptional regulators,” Nature 403, 335–338 (2000).

7. T. Danino, O. Mondragon-Palomino, L. Tsimring & J. Hasty, “A synchronized quorum of genetic clocks,” Nature 463, 326–330 (2010).

8. A. Prindle, P. Samayoa, I. Razinkov, T. Danino, L. S. Tsimring & J. Hasty, “A sensing array of radically coupled genetic ‘biopixels’,” Nature 481, 39–44 (2012).

9. S. Ausländer, D. Ausländer, M. Müller, M. Wieland & M. Fussenegger, “Programmable single-cell mammalian biocomputers,” Nature 487, 123–127 (2012).

10. T. S. Moon, C. Lou, A. Tamsir, B. C. Stanton & C. A. Voigt, “Genetic programs constructed from layered logic gates in single cells,” Nature 491, 249–253 (2012).

11. M. N. Win & C. D. Smolke, “Higher-order cellular information processing with synthetic RNA devices,” Science 322, 456–460 (2008).

12. P. Siuti, J. Yazbek & T. K. Lu, “Synthetic circuits integrating logic and memory in living cells,” Nat Biotechnol 31, 448–452 (2013).

13. L. Yang, A. A. K. Nielsen, J. Fernandez-Rodriguez, C. J. McClune, M. T. Laub, T. K. Lu & C. A. Voigt, “Permanent genetic memory with > 1-byte capacity,” Nature Methods 11, 1261–1266 (2014).

14. J. Bonnet, P. Subsoontorn & D. Endy, “Rewritable digital data storage in live cells via engineered control of recombination directionality,” Proc Natl Acad Sci U S A 109, 8884–8889 (2012).

15. R. Daniel, J. R. Rubens, R. Sarpeshkar & T. K. Lu, “Synthetic analog computation in living cells,” Nature 497, 619–623 (2013).

16. F. Farzadfard & T. K. Lu, “Synthetic biology. Genomically encoded analog memory with precise in vivo DNA writing in living cell populations,” Science 346, 1256272 (2014).

17. N. Roquet, A. P. Soleimany, A. C. Ferris, S. Aaronson & T. K. Lu, “Synthetic recombinase-based state machines in living cells,” Science 353, aad8559 (2016).

18. D. T. Riglar, T. W. Giessen, M. Baym, S. J. Kerns, M. J. Niederhuber, R. T. Bronson, J. W. Kotula, G. K. Gerber, J. C. Way & P. A. Silver, “Engineered bacteria can function in the mammalian gut long-term as live diagnostics of inflammation,” Nature Biotechnology 35, 653–658 (2017).

19. M. O. Din, T. Danino, A. Prindle, M. Skalak, J. Selimkhanov, K. Allen, E. Julio, E. Atolia, L. S. Tsimring, S. N. Bhatia & J. Hasty, “Synchronized cycles of bacterial lysis for in vivo delivery,” Nature 536, 81–85 (2016).

20. Z. Xie, L. Wroblewska, L. Prochazka, R. Weiss & Y. Benenson, “Multi-Input RNAi-Based Logic Circuit for Identification of Specific Cancer Cells,” Science 333, 1307–1311 (2011).

21. B. C. Stanton, A. A. Nielsen, A. Tamsir, K. Clancy, T. Peterson & C. A. Voigt, “Genomic mining of prokaryotic repressors for orthogonal logic gates,” Nat Chem Biol 10, 99–105 (2014).

22. A. A. Nielsen, B. S. Der, J. Shin, P. Vaidyanathan, V. Paralanov, E. A. Strychalski, D. Ross, D. Densmore & C. A. Voigt, “Genetic circuit design automation,” Science 352, aac7341 (2016).

23. L. B. Andrews, A. A. K. Nielsen & C. A. Voigt, “Cellular checkpoint control using programmable sequential logic,” Science 361, aap8987 (2018).

24. B. H. Weinberg, N. T. H. Pham, L. D. Caraballo, T. Lozanoski, A. Engel, S. Bhatia & W. W. Wong, “Large-scale design of robust genetic circuits with multiple inputs and outputs for mammalian cells,” Nature Biotechnology 35, 453–462 (2017).

25. X. J. Gao, L. S. Chong, M. S. Kim & M. B. Elowitz, “Programmable protein circuits in living cells,” Science 361, 1252–1258 (2018).

26. F. J. Isaacs, D. J. Dwyer, C. M. Ding, D. D. Pervouchine, C. R. Cantor & J. J. Collins, “Engineered riboregulators enable post-transcriptional control of gene expression,” Nature Biotechnology 22, 841–847 (2004).

27. J. B. Lucks, L. Qi, V. K. Mutalik, D. Wang & A. P. Arkin, “Versatile RNA-sensing transcriptional regulators for engineering genetic networks,” Proceedings of the National Academy of Sciences of the United States of America 108, 8617–8622 (2011).

28. M. K. Takahashi & J. B. Lucks, “A modular strategy for engineering orthogonal chimeric RNA transcription regulators,” Nucleic Acids Research 41, 7577–7588 (2013).

29. V. K. Mutalik, L. Qi, J. C. Guimaraes, J. B. Lucks & A. P. Arkin, “Rationally designed families of orthogonal RNA regulators of translation,” Nature Chemical Biology 8, 447–454 (2012).

30. A. A. Green, P. A. Silver, J. J. Collins & P. Yin, “Toehold switches: de-novo-designed regulators of gene expression,” Cell 159, 925–939 (2014).

31. B. Yurke, A. J. Turberfield, A. P. Mills, Jr., F. C. Simmel & J. L. Neumann, “A DNA-fuelled molecular machine made of DNA,” Nature 406, 605–608 (2000).

32. K. Pardee, A. A. Green, T. Ferrante, D. E. Cameron, A. DaleyKeyser, P. Yin & J. J. Collins, “Paper-based synthetic gene networks,” Cell 159, 940–954 (2014).

33. A. A. Green, J. Kim, D. Ma, P. A. Silver, J. J. Collins & P. Yin, “Complex cellular logic computation using ribocomputing devices,” Nature 548, 117–121 (2017).

34. K. Pardee, A. A. Green, T. Ferrante, D. E. Cameron, A. DaleyKeyser, P. Yin & J. J. Collins, “Paper-based synthetic gene networks,” Cell 159, 940–954 (2014).

35. K. Pardee, A. A. Green, M. K. Takahashi, D. Braff, G. Lambert, J. W. Lee, T. Ferrante, D. Ma, N. Donghia, M. Fan, N. M. Daringer, I. Bosch, D. M. Dudley, D. H. O’Connor, L. Gehrke & J. J. Collins, “Rapid, Low-Cost Detection of Zika Virus Using Programmable Biomolecular Components,” Cell 165, 1255–1266 (2016).

36. D. Ma, L. Shen, K. Wu, C. W. Diehnelt & A. A. Green, “Low-cost detection of norovirus using paper-based cell-free systems and synbody-based viral enrichment,” Synthetic Biology 3, ysy018 (2018).

37. M. K. Takahashi, X. Tan, A. J. Dy, D. Braff, R. T. Akana, Y. Furuta, N. Donghia, A. Ananthakrishnan & J. J. Collins, “A low-cost paper-based synthetic biology platform for analyzing gut microbiota and host biomarkers,” Nature Communications 9, 3347 (2018).

38. M. W. Gander, J. D. Vrana, W. E. Voje, J. M. Carothers & E. Klavins, “Digital logic circuits in yeast with CRISPR-dCas9 NOR gates,” Nature Communications 8, 15459 (2017).

39. B. C. Stanton, A. A. K. Nielsen, A. Tamsir, K. Clancy, T. Peterson & C. A. Voigt, “Genomic mining of prokaryotic repressors for orthogonal logic gates,” Nature Chemical Biology 10, 99–105 (2014).

40. L. A. Gilbert, M. H. Larson, L. Morsut, Z. Liu, G. A. Brar, S. E. Torres, N. Stern-Ginossar, O. Brandman, E. H. Whitehead, J. A. Doudna, W. A. Lim, J. S. Weissman & L. S. Qi, “CRISPR-mediated modular RNA-guided regulation of transcription in eukaryotes,” Cell 154, 442–451 (2013).

41. J. N. Zadeh, C. D. Steenberg, J. S. Bois, B. R. Wolfe, M. B. Pierce, A. R. Khan, R. M. Dirks & N. A. Pierce, “NUPACK: Analysis and design of nucleic acid systems,” Journal of computational chemistry 32, 170–173 (2011).

42. K. E. Watters, T. R. Abbott & J. B. Lucks, “Simultaneous characterization of cellular RNA structure and function with in-cell SHAPE-Seq,” Nucleic Acids Research 44, e12 (2016).

43. J. Kim, P. Yin & A. A. Green, “Ribocomputing: Cellular Logic Computation Using RNA Devices,” Biochemistry 57, 883–885 (2018).

44. X. Chen, “Expanding the Rule Set of DNA Circuitry with Associative Toehold Activation,” Journal of the American Chemical Society 134, 263–271 (2012).

45. A. J. Genot, J. Bath & A. J. Turberfield, “Combinatorial Displacement of DNA Strands: Application to Matrix Multiplication and Weighted Sums,” Angewandte Chemie International Edition 52, 1189–1192 (2013).

46. P. D. Carlson, M. E. Evans, A. M. Yu, E. J. Strobel & J. B. Lucks, “SnapShot: RNA Structure Probing Technologies,” Cell 175, 600–600.e1 (2018).

47. G. Rodrigo, T. E. Landrain & A. Jaramillo, “De novo automated design of small RNA circuits for engineering synthetic riboregulation in living cells,” Proceedings of the National Academy of Sciences of the United States of America 109, 15271–15276 (2012).

48. L. S. Qi, M. H. Larson, L. A. Gilbert, J. A. Doudna, J. S. Weissman, A. P. Arkin & W. A. Lim, “Repurposing CRISPR as an RNA-guided platform for sequence-specific control of gene expression,” Cell 152, 1173–1183 (2013).

49. D. Bikard, W. Jiang, P. Samai, A. Hochschild, F. Zhang & L. A. Marraffini, “Programmable repression and activation of bacterial gene expression using an engineered CRISPR-Cas system,” Nucleic Acids Res 41, 7429–7437 (2013).

50. J. Chappell, M. K. Takahashi & J. B. Lucks, “Creating small transcription activating RNAs,” Nature Chemical Biology 11, 214–220 (2015).

51. S. Meyer, J. Chappell, S. Sankar, R. Chew & J. B. Lucks, “Improving fold activation of small transcription activating RNAs (STARs) with rational RNA engineering strategies,” Biotechnology and Bioengineering 113, 216–225 (2016).

52. J. Chappell, A. Westbrook, M. Verosloff & J. B. Lucks, “Computational design of small transcription activating RNAs for versatile and dynamic gene regulation,” Nature Communications 8, 1051 (2017).

53. D. G. Gibson, L. Young, R.-Y. Chuang, J. C. Venter, C. A. Hutchison & H. O. Smith, “Enzymatic assembly of DNA molecules up to several hundred kilobases,” Nature Methods 6, 343–345 (2009).

54. J. B. Andersen, C. Sternberg, L. K. Poulsen, S. P. Bjorn, M. Givskov & S. Molin, “New unstable variants of green fluorescent protein for studies of transient gene expression in bacteria,” Applied and Environmental Microbiology 64, 2240–2246 (1998).

55. S. Aviran, J. B. Lucks & L. Pachter, “RNA structure characterization from chemical mapping experiments,” arXiv preprint arXiv:1106.5061 (2011).

