## Supplementary Information for "*De-Novo*-Designed Translational Repressors for Multi-Input Cellular Logic"

*Jongmin Kim[1,2]*, Yu Zhou[3]*, Paul Carlson[4,5], Mario Teichmann[6], Friedrich C. Simmel[6,7], Pamela A. Silver[1,8], James J. Collins[1,9,10], Julius B. Lucks[5,11], Peng Yin[1,8], Alexander A. Green[3]*
[1]Wyss Institute for Biologically Inspired Engineering, Harvard University, Boston, MA 02115, USA. [2]Integrative Biosciences and Biotechnology, Pohang University of Science and Technology, Pohang, Gyeongbuk 37673, Republic of Korea. [3]Biodesign Center for Molecular Design and Biomimetics, The Biodesign Institute and School of Molecular Sciences, Arizona State University, Tempe, AZ 85287, USA. [4]Robert F. Smith School of Chemical and Biomolecular Engineering, Cornell University, Ithaca, NY, 14853, USA. [5]Center for Synthetic Biology, Northwestern University, Evanston, IL, 60208, USA. [6]Physics Department E14 and ZNN/WSI, Technische Universität München, Garching 85748, Germany. [7]Nanosystems Initiative Munich, Munich 80799, Germany. [8]Department of Systems Biology, Harvard Medical School, Boston, MA 02115, USA. [9]Institute for Medical Engineering and Science, Department of Biological Engineering, and Synthetic Biology Center, Massachusetts Institute of Technology, Cambridge, MA 02139, USA. [10]Broad Institute of MIT and Harvard, Cambridge, MA 02142, USA. [11]Department of Chemical and Biological Engineering, Northwestern University, Evanston, IL, 60208, USA
**These authors contributed equally to this work*

**Design of Toehold Repressors and Three-Way Junction (3WJ) Repressors**
Nucleotide-level design schematics of the toehold and 3WJ repressors are shown in Figure S1. Toehold repressors were designed to provide a 15-nt toehold region for trigger binding and refold into a repressing hairpin structure identical to that used in toehold switches (Figure S1a). The repressed hairpin had a 12-nt loop and the top 3-bp of the stem was specified to contain only A-U base pairing. This latter condition was previously found to be a common characteristic of multiple high-performance toehold switches[1]. An additional 12-bp stem domain (**c*** shown in purple) was used to ensure that the repressing hairpin structure would only form upon binding to the trigger RNA. A 4-nt single-stranded region (AAAC) was used upstream of the main ribosomal binding site (RBS) sequence (AGAGGAGA) to ensure efficient translation of the output gene in the absence of the trigger. These design considerations resulted in a 30-nt long hairpin region for the switch RNA in its active translation state. Three bulges were included in this hairpin structure at 8-nt increments to discourage transcriptional termination through the strong secondary structure. As part of the RNA design, the switch RNA sequence was considered up to the 30 nts following the repressed hairpin structure, which included a 21-nt linker previously used for toehold switches and the first 9 nts of GFPmut3b (see Supplementary Table S1). Trigger RNAs for the toehold repressors were designed with a 5' hairpin region to increase RNA stability followed by the **c**, **b**, and **a** domains responsible for binding to the switch RNA. The trigger RNA was also designed with the 47-nt T7 terminator sequence (see Supplementary Table S1). Three-nucleotide spacers were added between the interaction domains and the outer hairpins as part of the trigger design.

The 3WJ repressor switch RNAs were designed using the core sequence of first-generation toehold switch number 1.[1] This core region is indicated by the gray and black bases within the hairpin structure shown in Figure S1b and has the sequence:

UUGUUAUAGUUAUGAAC**AGAGGAGA**CAUAAC**AUG**AACAA

where the RBS and start codon are shown in bold. In previous studies of toehold switch number 1, we found that this hairpin sequence provided very high translational output despite its secondary structure, which suggested that the lower stem of the structure comprising the sequence UUGUU remained predominantly unpaired when transcribed in the cell. This core translational element was integrated into the 3WJ repressor by appending binding domains **a*** and **b*** with lengths of 15 nts and 12 nts, respectively, to either side. A 7-nt single-stranded domain was added downstream of **b*** to preserve the correct reading frame followed by the 21-nt linker sequence and the first 9 nts of the GFPmut3b coding sequence (see Supplementary Table S1). The 3WJ repressor trigger RNAs featured a 17-nt toehold domain comprising the **a** domain and the last 2 nts of the **b** domain. The remaining 10 nts of the **b** domain were contained within an 8-bp stem structure and a portion of the 6-nt loop. A 3-nt single-stranded spacer region was used to separate the binding domains of the trigger RNA from the T7 terminator at the 3' end of the transcript.

Designs for the repressor libraries were generated using the NUPACK software package[2]. The toehold repressors were designed using a specified temperature of 37 °C; Serra and Turner, 1995 energy parameters[3]; and the prevented sequences AAAA, CCCC, GGGG, UUUU, KKKKKKKKKK, MMMMMMMMMM, RRRRRRRRRR, SSSSSSSSSS, WWWWWWWWWW, YYYYYYYYYY. The 3WJ repressors used a specified temperature of 37 °C; Mathews et al., 1999 energy parameters[4]; and the prevented sequences AAAA, CCCC, GGGG, UUUU, KKKKKKKK, MMMMMM, RRRRRR, SSSSSS, WWWWWW, YYYYYY.

**Toehold Repressor Forward Engineering**

A second-generation toehold repressor library was generated through an automated forward engineering procedure based on sequence-dependent thermodynamic parameters and GFP output obtained from the initial library of toehold repressors. A set of 114 easy-to-calculate thermodynamic parameters were defined as shown in Supplementary Table S9. These parameters can be broadly classified into seven different categories.

1. *MFE of RNA strands and critical RNA subsequences:* The minimum free energy (MFE) of the RNA strands and subsequences were calculated to assess the strength of the repressor secondary structures and the binding between trigger and switch RNAs.
2. *Measures of binding for critical RNA domains:* The free energy of duplex structures formed between toehold domains and 45-nt minimal trigger sequences were calculated to assess the binding strengths of these interactions.
3. *Net reaction free energies:* The net change in free energy starting from separate trigger and switch RNA strands to the trigger/switch complex.
4. *Influence of coding sequence secondary structure on translation in active state:* The MFE for the coding sequence over different subsequences of the switch RNA were used to assess the efficiency with which these regions would be translated.
5. *Influence of coding sequence secondary structure on translation in inactive state:* The MFE for the switch RNA subsequence immediately after the binding site for the trigger RNA was used to assess the degree of translation inhibition caused by sequestering the RBS and start codon in a stem loop. These parameters were also calculated over several switch RNA subsequences.
6. *Deviations of actual sequence from the ideal, design-specified secondary structure:* These parameters were used to assess the degree to which the predicted MFE structures differed from those specified in the toehold switch design. We expected that these parameters would capture, for instance, designs where base pairing within the toehold domain reduced the performance of the repressor.

7. *Measures of switch RNA stem sequence:* These parameters evaluated the strength of the switch RNA stem over different sequence ranges starting from the top or the bottom of the stem. Previous studies with toehold switches had revealed some performance improvements when a stem had A-U base pairs at particular locations. We expected that these parameters could capture such effects in the toehold repressors.

The thermodynamic parameters were calculated for 38 toehold repressors from the first-generation library using a local implementation of NUPACK with Mathews et al., 1999 energy parameters[4]. Six devices were excluded from the analysis due to their unusually low ON state expression level or high expression variability in either the ON or OFF state. A series of different linear regressions were then performed using the thermodynamic parameters and $\log_{10}$(GFP fold reduction) obtained from the repressor library. These regressions were calculated using the Matlab `regress` function, which implemented a multiple linear regression algorithm using least squares. Figure S4 provides a map of $R^2$ values obtained for all two-parameter linear regressions performed using the 114 thermodynamic parameters. This analysis identified several parameters showing stronger correlations with the experimental data and suggested extension of the approach to three-parameter regressions.

To develop a scoring function to rank potential toehold repressors, we first computed all possible combinations of up to three different thermodynamic parameters. The resulting 253,460 linear regressions were then ranked from highest to lowest $R^2$ value. Regression coefficients from the top 10 linear regressions were extracted to generate an overall scoring function. The combined regression coefficients were generated using equation (1):

$$B_i = \frac{1}{N}\sum_{j=1}^{N} C_{ij} \quad (1)$$

where $B_i$ is the combined regression coefficient for parameter $i$, $C_{ij}$ is the regression coefficient for parameter $i$ in regression $j$, and $N$ = 10 is the number of regressions combined. In cases where parameter $i$ was not used in the linear regression, the regression coefficient was equal to zero. The scoring function for ranking designs was then of the following form:

$$S_k = \sum_i B_i \Delta G_{ik} \quad (2)$$

where $S_k$ is the score computed for toehold repressor design $k$ and $\Delta G_{ik}$ is the free energy calculated for parameter $i$ for toehold repressor design $k$. Repressors predicted to offer higher performance were thus given higher scores. The final equation for the scoring function, using the parameter names from Supplementary Table S9, is:

$$\begin{aligned} S_k = {} & 0.025 \times \text{deltaG\_toeh\_binding}_k + 0.025 \times \text{deltaG\_toeh\_binding\_actual}_k \\ & + 0.003 \times \text{deltaG\_targ\_binding}_k + 0.003 \times \text{deltaG\_targ\_binding\_actual}_k \\ & + 0.377 \times \text{deltaG\_stem\_27}_k + 0.029 \times \text{dup2link\_deltaG}_k \\ & + 0.029 \times \text{dup2pos03\_deltaG}_k - 0.072 \times \text{deltaG\_stem\_26}_k \\ & - 0.021 \times \text{deltaG\_stem\_25}_k \end{aligned} \quad (3)$$

We then applied this automatically generated scoring function to a set of 265 new toehold repressor sequences produced using the same sequence and structural parameters as the first-generation library. For these designs, the values of several thermodynamic parameters were identical or nearly identical to one another across the set of sequences, enabling the scoring function to be simplified to the following expression:

$$\begin{aligned} S_k = {} & 0.050 \times \text{deltaG\_toeh\_binding}_k + 0.006 \times \text{deltaG\_targ\_binding}_k \\ & + 0.058 \times \text{dup2link\_deltaG}_k + 0.377 \times \text{deltaG\_stem\_27}_k \\ & - 0.072 \times \text{deltaG\_stem\_26}_k - 0.021 \times \text{deltaG\_stem\_25}_k \end{aligned} \quad (4)$$

The 96 new repressor sequences providing the highest scores were then assembled and tested. These devices yielded substantially improved performance compared to the first-generation library, with over 84% of the riboregulators providing at least 10-fold reductions in GFP expression compared to 48% for the initial library (Figure 2). The improvements were more striking on the high end of device performance with only one device (2%) in the first-generation library with 100-fold reduction in GFP compared to 8 devices (8%) in the second-generation set.

Analysis of the thermodynamic parameters and regression coefficients used in the scoring function provides some insight into the riboregulator characteristics favored for the second-generation library. The terms deltaG_toeh_binding and deltaG_targ_binding provided the largest overall contributions to the device score and encouraged selection of sequences with weaker binding through the toehold and minimal trigger regions. In effect, these parameters favored devices with low GC content in the toehold and trigger region. The next most important term was dup2link_deltaG, which also had a positive regression coefficient. This term measures the secondary structure of the switch RNA region being translated when the riboregulator is in its ON state. Accordingly, this term favored devices with low secondary structure in this region to encourage efficient translation of the output protein. The last three terms in the scoring function are related to the free energy of subsequences near the top of the switch RNA stem. The deltaG_stem_27 has a positive regression coefficient and thus selected for designs having low GC content in the top four base pairs at the top of the switch stem. In contrast, deltaG_stem_26 and deltaG_stem_25, which assessed five and six base-pair upper stems, respectively, had negative coefficients and thus favored stronger base pairing in the upper stem. Taken together, the last three parameters assigned higher scores to devices having low GC content in the top four base pairs of the stem and relatively higher GC content in the base pairs 5 and 6 nts from the stem top.

**SHAPE-Seq Measurements of a Second 3WJ Repressor**

We studied a second 3WJ repressor that demonstrated an unusual pattern of repression with increasing trigger length (Figure S5, see Supplementary Table S5 for sequence information). In this case, expression of a trigger with an 18-nt interaction length showed weak (3-fold) repression (Figure S5b). However, a small increase in interaction length to 20 nt dramatically increased the repression efficiency to 17-fold. We performed in-cell SHAPE-Seq on the switch and switch-trigger complexes to identify structural explanations for these performance differences. As expected, SHAPE-Seq of the switch RNA alone showed high reactivity across the molecule, corresponding to a translationally-active state (Figure S5c). Similarly, the functional 20-nt trigger variant showed a reactivity pattern consistent with 3WJ formation (Figure S5e). However, the poorly-repressing 18-nt variant only showed low reactivities within the **b-b*** binding region, with high reactivities observed elsewhere. This suggests that the trigger only efficiently forms a duplex at **b-b***, leaving the hairpin de-stabilized and accessible for ribosome binding. The modest repression from this trigger likely results from inhibition of translational elongation through **b-b***, rather than inhibition of translation initiation through 3WJ formation.

**Design of Repressor-Based Ribocomputing Devices**

A modified toehold repressor design was used for use in two-input NAND devices (see Figure S7). The toehold domain length was increased to 16 nts and the stem of the gate RNA was reduced to 24 nts. These changes enabled the trigger RNA sequence to be divided into two segments of similar length. Two different segment lengths were used in devices. Inputs A1 and

A2 used **a1** and **a2** domains with lengths of 24 and 16 nts, respectively (Figure S7a). Inputs B1 and B2 used **b1** and **b2** domains that were both 20 nts in length (Figure S7b). The repressed fold of the gate RNA retained the same stem secondary structure but had a loop domain of 18 nts rather than 12 nts. The expanded loop domain was implemented to reduce the potential for unintended repression caused by disruption of the bottom of the gate RNA stem, which could occur for some input RNA sequences. The input RNAs were also designed to hybridize through a **u** domain of 23 nts and form an RNA duplex with a single-nucleotide bulge at the midpoint.

**Design of 3WJ Repressor Two-Input NOR Gate RNA**

The two-input NOR gate RNA was designed using two hairpin sensor modules upstream of a 3WJ repressor hairpin (Figure S9a). Each sensor module consisted of a 15-nt toehold domain followed by an 18-bp stem. The loop region of this stem contained a sequestered internal trigger for the downstream repressor hairpin. The trigger sequence length was reduced by 5 nts within the **x** domain compared to that used for the original library characterization to reduce the probability of leakage from the internal trigger. The two sensor modules and the repressor hairpin were separated by 18-nt spacer sequences having small hairpin secondary structures. These spacers were used to ligate the three modules together through Gibson assembly during plasmid construction and their hairpin structures were used to reduce the effective distance between the intramolecular triggers and the downstream 3WJ repressor hairpin. Input RNAs complementary to the toehold and stem domain of the two sensor modules were also designed. These input RNAs had 33-nt binding domains to the gate RNA and were flanked by a 5’ hairpin structure and the T7 terminator sequence.

**Design of 3WJ Repressor NAND Gate RNAs**

The NAND gate RNAs based on 3WJ repressors were generated by taking the core regulatory sequence of the repressors running from the 5’ end of the **a*** domain through to the nucleotide immediately before the 21-nt linker sequence. Since this core regulatory sequence had a length of 73 nts, spacers of $3n+2$, where $n$ is a non-negative integer, were used to connect different 3WJ repressors together. Spacers of this length enabled successive repressor modules to remain in-frame through the full length of the gate RNA. For testing purposes, we selected 11- and 17-nt spacers to insert between different 3WJ repressor hairpins. The spacers were designed using NUPACK to have single-stranded secondary structures when flanked by the two repressor hairpins. Figure S9b shows the programmed secondary structure of the four-input NAND gate RNA used in Figure 6 of the main text.

### REFERENCES


1. A. A. Green, P. A. Silver, J. J. Collins & P. Yin, "Toehold switches: de-novo-designed regulators of gene expression," *Cell* **159**, 925-939 (2014).
2. J. N. Zadeh, C. D. Steenberg, J. S. Bois, B. R. Wolfe, M. B. Pierce, A. R. Khan, R. M. Dirks & N. A. Pierce, "NUPACK: Analysis and design of nucleic acid systems," *Journal of computational chemistry* **32**, 170-173 (2011).
3. M. J. Serra & D. H. Turner, "Predicting thermodynamic properties of RNA," *Methods in enzymology* **259**, 242-261 (1995).
4. D. H. Mathews, J. Sabina, M. Zuker & D. H. Turner, "Expanded sequence dependence of thermodynamic parameters improves prediction of RNA secondary structure," *Journal of molecular biology* **288**, 911-940 (1999).

### Supplementary Figures

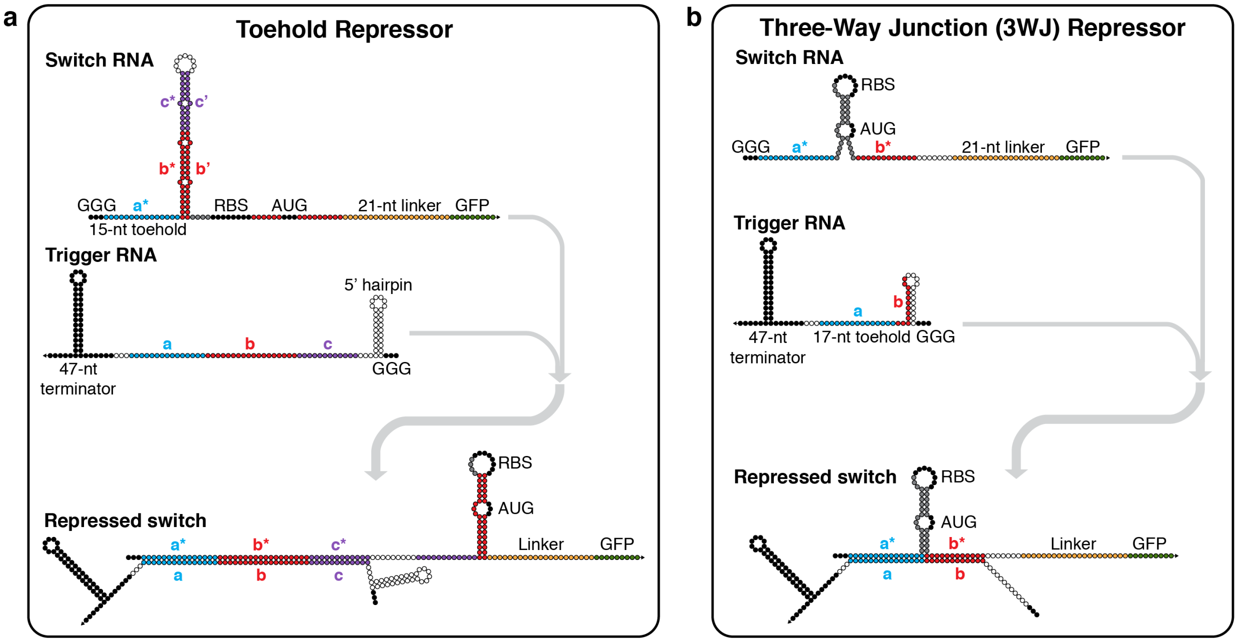


**Figure S1 | Nucleotide-level schematics of toehold repressors and three-way junction (3WJ) repressors. a**, Toehold repressors employ a switch RNA with a 15-nt toehold and a 30-nt stem to interact with a trigger RNA with a 45-nt single-stranded region. Binding of the trigger causes formation of a translation-repressing hairpin structure. **b**, 3WJ repressors employ a conserved hairpin structure in the switch RNA that places binding domains **a*** and **b*** in close proximity and enables effective translation. Binding of the trigger RNA through a toehold-mediated interaction forms a 3WJ structure that represses translation. Black bases designate sequences that are biologically conserved (e.g. terminators, RBS, start codons). White bases indicate sequences determined by NUPACK based on the specified secondary structure. Gray bases indicate sequences derived from previous riboregulators.

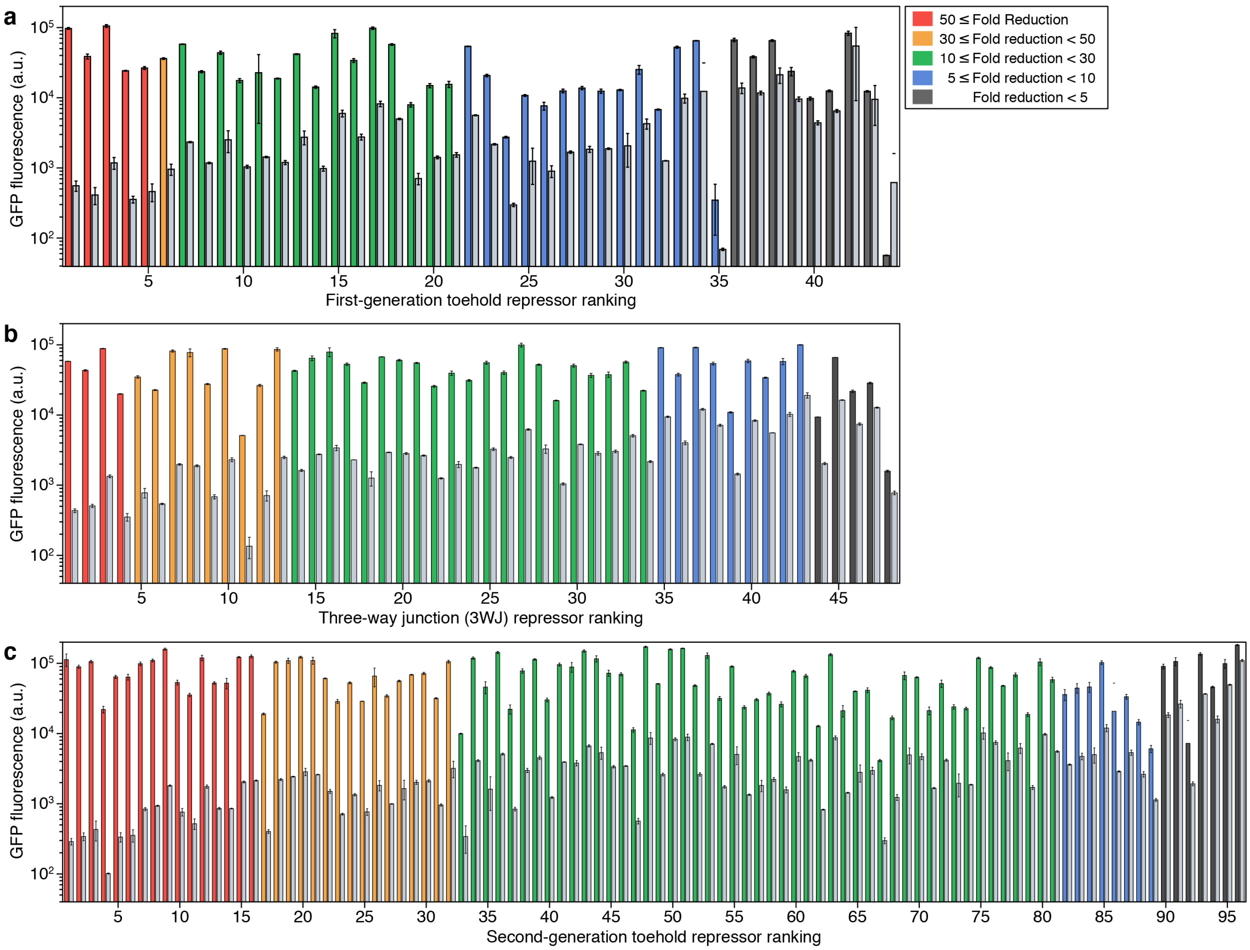


**Figure S2 | GFP fluorescence of the toehold and 3WJ repressor libraries. a-c**, GFP fluorescence levels measured via flow cytometry for the switch RNA expressed with a non-cognate trigger with high secondary structure (colored bars, ON state) and with the cognate trigger (gray bars, OFF state) for the first-generation toehold repressors (a), 3WJ repressors (b), and second-generation toehold repressors (c).

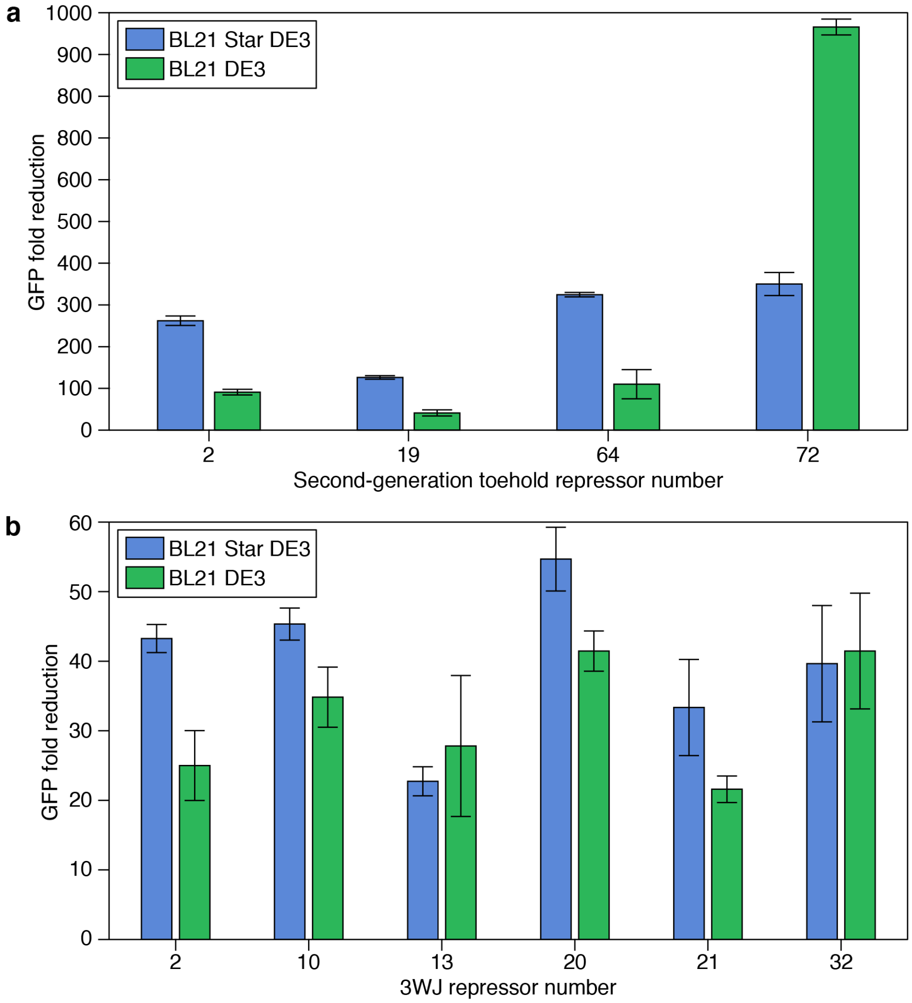


**Figure S3 | GFP fold reduction measured from synthetic repressors in different *E. coli* strains. a**, Comparison of GFP fold reduction for second-generation toehold repressors in *E. coli* BL21 Star DE3, which is RNase deficient, with *E. coli* DE3, which has wild-type RNase levels. The toehold repressors exhibit device dependent variations with strain but provide >40-fold reduction levels. Cells measured via flow cytometry after 4 hours of induction with 0.1 mM IPTG. **b**, Comparison of GFP fold reduction in the two strains for 3WJ repressors. The devices show comparable fold reductions in both strains. Cells were measured via flow cytometry after 5 hours of induction with 0.1 mM IPTG.

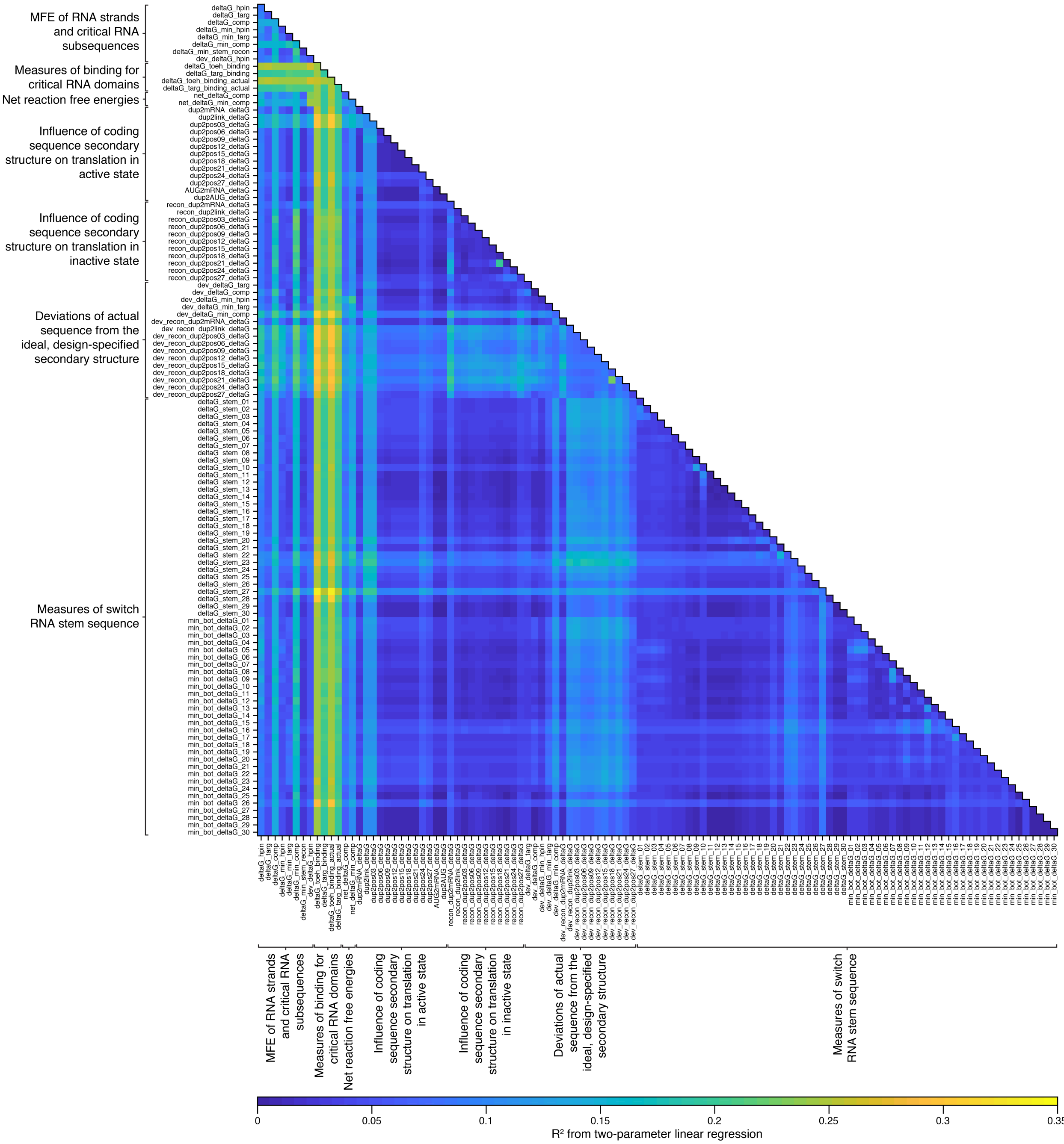


**Figure S4 | Map of $R^2$ values of two-parameter linear regressions for the first-generation toehold repressors.** Linear regressions were performed on 6,555 combinations of two thermodynamic parameters against the experimental GFP fold reduction values. Hotspots with stronger correlations to device performance can be observed for multiple parameters, such as measures of binding for critical RNA domains.

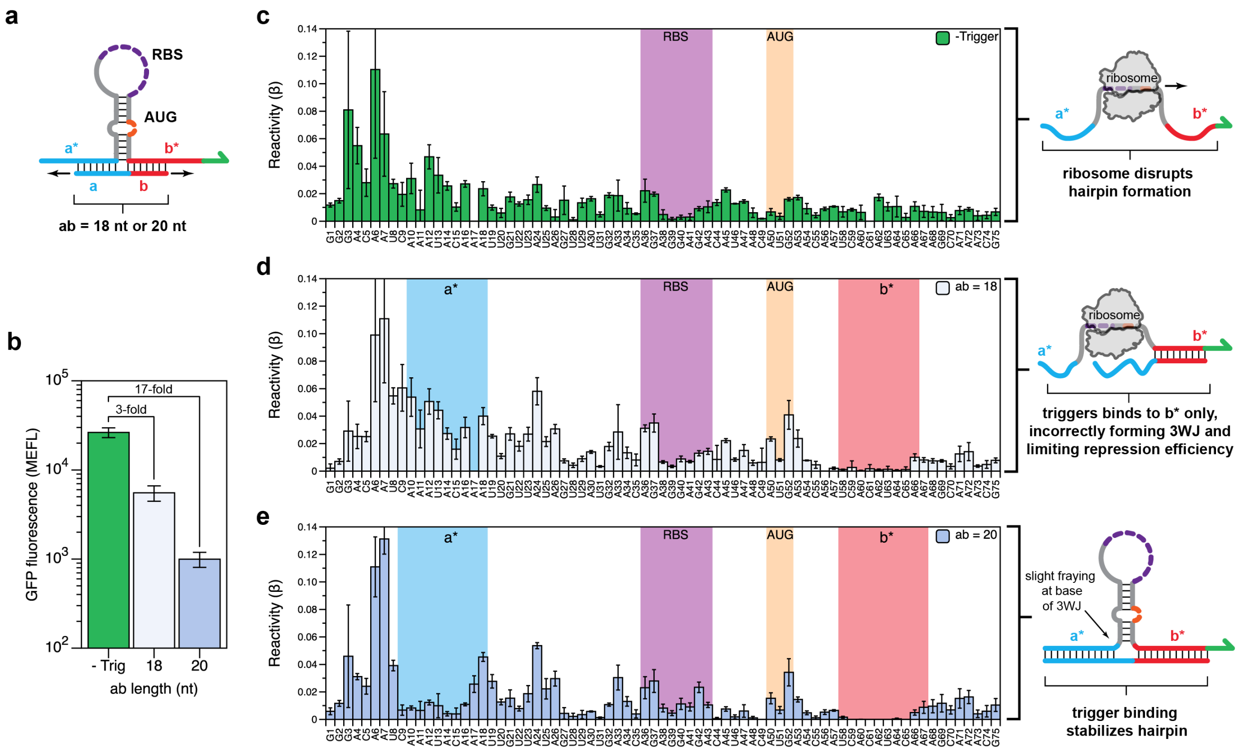


**Figure S5 I In-cell SHAPE-Seq characterization of trigger variants with varying repression efficiencies. a**, Design schematic for testing 3WJ repressor variants. A 3WJ repressor switch RNA was characterized using in-cell SHAPE-Seq, either expressed alone or co-expressed with a trigger RNA. Two triggers were tested, with designed binding lengths (**ab**) of 18 nt or 20 nt. **b**, Functional characterization of switch RNA expressed without trigger (green) and with triggers of increasing interaction length (blue). Weak repression (ON/OFF = 3) is observed when **ab** = 18 nt. Repression efficiency increases dramatically (ON/OFF = 17) when **ab** is increased to 20 nt. **c**, In-cell SHAPE-Seq reactivity profile of the switch RNA expressed alone. A trend of high reactivities is observed across the molecule, consistent with the design hypothesis that the switch hairpin can be disrupted by ribosome binding and actively translated. **d**, In-cell SHAPE-Seq reactivity profile of the switch RNA co-expressed with a poorly-repressing trigger RNA (**ab** = 18 nt). Drops in reactivity are only observed within the **b-b*** interaction domain, suggesting that trigger binding does not occur across the predicted 3WJ. Improper formation of the 3WJ is the likely cause of the weak repression efficiency for this switch-trigger pair. **e**, In-cell SHAPE-Seq reactivity profile of the switch RNA co-expressed with a longer trigger RNA (**ab** = 20 nt) showing improved repression efficiency compared to the shorter trigger (d). This length variant shows a reactivity profile more consistent with proper 3WJ formation, with reactivity drops observed at the **a-a*** and **b-b*** interaction regions, and within the switch hairpin. The RBS and start codon (AUG) positions are indicated.

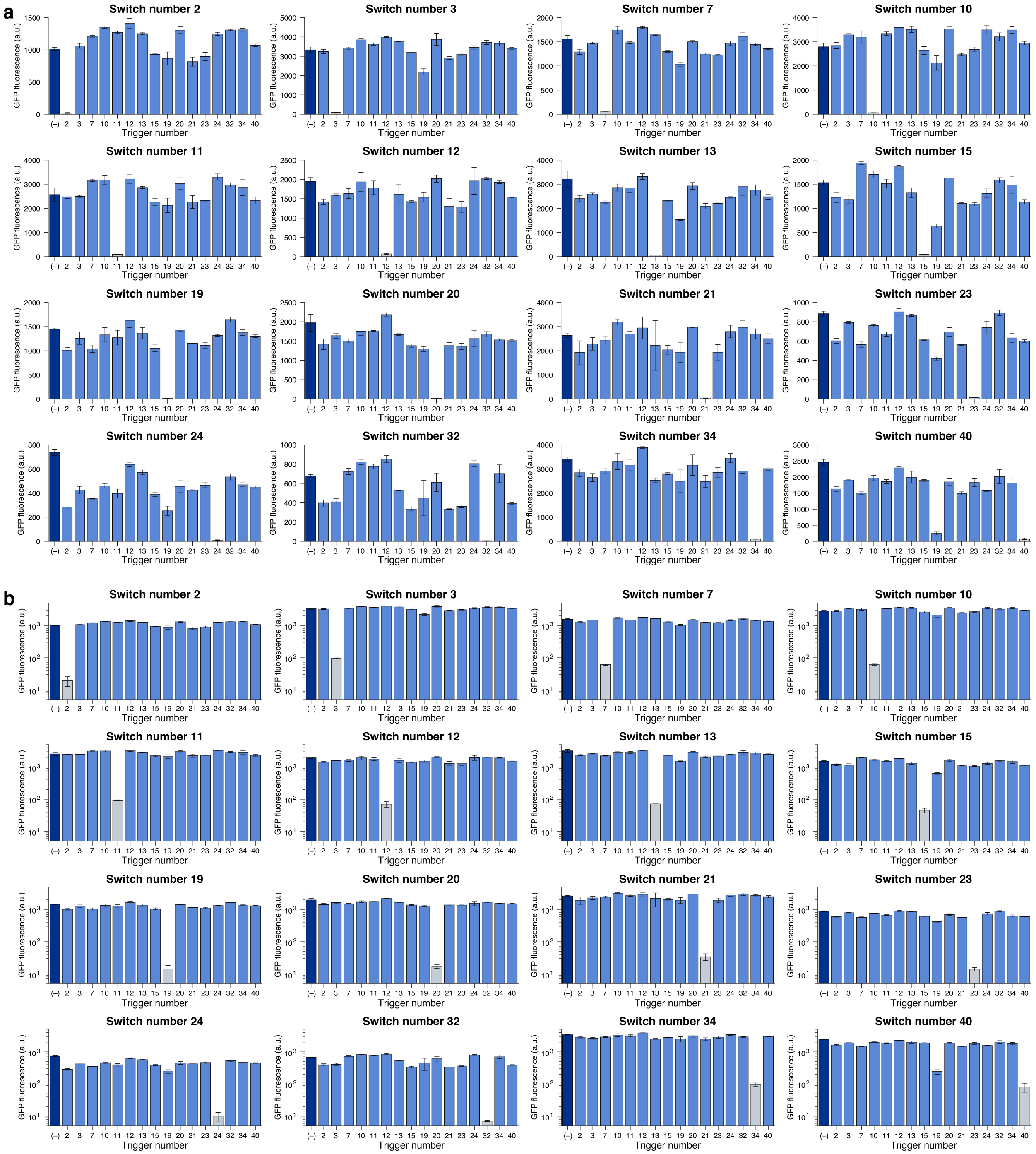


**Figure S6 | GFP fluorescence levels for 3WJ repressor orthogonality measurements. a-b**, Linear-scale (a) and logarithmic-scale (b) GFP fluorescence intensities from orthogonality measurements of 16 3WJ repressors after 3 hours of induction. Each switch was tested against the same panel of 17 different cognate (gray bars) and non-cognate trigger RNAs. The non-cognate trigger "(–)" is an RNA with high secondary structure (dark blue bars), while the other non-cognate RNAs are from other 3WJ repressors (light blue bars).

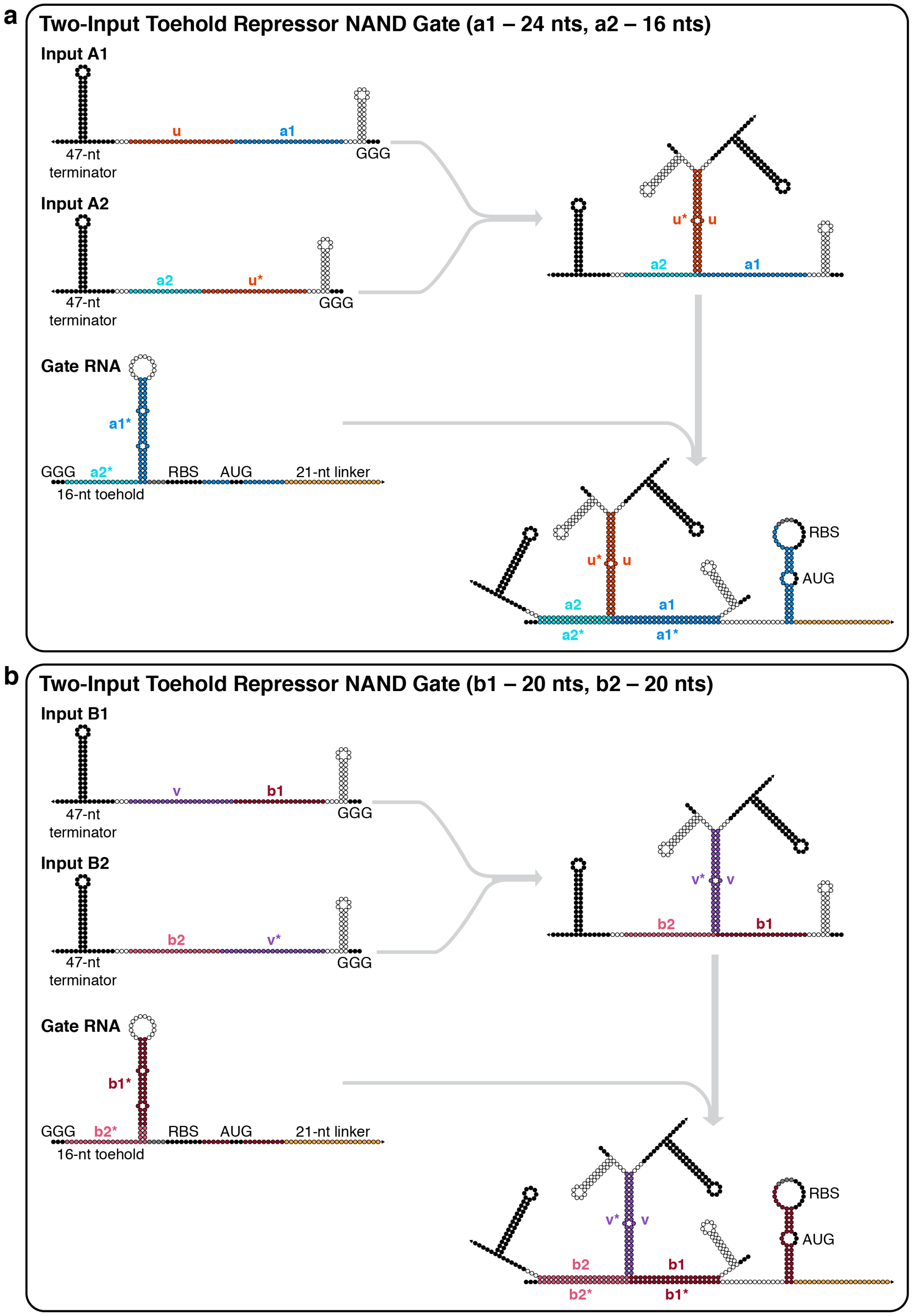


**Figure S7 | Nucleotide-level schematics for toehold-repressor-based NAND ribocomputing devices. a-b**, The two-input toehold repressor NAND gate features a modified switch RNA design and employs input RNAs that hybridize through complementary **u-u*** domains (a) and **v-v*** domains (b). The trigger RNA sequence is divided into separate segments of 20 nts each (a) or 16 nts and 24 nts (b).

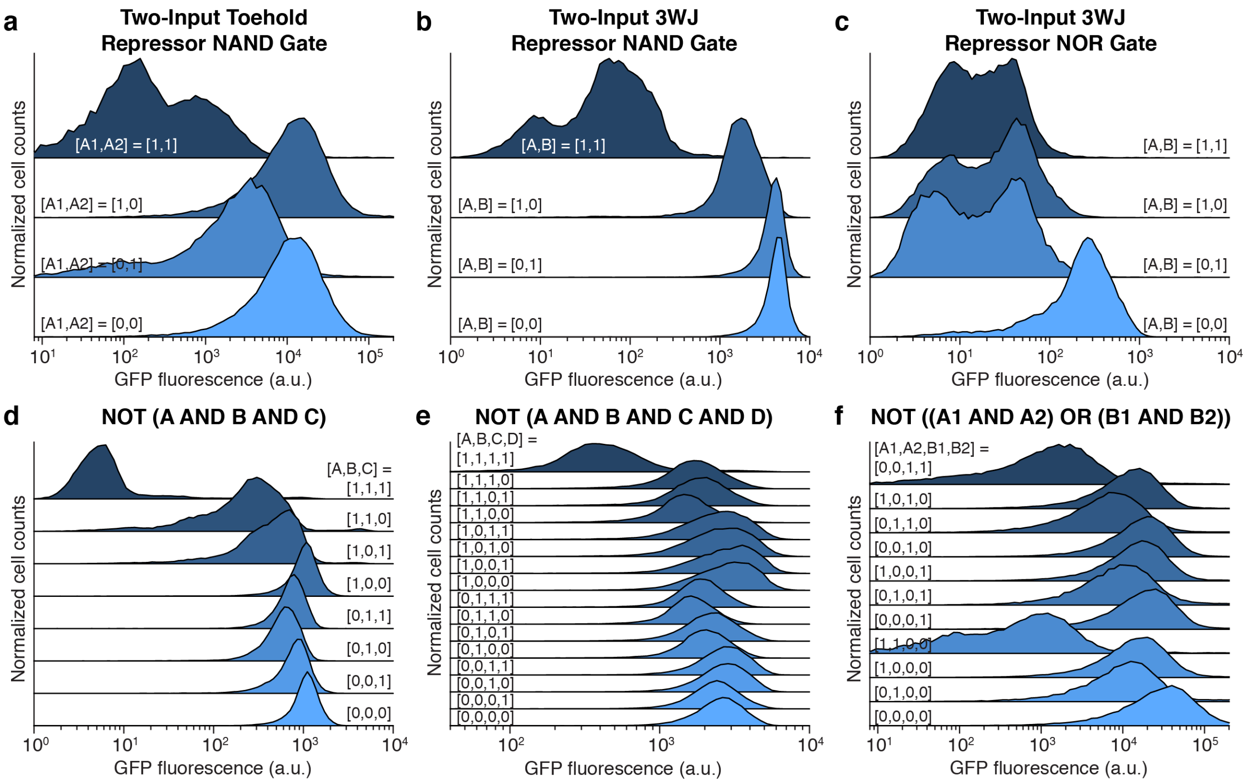


**Figure S8 | Cell population distributions for ribocomputing logic circuits. a**, Two-input toehold repressor NAND gate from Figure 5d-f. **b**, Two-input 3WJ repressor NAND gate from Figure 5g-i. **c**, Two-input 3WJ repressor NOR gate from Figure 5j-l. **d**, Three-input 3WJ repressor NAND gate from Figure 6a-c. **e**, Four-input 3WJ repressor NAND gate from Figure 6d-f. **f**, Toehold repressor NOT ((A1 AND A2) OR (B1 AND B2)) gate from Figure 6g-i.

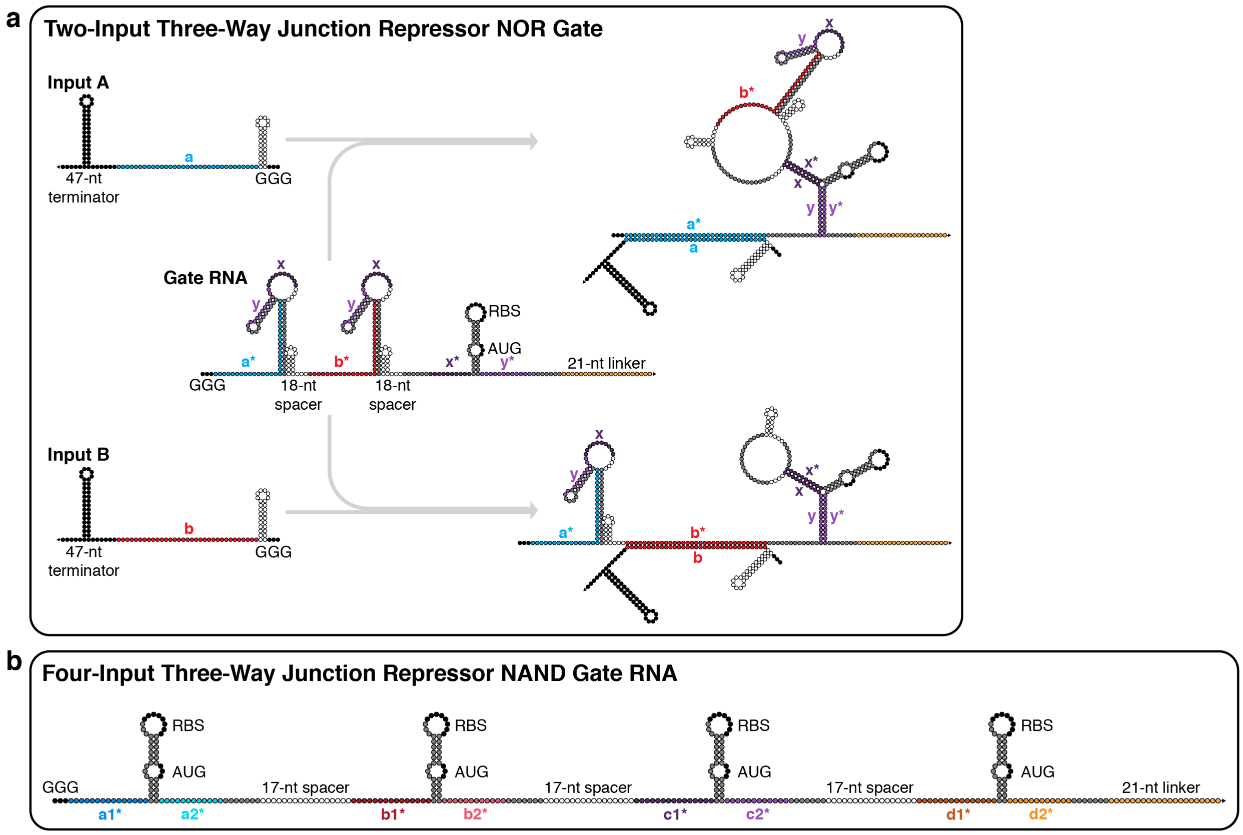


**Figure S9 | Nucleotide-level schematics for 3WJ-repressor-based ribocomputing devices.** **a**, The two-input 3WJ repressor NOR gate employs two input-sensing hairpins with loop-confined triggers for a downstream 3WJ repressor hairpin. **b**, The programmed secondary structure for four-input 3WJ repressor NAND gate RNA. Single-stranded 17-nt spacers separate four 3WJ repressor hairpins and do not encode in-frame stop codons.
