## Supplementary Tables for "*De-Novo*-Designed Translational Repressors for Multi-Input Cellular Logic"

**Table S1. PCR primers and other sequences used for experiments**

| Name | Sequence |
|---|---|
| T7 promoter | TAATACGACTCACTATA[GGG] |
| T7 terminator | TAGCATAACCCCTTGGGGCCTCTAAACGGGTCTTGAGGGGTTTTTTG |
| 21-nt Linker | AACCTGGCGGCAGCGCAAAAG |
| Switch universal forward primer | GCCGGGTTTAGAAATCTGAAGCTCTAGAGAGCGCTAATACGACTCACTATAGGG |
| Switch universal reverse primer | TTTACGCATCTTTTGCGCTGCCGCCAGGTT |
| Trigger universal forward primer | GCCGGGTTTAGAAATCTGAAGCTCTAGAGAGCGCTAATACGACTCACTATAGGG |
| Trigger universal reverse primer | CCCGTTTAGAGGCCCCAAGGGGTTATGCTA |
| Switch backbone forward primer | AACCTGGCGGCAGCGCAA |
| Switch backbone reverse primer | TCTCTAGAGCTTCAGATTTCTAAACCCGGCCATAAGGGAGAGCGTCGAGATC |
| Trigger backbone forward primer | TAGCATAACCCCTTGGGGC |
| Trigger backbone reverse primer | TCTCTAGAGCTTCAGATTTCTAAACCCGGCCGAGATCTCGATCCTCTACGC |
| Non-cognate RNA used for library characterization | GGGUCUCACGCCCUCAGCUGGGCGUGAGAUGAGCCUCGUCUCCAGAUGAC<br>GAGGCAACGUAGGAUCUGACUGAUCCUACUAU |
| GFPmut3b-ASV | ATGCGTAAAGGAGAAGAACTTTTCACTGGAGTTGTCCCAATTCTTGTTGAATTAGA<br>TGGTGATGTTAATGGGCACAAATTTTCTGTCAGTGGAGAGGGTGAAGGTGATGCA<br>ACATACGGAAAACTTACCCTTAAATTTATTTGCACTACTGGAAAACTACCTGTTCC<br>GTGGCCAACACTTGTCACTACTTTCGGTTATGGTGTTCAATGCTTTGCGAGATAC<br>CCAGATCACATGAAACAGCATGACTTTTTCAAGAGTGCCATGCCCGAAGGTTAC<br>GTACAGGAAAGAACTATATTTTTCAAAGATGACGGGAACTACAAGACACGTGCTG<br>AAGTCAAGTTTGAAGGTGATACCCTTGTTAATAGAATCGAGTTAAAAGGTATTGAT<br>TTTAAAGAAGATGGAAACATTCTTGGACACAAATTGGAATACAACTATAACTCACA<br>CAATGTATACATCATGGCAGACAAACAAAAGAATGGAATCAAAGTTAACTTCAAA<br>ATTAGACACAACATTGAAGATGGAAGCGTTCAACTAGCAGACCATTATCAACAAA<br>ATACTCCGATTGGCGATGGCCCTGTCCTTTTACCAGACAACCATTACCTGTCCA<br>CACAATCTGCCCTTTCGAAAGATCCCAACGAAAAGAGAGACCACATGGTCCTTC<br>TTGAGTTTGTAACCGCTGCTGGGATTACACATGGCATGGATGAACTATACAAAAG<br>GCCTGCAGCAAACGACGAAAACTACGCTGCATCAGTTTAATAA |

**Table S2. Sequence and performance information for first-generation toehold repressors**

**NOTES:**
1. Trigger RNA sequences are listed up to the base immediately before the T7 terminator used to terminate transcription.
2. Switch RNA sequences are listed up to the 30th base (inclusive) following the end of their trigger binding domain. This 30-nt sequence contains the 21-nt linker sequence and the first 9-nts of GFPmut3b.

| First-generation toehold repressor ranking | First-generation toehold repressor index | GFP fold reduction | Switch sequence | Trigger sequence | Switch plasmid | Switch plasmid origin/Resistance | Trigger plasmid | Trigger plasmid origin/Resistance |
|---|---|---|---|---|---|---|---|---|
| 1 | 9 | 172.8 ± 29.4 | GGGAAAGUGAAGAAGAAUAUAAAGUUGAAUGAAGGCGAUGAUUGUAAGGUUACUACACUUACACUCAUCGCUUUCAUUCUACUUUAUAAACAGAGGAGAAUAAAGAUGAAUGAAAGCAACCUGGCGGCAGCGCAAAAGAUGCGUAAA | GGGUCAAGGCAUUAGCCCUAAUGCCUUGAAUACUUACAAUCAUCGCCUUCAUUCAACUUUAUAUUCUUCUUCACUUUACU | pAG_ToeRep_N09_switch | ColA/kanamycin | pAG_ToeRep_N09_trigger | CDF/spectinomycin |
| 2 | 40 | 93.3 ± 26.6 | GGGAAUAAAGAAAGAUGAAUAAAUGUGAAGAUAAGGUUGAUGAAUGUAUACUAUAAUUACAUUUAUCAACCUUAUCUUCUCAUUUAUAAACAGAGGAGAAUAAAUAUGAAGAUAAGGAACCUGGCGGCAGCGCAAAAGAUGCGUAAA | GGGAGUCGGUCUCGCGGUUGAGACCGACUAUUUACAUUCAUCAACCUUAUCUUCACAUUUAUUCAUCUUUCUUUAUUACU | pAG_ToeRep_N40_switch | ColA/kanamycin | pAG_ToeRep_N40_trigger | CDF/spectinomycin |
| 3 | 1 | 88.5 ± 17.4 | GGGAGAAAUGAAGAUAAGAUAGAUGUGAUGAAGUGCGAAGUUGUAAUAACAAAUAACUAUUACUACUUCGCUCUUCAUCUCAUCUAUAAACAGAGGAGAAUAGAUAUGAUGAAGAGCAACCUGGCGGCAGCGCAAAAGAUGCGUAAA | GGGUCAGACAAACCGGUAGGUUUGUCUGAAUAUAUUACAACUUCGCACUUCAUCACAUCUAUCUUAUCUUCAUUUCUAAC | pAG_ToeRep_N01_switch | ColA/kanamycin | pAG_ToeRep_N01_trigger | CDF/spectinomycin |
| 4 | 42 | 67.5 ± 7.5 | GGGAGAAUAAUAAAGAGAAUAGAGGUGAGAAGAACGGUGAAAGAAAUAUCAACAUAUUAUUUCUUUCACCGUUCUUCUCGCCUCUAUAAACAGAGGAGAAUAGAGAUGAGAAGAACGAACCUGGCGGCAGCGCAAAAGAUGCGUAAA | GGGCGAACUUGAAACAGGAUUCAAGUUCGCAUUAUUUCUUUCACCGUUCUUCUCACCUCUAUUCUCUUUAUUAUUCUACU | pAG_ToeRep_N42_switch | ColA/kanamycin | pAG_ToeRep_N42_trigger | CDF/spectinomycin |
| 5 | 36 | 57.2 ± 16.4 | GGGAGAAUAAUGAUGGAGAUAGAGUGGAAGAUGAGGAGAGUGAGUAGUAACAAAUACACUACUAACUCUCCUCAUCUUCUACUCUAUAAACAGAGGAGAAUAGAGAUGAAGAUGAGGAACCUGGCGGCAGCGCAAAAGAUGCGUAAA | GGGCCUUGACUCGAUGCAACGAGUCAAGGCAAACUACUCACUCUCCUCAUCUUCCACUCUAUCUCCAUCAUUAUUCUACU | pAG_ToeRep_N36_switch | ColA/kanamycin | pAG_ToeRep_N36_trigger | CDF/spectinomycin |
| 6 | 25 | 37.5 ± 6.9 | GGGAGAAAUGAAAGAUGAAUAAGAAAUGAGAGAACGGGAAUAAUAAAGAACUUACUACUUUAUAAUUCCCGUUCUCUCACUUCUUAUAAACAGAGGAGAAUAAGAAUGGAGAGAACGAACCUGGCGGCAGCGCAAAAGAUGCGUAAA | GGGUGGAGUACGAGCGAGUUCGUACUCCACGACUUUAUUAUUCCCGUUCUCUCAUUUCUUAUUCAUCUUUCAUUUCUCUU | pAG_ToeRep_N25_switch | ColA/kanamycin | pAG_ToeRep_N25_trigger | CDF/spectinomycin |
| 7 | 35 | 24.7 ± 0.5 | GGGAAGAUGAUAGAGAGAAUAGAAUGUAAGAUGUGGUGCGCGUUAUACAAUGAAUAUAACGAGCGCACCUCAUCUUAUAUUCUAUAAACAGAGGAGAAUAGAAAUGAAGAUGAGGAACCUGGCGGCAGCGCAAAAGAUGCGUAAA | GGGUCAGUAGGGACUUGAUUCCCUACUGAUAAAUAACGCGCGCACCACAUCUUACAUUCUAUUCUCUCUAUCAUCUUAAU | pAG_ToeRep_N35_switch | ColA/kanamycin | pAG_ToeRep_N35_trigger | CDF/spectinomycin |
| 8 | 6 | 19.9 ± 0.8 | GGGAAAGAUGGAGAAGAUAUAGAUGGGAGGGAUUGGGAGGUGUGUGUAAGAAACAGAUACACAAACCUCCCUAUCCCUCCCAUCUAUAAACAGAGGAGAAUAGAUAUGAGGGAUAGGAACCUGGCGGCAGCGCAAAAGAUGCGUAAA | GGGCGGCAGAUGCUAGUCAGCAUCUGCCGCAAUACACACACCUCCCAAUCCCUCCCAUCUAUAUCUUCUCCAUCUUUAAU | pAG_ToeRep_N06_switch | ColA/kanamycin | pAG_ToeRep_N06_trigger | CDF/spectinomycin |
| 9 | 21 | 17.3 ± 6.0 | GGGAAAGUAAAGAUAAGAAUAAAUGAUGAAUGGCGCUGUAUGUGUUGUACGGAAGAGACAACAGAUACAGCUCCAUUCACCAUUUAUAAACAGAGGAGAAUAAAUAUGGAAUGGAGCAACCUGGCGGCAGCGCAAAAGAUGCGUAAA | GGGUCUGGGAUGCUUGGUAGCAUCCCAGAGAAACAACACAUACAGCGCCAUUCAUCAUUUAUUCUUAUCUUUACUUUAUU | pAG_ToeRep_N21_switch | ColA/kanamycin | pAG_ToeRep_N21_trigger | CDF/spectinomycin |
| 10 | 3 | 16.8 ± 1.4 | GGGAGUGAAAGAGUGAAGAUAAGAGGUGAGAGUUGGGAUGUGUGUGAGACACAAGAACUCACAAACAUCCCUACUCUCAGCUCUUAUAAACAGAGGAGAAUAAGAAUGGAGAGUAGGAACCUGGCGGCAGCGCAAAAGAUGCGUAAA | GGGCGGCCAGGUCGAAGUGGACCUGGCCGCGACUCACACACAUCCCAACUCUCACCUCUUAUCUUCACUCUUUCACUUUA | pAG_ToeRep_N03_switch | ColA/kanamycin | pAG_ToeRep_N03_trigger | CDF/spectinomycin |
| 11 | 13 | 15.8 ± 12.8 | GGGAAGAUAAGAAUGAAUAUAAAGAAUGAUGAAACGUGUAAGAAUAGUCAUAAACAAACUAUUAUUACACGUUUCAUCACUCUUUAUAAACAGAGGAGAAUAAAGAUGGAUGAAACGAACCUGGCGGCAGCGCAAAAGAUGCGUAAA | GGGAGGUGCGAAUCUCAGAAUUCGCACCUAUUACUAUUCUUACACGUUUCAUCAUUCUUUAUAUUCAUUCUUAUCUUAGC | pAG_ToeRep_N13_switch | ColA/kanamycin | pAG_ToeRep_N13_trigger | CDF/spectinomycin |
| 12 | 34 | 15.6 ± 1.1 | GGGAUAAUGGAAGGAGGUAUAAAGGGUGGAGAGUGCUGUGUGUUGAGUGACAAUAAUACUCAAGACACAGCUCUCUCCACCCUUUAUAAACAGAGGAGAAUAAAGAUGGGAGAGAGCAACCUGGCGGCAGCGCAAAAGAUGCGUAAA | GGGCCACGUUAUGAAUCGCCAUAACGUGGCAAACUCAACACACAGCACUCUCCACCCUUUAUACCUCCUUCCAUUAUCUU | pAG_ToeRep_N34_switch | ColA/kanamycin | pAG_ToeRep_N34_trigger | CDF/spectinomycin |
| 13 | 26 | 15.2 ± 3.4 | GGGAAAUGGAGAGAAGUGAUAGAAGAGAAUGAGGCGAUGAAAGUAGAUCCAACCUUUAUCUACAUUCAUCGUCUCAUUCCCUUCUAUAAACAGAGGAGAAUAGAAAUGAAUGAGACGAACCUGGCGGCAGCGCAAAAGAUGCGUAAA | GGGUCCGGUUAAACGCCGAUUUAACCGGACAUAUCUACUUUCAUCGCCUCAUUCUCUUCUAUCACUUCUCUCCAUUUACU | pAG_ToeRep_N26_switch | ColA/kanamycin | pAG_ToeRep_N26_trigger | CDF/spectinomycin |
| 14 | 20 | 14.4 ± 1.3 | GGGAGAAGUGAGGAGUGGAUAAGAGGGAGUAAGUGCGGUGAGUGUAGUCACACAAUAACUACAAUCACCGCUCUUACUCUCUCUUAUAAACAGAGGAGAAUAAGAAUGAGUAAGAGCAACCUGGCGGCAGCGCAAAAGAUGCGUAAA | GGGCCUGUGUAGAAUACCCUCUACACAGGCAUACUACACUCACCGCACUUACUCCCUCUUAUCCACUCCUCACUUCUAUA | pAG_ToeRep_N20_switch | ColA/kanamycin | pAG_ToeRep_N20_trigger | CDF/spectinomycin |
| 15 | 27 | 13.7 ± 2.4 | GGGAUGGAGAGAGAUGAGAUAAAGUGUAGAGAGGCGUUGUUGAGUAGUAAUAAGAGAACUACUAAACAACGUCUCUCUAAACUUUAUAAACAGAGGAGAAUAAAGAUGAGAGAGACGAACCUGGCGGCAGCGCAAAAGAUGCGUAAA | GGGUCCGAUAGCUCAGAUUAGCUAUCGGACAAACUACUCAACAACGCCUCUCUACACUUUAUCUCAUCUCUCUCCAUAUU | pAG_ToeRep_N27_switch | ColA/kanamycin | pAG_ToeRep_N27_trigger | CDF/spectinomycin |
| 16 | 33 | 12.3 ± 1.4 | GGGAGAAAGUGGAGAAGUAAUGAAGGGAAGAUGAGGAGAGUGAGUAAAGGACAUACAUUUACUAACUCUCCUCAUCUUCCCUUCAUUAAACAGAGGAGAAAUGAAAUGAAGAUGAGGAACCUGGCGGCAGCGCAAAAGAUGCGUAAA | GGGCGUGGUUCAGGUUGGCCUGAACCACGCAAUUUACUCACUCUCCUCAUCUUCCCUUCAUUACUUCUCCACUUUCUAUA | pAG_ToeRep_N33_switch | ColA/kanamycin | pAG_ToeRep_N33_trigger | CDF/spectinomycin |
| 17 | 8 | 11.9 ± 1.2 | GGGAGAAUGAUAGAUGAAAUAAUAGGUAAAGAGUCGAGUUGUUGUAAUGAAACCUGGAUUACAACAACUCGUCUCUUUAACUAUUAUAAACAGAGGAGAAUAAUAAUGAAAGAGACGAACCUGGCGGCAGCGCAAAAGAUGCGUAAA | GGGAGUGACCGGUGUGGCAACCGGUCACUAAUAUUACAACAACUCGACUCUUUACCUAUUAUUUCAUCUAUCAUUCUUAU | pAG_ToeRep_N08_switch | ColA/kanamycin | pAG_ToeRep_N08_trigger | CDF/spectinomycin |
| 18 | 28 | 11.4 ± 0.4 | GGGAUGAAGAUAAGAAAGAUAAAGGGUGAGAAAUGGAGAGAGAUAAGUUCAAAUUUCACUUAUAUCUCUCCAUUUCUCAACCUUUAUAAACAGAGGAGAAUAAAGAUGGAGAAAUGGAACCUGGCGGCAGCGCAAAAGAUGCGUAAA | GGGCGAGUGCCUUUAGACAAAGGCACUCGCAUACUUAUCUCUCUCCAUUUCUCACCCUUUAUCUUUCUUAUCUUCAUCUU | pAG_ToeRep_N28_switch | ColA/kanamycin | pAG_ToeRep_N28_trigger | CDF/spectinomycin |
| 19 | 37 | 11.2 ± 2.2 | GGGUGGAAGAUGGAGGAGAUUGAAGAGGAAUGGAGGUGGUAUGUAUGUACGAACCAAACAUACUUACCACCUCCAUUCCUCUUCAAUAAACAGAGGAGAAUUGAAAUGGAAUGGAGGAACCUGGCGGCAGCGCAAAAGAUGCGUAAA | GGGUCUAUCCCUCGCGAAUGAGGGAUAGACAAACAUACAUACCACCUCCAUUCCUCUUCAAUCUCCUCCAUCUUCCAAUU | pAG_ToeRep_N37_switch | ColA/kanamycin | pAG_ToeRep_N37_trigger | CDF/spectinomycin |
| 20 | 5 | 10.5 ± 0.8 | GGGAGAAGUGGAAGAGUGAUAAGAGGUGAGAUGUGGUUGAGGAGUUGUAACGACUCGACAACUACUCAACCUCAUCUCAGCUCUUAUAAACAGAGGAGAAUAAGAAUGGAGAUGAGGAACCUGGCGGCAGCGCAAAAGAUGCGUAAA | GGGCGCAAUAUACUGUCAGGUAUAUUGCGCAAACAACUCCUCAACCACAUCUCACCUCUUAUCACUCUUCCACUUCUUAU | pAG_ToeRep_N05_switch | ColA/kanamycin | pAG_ToeRep_N05_trigger | CDF/spectinomycin |
| 21 | 22 | 10.0 ± 1.3 | GGGAGAAGUAAGAGAGUGAUUAGAUGUGUGAUGUCGGUUGUGCUUAUUAACGAAACAAAUAAGAACAACCGUCAUCACAUAUCUAAUAAACAGAGGAGAAUUAGAAUGGUGAUGACGAACCUGGCGGCAGCGCAAAAGAUGCGUAAA | GGGCCCUUCUGGGCAUGGUCCCAGAAGGGAAUAAUAAGCACAACCGACAUCACACAUCUAAUCACUCUCUUACUUCUAUA | pAG_ToeRep_N22_switch | ColA/kanamycin | pAG_ToeRep_N22_trigger | CDF/spectinomycin |
| 22 | 41 | 9.6 ± 0.2 | GGGAAAGUGAAGAAGAUGAUAAGAAGUGAAGAAGGCUGUGUAAAGUAUCCUACUAUCAUACUUAACACAGCUUUCUUCACUUCUUAUAAACAGAGGAGAAUAAGAAUGGAAGAAAGCAACCUGGCGGCAGCGCAAAAGAUGCGUAAA | GGGUCUGGACGUAUGUUUGUACGUCCAGAAUUAUACUUUACACAGCCUUCUUCACUUCUUAUCAUCUUCUUCACUUUACU | pAG_ToeRep_N41_switch | ColA/kanamycin | pAG_ToeRep_N41_trigger | CDF/spectinomycin |
| 23 | 44 | 9.5 ± 0.5 | GGGAUAAGAGAAGUGGAGAUAAAGGGUGAAAGUGGGAGGGUGAGAGAACAUACAUCAUUCUCUUACCCUCCUACUUUCACCCUUUAUAAACAGAGGAGAAUAAAGAUGGAAAGUAGGAACCUGGCGGCAGCGCAAAAGAUGCGUAAA | GGGCGCUCAGAACCCGUCCGUUCUGAGCGCAUUUCUCUCACCCUCCCACUUUCACCCUUUAUCUCCACUUCUCUUAUAUU | pAG_ToeRep_N44_switch | ColA/kanamycin | pAG_ToeRep_N44_trigger | CDF/spectinomycin |
| 24 | 11 | 9.2 ± 0.6 | GGGAGGAAUGGAGAGAAUAUAGAAGGUGAGGGCUGGUUUAGGUUGAUUCAGAACAAUAAUCAAACUAAACCUGCCCUCAGCUUCUAUAAACAGAGGAGAAUAGAAAUGGAGGGCAGGAACCUGGCGGCAGCGCAAAAGAUGCGUAAA | GGGACACUAGCCGUUCGAUCGGCUAGUGUCGAAAUCAACCUAAACCAGCCCUCACCUUCUAUAUUCUCUCCAUUCCUUAU | pAG_ToeRep_N11_switch | ColA/kanamycin | pAG_ToeRep_N11_trigger | CDF/spectinomycin |
| 25 | 23 | 8.6 ± 4.6 | GGGCGUGAGCGGAUGAAUAUAGAUGUGAUAAAGUGGUGUGAUGUAGUAACGACGACAUACUACAUCACACCUCUUUAUCUCAUCUAUAAACAGAGGAGAAUAGAUAUGAUAAAGAGGAACCUGGCGGCAGCGCAAAAGAUGCGUAAA | GGGAGUGUACAACAGGUGCGUUGUACACUAAAUACUACAUCACACCACUUUAUCACAUCUAUAUUCAUCCGCUCACGAAC | pAG_ToeRep_N23_switch | ColA/kanamycin | pAG_ToeRep_N23_trigger | CDF/spectinomycin |

| | | | | | | | | |
|---|---|---|---|---|---|---|---|---|
| 26 | 4 | 8.5 ± 1.9 | GGGAAUGGAGAGAGUGGGAUAGAGUGUGAGGAGUGCGAAGGAAGUGUAC<br>ACAAGACAUACACUACCUUCGCUCUCCUCAGACUCUAUAAACAGAGGAG<br>AAUAGAGAUGGAGGAGAGCAACCUGGCGGCAGCGCAAAAGAUGCGUAAA | GGGCGAGAGGUUUCAGGACAAACCUCUC<br>GCAUUACACUUCCUUCGCACUCCUCACA<br>CUCUAUCCCACUCUCUCCAUUAAU | pAG_ToeRep_N04_switch | ColA/kanamycin | pAG_ToeRep_N04_trigger | CDF/spectinomycin |
| 27 | 10 | 7.4 ± 0.5 | GGGUGGUGGAGGAAGUGGAAUGAAUGGAAGGAGUGCGAGGUGAGUGUAG<br>AAAGAAGAUACACUAACCUCGCUCUCCUUCUAUUCAUUAAACAGAGGAGA<br>AAUGAAAUGAAGGAGAGCAACCUGGCGGCAGCGCAAAAGAUGCGUAAA | GGGCCUAAGGGCCUGGAGCGGCCCUUA<br>GGCAUUACACUCACCUCGCACUCCUUC<br>CAUUCAUUCCACUUCCUCCACCAUAU | pAG_ToeRep_N10_switch | ColA/kanamycin | pAG_ToeRep_N10_trigger | CDF/spectinomycin |
| 28 | 39 | 7.4 ± 0.8 | GGGAGGAGAAAUGAAAGAAUAGAAGGUAAGAAGGCGGUGUGGAUUGUCU<br>UAAUAACUGACAAUACACACCGUCUUCUUACCUUCUAUAAACAGAGGAGA<br>AUAGAAAUGAAGAAGACGAACCUGGCGGCAGCGCAAAAGAUGCGUAAA | GGGACAUGCGAAGGCGGAACUUCGCAUG<br>UCACGACAAUCCACACCGCCUUCUUACC<br>UUCUAUUCUUUCAUUUCUCCUUAU | pAG_ToeRep_N39_switch | ColA/kanamycin | pAG_ToeRep_N39_trigger | CDF/spectinomycin |
| 29 | 32 | 6.6 ± 0.5 | GGGAAUGGAGAGAAUGAAAUACAGAGUGAGAUGAGGAGAGUGAGAGUAAC<br>UACAACAUACUCUAACUCUCCUCAUCUCAUUCUGUAUAAACAGAGGAGA<br>AUACAGAUGGAGAUGAGGAACCUGGCGGCAGCGCAAAAGAUGCGUAAA | GGGCCCUUAUGGCAACGAAGCCAUAAGG<br>GCAUUACUCUCACUCUCCUCAUCUCAC<br>UCUGUAUUUCAUUCUCUCCAUUUAU | pAG_ToeRep_N32_switch | ColA/kanamycin | pAG_ToeRep_N32_trigger | CDF/spectinomycin |
| 30 | 14 | 6.2 ± 3.1 | GGGAAGAUGGAGAGAUGGAUAAGAGAGAAAUGGUGCGAGAGAAGAGUUAU<br>UAUCUUCAACUCUACUCUCGCUCCAUUUCUCUCUUAUAAACAGAGGAGA<br>AUAAGAAUGAAAUGGAGCAACCUGGCGGCAGCGCAAAAGAUGCGUAAA | GGGCCGUUGUCGGGCGAGUCCGACAAC<br>GGCAUAACUCUUCUCUCGCACCAUUUC<br>UCUCUUAUCCAUCUCUCCAUCUUUAU | pAG_ToeRep_N14_switch | ColA/kanamycin | pAG_ToeRep_N14_trigger | CDF/spectinomycin |
| 31 | 18 | 5.9 ± 1.3 | GGGAGAAGUUACGCCGAAAUAGAAGAUAGAAUAAGGUUGAUGUGGCUGAA<br>ACAAAUACAGCCAAAUCAACCUUAUUCUAGCUUCUAUAAACAGAGGAGAA<br>UAGAAAUGAGAAUAAGGAACCUGGCGGCAGCGCAAAAGAUGCGUAAA | GGGUCCUUGGUUCGUUGUAGAACCAAG<br>GAAAGCAGCCACAUCAACCUUAUUCUAU<br>CUUCUAUUUCGGCGUAACUUCUAUA | pAG_ToeRep_N18_switch | ColA/kanamycin | pAG_ToeRep_N18_trigger | CDF/spectinomycin |
| 32 | 29 | 5.4 ± 0.1 | GGGAAAGUGAGAGGAAGUAUAGAUGGUGGAGGAGGCUGUGUGUGUAAUA<br>CUCUAAACAUUACAAACACAGCUUCCUCCACCAUCUAUAAACAGAGGAGA<br>AUAGAUAUGGGAGGAAGCAACCUGGCGGCAGCGCAAAAGAUGCGUAAA | GGGUCUGUGUUCCGAGUCUGGAACACA<br>GACAAAUUACACACACAGCCUCCUCCAC<br>CAUCUAUACUUCCUCUCACUUUAUU | pAG_ToeRep_N29_switch | ColA/kanamycin | pAG_ToeRep_N29_trigger | CDF/spectinomycin |
| 33 | 7 | 5.3 ± 0.8 | GGGAUUGGCCCGCGGAAUAUAAAGUUGAAGAGCUGCCUUUGUUUCUUUA<br>AACAAAUGAAAGAAACAAAGGCUGCUCUUCCACUUUAUAAACAGAGGAGA<br>AUAAAGAUGAAGAGCAGCAACCUGGCGGCAGCGCAAAAGAUGCGUAAA | GGGUGAUGGAACUACACAAAGUUCCAUC<br>AAAUAAAGAAACAAAGGCAGCUCUUCAAC<br>UUUAUAUUCCGCGGGCCAAUAAU | pAG_ToeRep_N07_switch | ColA/kanamycin | pAG_ToeRep_N07_trigger | CDF/spectinomycin |
| 34 | 12 | 5.3 ± 8.1 | GGGGAUGAAGAAGAUGAUAAUGAAGAGAAGAUGAGGUGUAUAUGUUUGGA<br>GGAAAUACAAACACAUACACCUCAUCUUCCCUUCAUUAAACAGAGGAGAA<br>AUGAAAUGAAGAUGAGGAACCUGGCGGCAGCGCAAAAGAUGCGUAAA | GGGCGCCAACUCCCAUUUCGGAGUUGG<br>CGAAACAAACAUAUACACCUCAUCUUCU<br>CUUCAUUAUCAUCUUCUUCAUCAUU | pAG_ToeRep_N12_switch | ColA/kanamycin | pAG_ToeRep_N12_trigger | CDF/spectinomycin |
| 35 | 38 | 5.0 ± 3.4 | GGGAAAUGGAGGGAGUGGAUAAGGUGGGUGGAGAGGAGUAAAGAAGUAC<br>UCUUCUCAUACUUCAUUACUCCUCUCCACCUACCUUAUAAACAGAGGAG<br>AAUAAGGAUGGUGGAGAGGAACCUGGCGGCAGCGCAAAAGAUGCGUAAA | GGGCGCGCAUCCCAGAGAUGGGAUGCG<br>CGCAUUACUUCUUUACUCCUCUCCACC<br>CACCUUAUCCACUCCCUCCAUUUACU | pAG_ToeRep_N38_switch | ColA/kanamycin | pAG_ToeRep_N38_trigger | CDF/spectinomycin |
| 36 | 2 | 4.8 ± 0.9 | GGGAGAGUAAGAAAGAUGAUAAAGGAAAGUAAGUGCUGUGUGUGUAGUAA<br>CGAAGAAACUACAAACACAGCUCUUACUUACCUUUAUAAACAGAGGAGAA<br>UAAAGAUGAGUAAGAGCAACCUGGCGGCAGCGCAAAAGAUGCGUAAA | GGGCGCAAAUCGUCGAGGAACGAUUUGC<br>GAAUACUACACACACAGCACUUACUUUC<br>CUUUAUCAUCUUUCUUACUCUUAU | pAG_ToeRep_N02_switch | ColA/kanamycin | pAG_ToeRep_N02_trigger | CDF/spectinomycin |
| 37 | 43 | 3.3 ± 0.2 | GGGAGAAGAAGUGAUGAAAUAAAGUGGAUAAAGAGGUGUAGUAAUAAGAC<br>AAACAAGCUUAUUACUACACCUCUUUAUCUACUUUAUAAACAGAGGAGAA<br>UAAAGAUGAUAAAGAGGAACCUGGCGGCAGCGCAAAAGAUGCGUAAA | GGGUCCAAAUUAUAGGUGAAUAAUUUGG<br>ACGACUUAUUACUACACCUCUUUAUCCA<br>CUUUAUUUCAUCACUUCUUCUUUA | pAG_ToeRep_N43_switch | ColA/kanamycin | pAG_ToeRep_N43_trigger | CDF/spectinomycin |
| 38 | 17 | 3.1 ± 0.8 | GGGAAUGGAGAGAAACUUAUAGAAUGUGUGAAGCGCUGUGUUAUUGUAC<br>AAGAGAAAUACAAUGACACAGCUCUUCACAAAUUCUAUAAACAGAGGAGA<br>AUAGAAAUGGUGAAGAGCAACCUGGCGGCAGCGCAAAAGAUGCGUAAA | GGGACAGUGGCAGAGGACCCUGCCACUG<br>UAAAUACAAUAACACAGCGCUUCACACA<br>UUCUAUAAGUUUCUCUCCAUUAAU | pAG_ToeRep_N17_switch | ColA/kanamycin | pAG_ToeRep_N17_trigger | CDF/spectinomycin |
| 39 | 30 | 2.5 ± 0.4 | GGGAGAGUAAAGAGAAGUAAUGAUGAGAGGGAUGGGAGAAGAGUGUAUGA<br>CAGAUACAUACACACUUCUCCUAUCCCUCCCAUCAUUAAACAGAGGAGA<br>AAUGAUAUGAGGGAUAGGAACCUGGCGGCAGCGCAAAAGAUGCGUAAA | GGGCCAAGCGGAGCGUCUUCUCCGCUU<br>GGCAUAUACACUCUUCUCCCAUCCCUC<br>UCAUCAUUACUUCUCUUUACUCUUAU | pAG_ToeRep_N30_switch | ColA/kanamycin | pAG_ToeRep_N30_trigger | CDF/spectinomycin |
| 40 | 31 | 2.2 ± 0.2 | GGGAGAAAUGGAGAAGAUAAUGGAGGAGAGAUGUGGUGUAAGGUUGUGA<br>AUACAGAUCACAACAUUACACCUCAUCUCUCCUCCAUUAAACAGAGGAGA<br>AAUGGAAUGGAGAUGAGGAACCUGGCGGCAGCGCAAAAGAUGCGUAAA | GGGCGUCGGUCCGUGCCAGCGGACCGA<br>CGCGACACAACCUUACACCACAUCUCUC<br>CUCCAUUAUCUUCUCCAUUUCUUAU | pAG_ToeRep_N31_switch | ColA/kanamycin | pAG_ToeRep_N31_trigger | CDF/spectinomycin |
| 41 | 19 | 1.9 ± 0.1 | GGGAGGGAAGUGGGAGUGAUAAGAGGUGAAGAGUGCUGGUUGAUGUGUG<br>AGACAAGAACACAUAAACCAGCUCUCUUCACCUCUUAUAAACAGAGGAGA<br>AUAAGAAUGGAAGAGAGCAACCUGGCGGCAGCGCAAAAGAUGCGUAAA | GGGCCGUGCACAACGUGGUUUGUGCAC<br>GGCAAACACAUCAACCAGCACUCUUCAC<br>CUCUUAUCACUCCCACUUCCCUAUA | pAG_ToeRep_N19_switch | ColA/kanamycin | pAG_ToeRep_N19_trigger | CDF/spectinomycin |
| 42 | 15 | 1.5 ± 1.3 | GGGAAGAUGAAGAUAAGAAUAAAUGAGAAAGACGCGGAGAGAAUAAGUAU<br>CAAAUCAACUUAUACUCUCCGUGUCUUUCCCAUUUAUAAACAGAGGAGA<br>AUAAAUAUGAAAGACACGAACCUGGCGGCAGCGCAAAAGAUGCGUAAA | GGGUGAUUUCAAGCUAGUUCUUGAAAUC<br>ACAUACUUAUUCUCUCCGCGUCUUUCU<br>CAUUUAUUCUUAUCUUCAUCUUUAU | pAG_ToeRep_N15_switch | ColA/kanamycin | pAG_ToeRep_N15_trigger | CDF/spectinomycin |
| 43 | 16 | 1.3 ± 0.7 | GGGAGUGUGGAGAAGUGGAUAAUGGAUGAAGAGUGCUGUGUGUAUGUAA<br>CUCAUUUAUACAUACACACAGCUCUCUUCAGCCAUUAUAAACAGAGGAG<br>AAUAAUGAUGGAAGAGAGCAACCUGGCGGCAGCGCAAAAGAUGCGUAAA | GGGUCGCCCGGAGCUCUCGCUCCGGGC<br>GACAUUACAUACACACAGCACUCUUCAU<br>CCAUUAUCCACUUCUCCACACUAUA | pAG_ToeRep_N16_switch | ColA/kanamycin | pAG_ToeRep_N16_trigger | CDF/spectinomycin |
| 44 | 24 | 0.1 ± 0.1 | GGGAUGGAAUGGUGGUGGAAUGAAGAGAAAUGGAGGAUGAAAGAGUGUAA<br>CUUACCAACACUCAUUCAUCCUCCAUUUCCCUUCAUUAAACAGAGGAGA<br>AAUGAAAUGAAAUGGAGGAACCUGGCGGCAGCGCAAAAGAUGCGUAAA | GGGCGGACUACCAAGGGUUUGGUAGUC<br>CGCAUACACUCUUUCAUCCUCCAUUUC<br>UCUUCAUUCCACCACCAUUCCAUUAU | pAG_ToeRep_N24_switch | ColA/kanamycin | pAG_ToeRep_N24_trigger | CDF/spectinomycin |

**Table S3. Sequence and performance information for three-way junction (3WJ) repressors**

**NOTES:**
1. Trigger RNA sequences are listed up to the base immediately before the T7 terminator used to terminate transcription.
2. Switch RNA sequences are listed up to the 30th base (inclusive) following the end of their trigger binding domain. This 30-nt sequence contains the 21-nt linker sequence and the first 9-nts of GFPmut3b.

| 3WJ repressor ranking | 3WJ repressor index | GFP fold reduction | Switch sequence | Trigger sequence | Switch plasmid | Switch plasmid origin/Resistance | Trigger plasmid | Trigger plasmid origin/Resistance |
|---|---|---|---|---|---|---|---|---|
| 1 | 20 | 134.2 ± 8.1 | GGGAUGAAUGAUAUACACUUGUUAUAGUUAUGAACA<br>GAGGAGACAUAACAUGAACAAGCACGAAUUGACUAC<br>ACUAAACCUGGCGGCAGCGCAAAAGAUGCGUAAA | GGGACGAAUUGAUUUGUCAAUUCGU<br>GCGUGUAUAUCAUUCAUCAU | pYZ_3WJrep_N20_switch | ColA/kanamycin | pYZ_3WJrep_N20_trigger | ColE1/ampicillin |
| 2 | 19 | 85.2 ± 4.9 | GGGACUAAUCAGAUCUACUUGUUAUAGUUAUGAACA<br>GAGGAGACAUAACAUGAACAAGCACCUAACAAGACU<br>AAUCAACCUGGCGGCAGCGCAAAAGAUGCGUAAA | GGGACCUAACAUAAACUUGUUAGGU<br>GCGUAGAUCUGAUUAGUGUG | pYZ_3WJrep_N19_switch | ColA/kanamycin | pYZ_3WJrep_N19_trigger | ColE1/ampicillin |
| 3 | 10 | 66.1 ± 3.2 | GGGUAGUAAGAUUAGAUAUUGUUAUAGUUAUGAACA<br>GAGGAGACAUAACAUGAACAAGUACAAGAACCGAAAC<br>GAGAACCUGGCGGCAGCGCAAAAGAUGCGUAAA | GGGACAAGAACAAUACGGUUCUUGU<br>ACUAUCUAAUCUUACUAAAC | pYZ_3WJrep_N10_switch | ColA/kanamycin | pYZ_3WJrep_N10_trigger | ColE1/ampicillin |
| 4 | 24 | 56.9 ± 7.0 | GGGCUCCUAUCACUUUACUUGUUAUAGUUAUGAAC<br>AGAGGAGACAUAACAUGAACAAGCACACUAACUACAA<br>AUUCAACCUGGCGGCAGCGCAAAAGAUGCGUAAA | GGGACACUAACCAUAUAGUUAGUGU<br>GCGUAAAGUGAUAGGAGUAA | pYZ_3WJrep_N24_switch | ColA/kanamycin | pYZ_3WJrep_N24_trigger | ColE1/ampicillin |
| 5 | 15 | 44.7 ± 7.1 | GGGUCCAAAUCUUACUUAUUGUUAUAGUUAUGAACA<br>GAGGAGACAUAACAUGAACAAUCACUACCUCGAAUA<br>CUUAAACCUGGCGGCAGCGCAAAAGAUGCGUAAA | GGGACUACCUCACACUCGAGGUAGU<br>GAUAAGUAAGAUUUGGAAGU | pYZ_3WJrep_N15_switch | ColA/kanamycin | pYZ_3WJrep_N15_trigger | ColE1/ampicillin |
| 6 | 32 | 41.8 ± 1.1 | GGGUACUCAAGAAUUCCAUUGUUAUAGUUAUGAACA<br>GAGGAGACAUAACAUGAACAAUCACAUACUAACAUU<br>ACUAAACCUGGCGGCAGCGCAAAAGAUGCGUAAA | GGGACAUACUAUAUCGUUAGUAUGU<br>GAUGGAAUUCUUGAGUAAUG | pYZ_3WJrep_N32_switch | ColA/kanamycin | pYZ_3WJrep_N32_trigger | ColE1/ampicillin |
| 7 | 21 | 41.2 ± 1.8 | GGGACUACUAUUCACUUAUUGUUAUAGUUAUGAACA<br>GAGGAGACAUAACAUGAACAAUCCGACACUACAAUU<br>UCGAAACCUGGCGGCAGCGCAAAAGAUGCGUAAA | GGGCGACACUAAUAGUGUAGUGUCG<br>GAUAAGUGAAUAGUAGUAGA | pYZ_3WJrep_N21_switch | ColA/kanamycin | pYZ_3WJrep_N21_trigger | ColE1/ampicillin |
| 8 | 13 | 41.0 ± 5.4 | GGGACAAUCAAAUACAAAUUGUUAUAGUUAUGAACA<br>GAGGAGACAUAACAUGAACAAUCACAUACAAAGCAAA<br>CGAAACCUGGCGGCAGCGCAAAAGAUGCGUAAA | GGGACAUACAAUGAACUUUGUAUGU<br>GAUUUGUAUUUGAUUGUAGC | pYZ_3WJrep_N13_switch | ColA/kanamycin | pYZ_3WJrep_N13_trigger | ColE1/ampicillin |
| 9 | 2 | 40.3 ± 2.8 | GGGAUGAUAUUGAAUUACUUGUUAUAGUUAUGAACA<br>GAGGAGACAUAACAUGAACAAGCACACUACACUAAA<br>CAUAAACCUGGCGGCAGCGCAAAAGAUGCGUAAA | GGGACACUACAACUCAGUGUAGUGU<br>GCGUAAUUCAAUAUCAUACU | pYZ_3WJrep_N02_switch | ColA/kanamycin | pYZ_3WJrep_N02_trigger | ColE1/ampicillin |
| 10 | 3 | 37.9 ± 2.4 | GGGAGUAUAAGAUAGAUAUUGUUAUAGUUAUGAACA<br>GAGGAGACAUAACAUGAACAAGUACGAAGCAACGAA<br>CAUAAACCUGGCGGCAGCGCAAAAGAUGCGUAAA | GGGACGAAGCAUAAAGUUGCUUCGU<br>ACUAUCUAUCUUAUACUAAC | pYZ_3WJrep_N03_switch | ColA/kanamycin | pYZ_3WJrep_N03_trigger | ColE1/ampicillin |
| 11 | 16 | 37.7 ± 12.7 | GGGUCCCAUUAUCUUACCUUGUUAUAGUUAUGAAC<br>AGAGGAGACAUAACAUGAACAAGCCUACACUACCAU<br>UCCCGAACCUGGCGGCAGCGCAAAAGAUGCGUAAA | GGGCUACACUAAACAGGUAGUGUAG<br>GCGGUAAGAUAAUGGGAGUA | pYZ_3WJrep_N16_switch | ColA/kanamycin | pYZ_3WJrep_N16_trigger | ColE1/ampicillin |
| 12 | 23 | 37.2 ± 6.2 | GGGCAAAUACUCCAUAUCUUGUUAUAGUUAUGAACA<br>GAGGAGACAUAACAUGAACAAGCACAUAAACUCACU<br>GAUAAACCUGGCGGCAGCGCAAAAGAUGCGUAAA | GGGACAUAAACCUAUGAGUUUAUGU<br>GCGAUAUGGAGUAUUUGAAA | pYZ_3WJrep_N23_switch | ColA/kanamycin | pYZ_3WJrep_N23_trigger | ColE1/ampicillin |
| 13 | 34 | 34.5 ± 2.2 | GGGCAAAGUAUCCAUCAUUUGUUAUAGUUAUGAACA<br>GAGGAGACAUAACAUGAACAAUCACAUACUAGAAUC<br>UAAGAACCUGGCGGCAGCGCAAAAGAUGCGUAAA | GGGACAUACUAACAAUCUAGUAUGU<br>GAAUGAUGGAUACUUUGAAA | pYZ_3WJrep_N34_switch | ColA/kanamycin | pYZ_3WJrep_N34_trigger | ColE1/ampicillin |
| 14 | 7 | 26.2 ± 1.0 | GGGCAAGAUUUAGUAGAUUUGUUAUAGUUAUGAACA<br>GAGGAGACAUAACAUGAACAAUCCGAACGACCGAAA<br>UGAUAACCUGGCGGCAGCGCAAAAGAUGCGUAAA | GGGCGAACGACGAAACGGUCGUUCG<br>GAAUCUACUAAAUCUUGAAA | pYZ_3WJrep_N07_switch | ColA/kanamycin | pYZ_3WJrep_N07_trigger | ColE1/ampicillin |
| 15 | 40 | 23.4 ± 1.8 | GGGAUACUUUCAAACUUAUUGUUAUAGUUAUGAACA<br>GAGGAGACAUAACAUGAACAAUCACACUAAUCGACU<br>GAAUAACCUGGCGGCAGCGCAAAAGAUGCGUAAA | GGGACACUAAUACUACGAUUAGUGU<br>GAUAAGUUUGAAAGUAUAAG | pYZ_3WJrep_N40_switch | ColA/kanamycin | pYZ_3WJrep_N40_trigger | ColE1/ampicillin |
| 16 | 11 | 23.2 ± 3.9 | GGGAUCAAUCAAUUCUACUUGUUAUAGUUAUGAACA<br>GAGGAGACAUAACAUGAACAAUCACAUAAACCUAAGA<br>ACUAACCUGGCGGCAGCGCAAAAGAUGCGUAAA | GGGACAUAAACAUAGAGGUUUAUGU<br>GAGUAGAAUUGAUUGAUAAG | pYZ_3WJrep_N11_switch | ColA/kanamycin | pYZ_3WJrep_N11_trigger | ColE1/ampicillin |
| 17 | 33 | 23.1 ± 0.8 | GGGUCCAUAUACUUCAAUUUGUUAUAGUUAUGAACA<br>GAGGAGACAUAACAUGAACAAUCAACCUAAUUCUGA<br>GAUAAACCUGGCGGCAGCGCAAAAGAUGCGUAAA | GGGAACCUAAUGUACGAAUUAGGUU<br>GAAUUGAAGUAUAUGGAAUC | pYZ_3WJrep_N33_switch | ColA/kanamycin | pYZ_3WJrep_N33_trigger | ColE1/ampicillin |
| 18 | 9 | 22.9 ± 5.3 | GGGCGAAGAUGAUACAAUUUGUUAUAGUUAUGAACA<br>GAGGAGACAUAACAUGAACAAGCCACCUCACACUCA<br>ACAUAACCUGGCGGCAGCGCAAAAGAUGCGUAAA | GGGCACCUCACCUACGUGUGAGGU<br>GGCAUUGUAUCAUCUUCGUAU | pYZ_3WJrep_N09_switch | ColA/kanamycin | pYZ_3WJrep_N09_trigger | ColE1/ampicillin |
| 19 | 12 | 22.9 ± 0.2 | GGGUGAUUGAAUAAGAAUUUGUUAUAGUUAUGAACA<br>GAGGAGACAUAACAUGAACAAGCCGAUAAUGACACU<br>AACUAACCUGGCGGCAGCGCAAAAGAUGCGUAAA | GGGCGAUAAUGUUCCGUCAUUAUC<br>GGCAUUCUUAUUCAAUCAUAG | pYZ_3WJrep_N12_switch | ColA/kanamycin | pYZ_3WJrep_N12_trigger | ColE1/ampicillin |
| 20 | 36 | 21.3 ± 0.8 | GGGAUGUGAUUACUAGAUUUGUUAUAGUUAUGAACA<br>GAGGAGACAUAACAUGAACAAUCCGAACAAGCAAUA<br>CAAGAACCUGGCGGCAGCGCAAAAGAUGCGUAAA | GGGCGAACAAGAACAUGCUUGUUCG<br>GAAUCUAGUAAUCACAUCUA | pYZ_3WJrep_N36_switch | ColA/kanamycin | pYZ_3WJrep_N36_trigger | ColE1/ampicillin |
| 21 | 42 | 20.8 ± 0.5 | GGGAUUCACUUACAAGAUUUGUUAUAGUUAUGAACA<br>GAGGAGACAUAACAUGAACAAUCACAUAGACCAUAG<br>AACGAACCUGGCGGCAGCGCAAAAGAUGCGUAAA | GGGACAUAGACAAUUUGGUCUAUGU<br>GAAUCUUGUAAGUGAAUAUG | pYZ_3WJrep_N42_switch | ColA/kanamycin | pYZ_3WJrep_N42_trigger | ColE1/ampicillin |
| 22 | 8 | 20.5 ± 0.7 | GGGACAAAUCAGAUAAACUUGUUAUAGUUAUGAACA<br>GAGGAGACAUAACAUGAACAAGCCAAUAGAUAGAACA<br>CGAAACCUGGCGGCAGCGCAAAAGAUGCGUAAA | GGGCAAUAGAUCAAACUAUCUAUUG<br>GCGUUUAUCUGAUUUGUAUG | pYZ_3WJrep_N08_switch | ColA/kanamycin | pYZ_3WJrep_N08_trigger | ColE1/ampicillin |
| 23 | 1 | 19.9 ± 2.4 | GGGCAAUUAGUACUAUCCUUGUUAUAGUUAUGAACA<br>GAGGAGACAUAACAUGAACAAGUCACAUAACCGCAA<br>ACUAAACCUGGCGGCAGCGCAAAAGAUGCGUAAA | GGGCACAUAACAUAACGGUUAUGUG<br>ACGGAUAGUACUAAUUGAAA | pYZ_3WJrep_N01_switch | ColA/kanamycin | pYZ_3WJrep_N01_trigger | ColE1/ampicillin |
| 24 | 47 | 17.5 ± 0.6 | GGGCAAUUCCUAAUCCUAUUGUUAUAGUUAUGAACA<br>GAGGAGACAUAACAUGAACAAUCCAAAUAGAGCAAAG<br>CAGAACCUGGCGGCAGCGCAAAAGAUGCGUAAA | GGGCAAAUAGAUAGAGCUCUAUUUG<br>GAUAGGAUUAGGAAUUGAAA | pYZ_3WJrep_N47_switch | ColA/kanamycin | pYZ_3WJrep_N47_trigger | ColE1/ampicillin |

| | | | | | | | | |
|---|---|---|---|---|---|---|---|---|
| 25 | 38 | 17.0 ± 1.1 | GGGAUGAUCGAAACGAAUUUGUUAUAGUUAUGAACAGAGGAGACAUAACAUGAACAAUACAAGAAUAGACAAUAGAAACCUGGCGGCAGCGCAAAAGAUGCGUAAA | GGGCAAGAAUAACAUUCUAUUCUUGUAAUUCGUUUCGAUCAUAUG | pYZ_3WJrep_N38_switch | ColA/kanamycin | pYZ_3WJrep_N38_trigger | ColE1/ampicillin |
| 26 | 25 | 16.1 ± 1.0 | GGGUAAGAAUUAGAAUCAUUGUUAUAGUUAUGAACAGAGGAGACAUAACAUGAACAAUCAGCGAACACGAAACAUAAACCUGGCGGCAGCGCAAAAGAUGCGUAAA | GGGAGCGAACAAUAACGUGUUCGCUGAUGAUUCUAAUUCUUAACA | pYZ_3WJrep_N25_switch | ColA/kanamycin | pYZ_3WJrep_N25_trigger | ColE1/ampicillin |
| 27 | 35 | 15.9 ± 1.0 | GGGAAUCCAAGUUAUCAUUUGUUAUAGUUAUGAACAGAGGAGACAUAACAUGAACAAUACACAUAAAGACCUCAUAAACCUGGCGGCAGCGCAAAAGAUGCGUAAA | GGGCACAUAAACUAAUCUUUAUGUGUAAUGAUAACUUGGAUUGAA | pYZ_3WJrep_N35_switch | ColA/kanamycin | pYZ_3WJrep_N35_trigger | ColE1/ampicillin |
| 28 | 27 | 15.8 ± 2.1 | GGGAAUGAAGAUAGUGAUUUGUUAUAGUUAUGAACAGAGGAGACAUAACAUGAACAAUCCGAAUAAGCGAACUAAGAACCUGGCGGCAGCGCAAAAGAUGCGUAAA | GGGCGAAUAAGAAUGCGCUUAUUCGGAAUCACUAUCUUCAUUAAC | pYZ_3WJrep_N27_switch | ColA/kanamycin | pYZ_3WJrep_N27_trigger | ColE1/ampicillin |
| 29 | 41 | 15.4 ± 0.5 | GGGAUCUCACCGUGUCUAUUGUUAUAGUUAUGAACAGAGGAGACAUAACAUGAACAAUCAACAUACAUCAUACAAGAACCUGGCGGCAGCGCAAAAGAUGCGUAAA | GGGAACAUACAAAUUGAUGUAUGUUGAUAGACACGGUGAGAUAGA | pYZ_3WJrep_N41_switch | ColA/kanamycin | pYZ_3WJrep_N41_trigger | ColE1/ampicillin |
| 30 | 45 | 13.2 ± 0.6 | GGGCGACUUAAACUACUAUUGUUAUAGUUAUGAACAGAGGAGACAUAACAUGAACAAUCACUCCACGGAAACUACUAACCUGGCGGCAGCGCAAAAGAUGCGUAAA | GGGACUCCACGAAUAUCCGUGGAGUGAUAGUAGUUUAAGUCGUAG | pYZ_3WJrep_N45_switch | ColA/kanamycin | pYZ_3WJrep_N45_trigger | ColE1/ampicillin |
| 31 | 4 | 12.8 ± 1.1 | GGGUAAUGAAGUGAAUACUUGUUAUAGUUAUGAACAGAGGAGACAUAACAUGAACAAGCCAAAUGAUACUAAACCGAACCUGGCGGCAGCGCAAAAGAUGCGUAAA | GGGCAAAUGAUCAUCGUAUCAUUUGGCGUAUUCACUUCAUUAACA | pYZ_3WJrep_N04_switch | ColA/kanamycin | pYZ_3WJrep_N04_trigger | ColE1/ampicillin |
| 32 | 48 | 12.4 ± 1.2 | GGGACUACUCCAUUCUAUUUGUUAUAGUUAUGAACAGAGGAGACAUAACAUGAACAAUCCAUACCUCAUACAACGAAACCUGGCGGCAGCGCAAAAGAUGCGUAAA | GGGCAUACCUCUACAAUGAGGUAUGGAAUAGAAUGGAGUAGUAAG | pYZ_3WJrep_N48_switch | ColA/kanamycin | pYZ_3WJrep_N48_trigger | ColE1/ampicillin |
| 33 | 37 | 11.1 ± 0.6 | GGGAUGUGAACUUAAAGAUUGUUAUAGUUAUGAACAGAGGAGACAUAACAUGAACAAUGCGAAAUACGGAAAUAGAAACCUGGCGGCAGCGCAAAAGAUGCGUAAA | GGGCGAAAUACAAUACCGUAUUUCGCAUCUUUAAGUUCACAUCGA | pYZ_3WJrep_N37_switch | ColA/kanamycin | pYZ_3WJrep_N37_trigger | ColE1/ampicillin |
| 34 | 44 | 10.2 ± 0.3 | GGGUACCUUCAAAUUCCAUUGUUAUAGUUAUGAACAGAGGAGACAUAACAUGAACAAUCACCUCACUAACAUUAUCAACCUGGCGGCAGCGCAAAAGAUGCGUAAA | GGGACCUCACUUACAUUAGUGAGGUGAUGGAAUUUGAAGGUAAUG | pYZ_3WJrep_N44_switch | ColA/kanamycin | pYZ_3WJrep_N44_trigger | ColE1/ampicillin |
| 35 | 18 | 9.6 ± 0.2 | GGGCAAUCCUAUUCAUCAUUGUUAUAGUUAUGAACAGAGGAGACAUAACAUGAACAAUUCUCAACAUCAAUCACUCAACCUGGCGGCAGCGCAAAAGAUGCGUAAA | GGGCUCAACAUAACAUGAUGUUGAGAAUGAUGAAUAGGAUUGAGA | pYZ_3WJrep_N18_switch | ColA/kanamycin | pYZ_3WJrep_N18_trigger | ColE1/ampicillin |
| 36 | 30 | 9.3 ± 0.7 | GGGCGAUAUGAGUAAAGAUUGUUAUAGUUAUGAACAGAGGAGACAUAACAUGAACAAUGCAAUAGAACGAUACAAGAACCUGGCGGCAGCGCAAAAGAUGCGUAAA | GGGCAAUAGAAAUACCGUUCUAUUGCAUCUUUACUCAUAUCGAUU | pYZ_3WJrep_N30_switch | ColA/kanamycin | pYZ_3WJrep_N30_trigger | ColE1/ampicillin |
| 37 | 22 | 7.5 ± 0.2 | GGGUCAAUCCGAUAAUCUUUGUUAUAGUUAUGAACAGAGGAGACAUAACAUGAACAAUUACUCCAAGGCAAUGAUAAACCUGGCGGCAGCGCAAAAGAUGCGUAAA | GGGACUCCAAGAUAAGCCUUGGAGUAAAGAUUAUCGGAUUGAAUG | pYZ_3WJrep_N22_switch | ColA/kanamycin | pYZ_3WJrep_N22_trigger | ColE1/ampicillin |
| 38 | 6 | 7.5 ± 0.4 | GGGAUUUAGAAGUAAGUAUUGUUAUAGUUAUGAACAGAGGAGACAUAACAUGAACAAUUCGAAGAUAUGGGACGAUAACCUGGCGGCAGCGCAAAAGAUGCGUAAA | GGGCGAAGAUAGAUACAUAUCUUCGAAUACUUACUUCUAAAUCUA | pYZ_3WJrep_N06_switch | ColA/kanamycin | pYZ_3WJrep_N06_trigger | ColE1/ampicillin |
| 39 | 29 | 7.5 ± 0.3 | GGGACUCCAAUUGACGAUUUGUUAUAGUUAUGAACAGAGGAGACAUAACAUGAACAAUACAUAUAGACUAAACAUAAACCUGGCGGCAGCGCAAAAGAUGCGUAAA | GGGCAUAUAGAAUAGAGUCUAUAUGUAAUCGUCAAUUGGAGUAGA | pYZ_3WJrep_N29_switch | ColA/kanamycin | pYZ_3WJrep_N29_trigger | ColE1/ampicillin |
| 40 | 31 | 7.0 ± 0.4 | GGGAAUGAAUGAUGAGAUUUGUUAUAGUUAUGAACAGAGGAGACAUAACAUGAACAAUGAGCGAACGAGCAAUGGAAACCUGGCGGCAGCGCAAAAGAUGCGUAAA | GGGAGCGAACGCGAACUCGUUCGCUCAAUCUCAUCAUUCAUUAAG | pYZ_3WJrep_N31_switch | ColA/kanamycin | pYZ_3WJrep_N31_trigger | ColE1/ampicillin |
| 41 | 39 | 6.1 ± 0.1 | GGGCGAAUUGAAAUGAAAUUGUUAUAGUUAUGAACAGAGGAGACAUAACAUGAACAAUGACGGAAUAGAAACUAAGAACCUGGCGGCAGCGCAAAAGAUGCGUAAA | GGGACGGAAUAACACUCUAUUCCGUCAUUUCAUUUCAAUUCGUAA | pYZ_3WJrep_N39_switch | ColA/kanamycin | pYZ_3WJrep_N39_trigger | ColE1/ampicillin |
| 42 | 26 | 5.6 ± 0.6 | GGGUAUUACUUACCGAUAUUGUUAUAGUUAUGAACAGAGGAGACAUAACAUGAACAAUCACUCCAACGGAACAAUCAACCUGGCGGCAGCGCAAAAGAUGCGUAAA | GGGACUCCAACUAAACCGUUGGAGUGAUAUCGGUAAGUAAUAGAA | pYZ_3WJrep_N26_switch | ColA/kanamycin | pYZ_3WJrep_N26_trigger | ColE1/ampicillin |
| 43 | 43 | 5.2 ± 0.4 | GGGCAUUUACCCUACUAUUUGUUAUAGUUAUGAACAGAGGAGACAUAACAUGAACAAUACACUCCAAAUCAAAGAUAACCUGGCGGCAGCGCAAAAGAUGCGUAAA | GGGCACUCCAAUAGGAUUUGGAGUGUAAUAGUAGGGUAAAUGAGA | pYZ_3WJrep_N43_switch | ColA/kanamycin | pYZ_3WJrep_N43_trigger | ColE1/ampicillin |
| 44 | 28 | 4.6 ± 0.1 | GGGAUUUACUUAGAUACUUUGUUAUAGUUAUGAACAGAGGAGACAUAACAUGAACAAUUACAUAUACAUACGAACGAACCUGGCGGCAGCGCAAAAGAUGCGUAAA | GGGACAUAUACUUUCAUGUAUAUGUAAAGUAUCUAAGUAAAUGAA | pYZ_3WJrep_N28_switch | ColA/kanamycin | pYZ_3WJrep_N28_trigger | ColE1/ampicillin |
| 45 | 46 | 4.0 ± 0.0 | GGGCUCAUACUUCACUAUUUGUUAUAGUUAUGAACAGAGGAGACAUAACAUGAACAAUACACCUAACAUAAUCACUAACCUGGCGGCAGCGCAAAAGAUGCGUAAA | GGGCACCUAACUAACAUGUUAGGUGUAAUAGUGAAGUAUGAGAUG | pYZ_3WJrep_N46_switch | ColA/kanamycin | pYZ_3WJrep_N46_trigger | ColE1/ampicillin |
| 46 | 5 | 2.9 ± 0.1 | GGGAUCUCCAUCUAUCCAUUGUUAUAGUUAUGAACAGAGGAGACAUAACAUGAACAAUCCAUUAAGUAACAAGCAGAACCUGGCGGCAGCGCAAAAGAUGCGUAAA | GGGCAUUAAGUUAAGUUACUUAAUGGAUGGAUAGAUGGAGAUGAU | pYZ_3WJrep_N05_switch | ColA/kanamycin | pYZ_3WJrep_N05_trigger | ColE1/ampicillin |
| 47 | 14 | 2.2 ± 0.0 | GGGUAGAUUAAGAGUGAUUUGUUAUAGUUAUGAACAGAGGAGACAUAACAUGAACAAUGACCUAAACGCACGAACUAACCUGGCGGCAGCGCAAAAGAUGCGUAAA | GGGACCUAAACUAACGCGUUUAGGUCAAUCACUCUUAAUCUACUA | pYZ_3WJrep_N14_switch | ColA/kanamycin | pYZ_3WJrep_N14_trigger | ColE1/ampicillin |
| 48 | 17 | 2.0 ± 0.1 | GGGCAAUCUCCAAUACGUUUGUUAUAGUUAUGAACAGAGGAGACAUAACAUGAACAAGCACUUAAACCGAAUAACUAACCUGGCGGCAGCGCAAAAGAUGCGUAAA | GGGACUUAAACGAAACGGUUUAAGUGCACGUAUUGGAGAUUGGGA | pYZ_3WJrep_N17_switch | ColA/kanamycin | pYZ_3WJrep_N17_trigger | ColE1/ampicillin |

**Table S4. Sequence and performance information for second-generation toehold repressors**

**NOTES:**
1. Trigger RNA sequences are listed up to the base immediately before the T7 terminator used to terminate transcription.
2. Switch RNA sequences are listed up to the 30th base (inclusive) following the end of their trigger binding domain. This 30-nt sequence contains the 21-nt linker sequence and the first 9-nts of GFPmut3b.

| Second-generation toehold repressor ranking | Second-generation toehold repressor index | GFP fold reduction | Switch sequence | Trigger sequence | Switch plasmid | Switch plasmid origin/Resistance | Trigger plasmid | Trigger plasmid origin/Resistance |
|---|---|---|---|---|---|---|---|---|
| 1 | 2 | 390.7 ± 89.4 | GGGAAAGUGAAAGUAAGAAUAAAGUGGAAAGAGACGAGAAUAAGAAGAGA<br>CAUCAUAUCUUCUUAUUCUCGUCUCUUUCUACUUUAUAAACAGAGGAG<br>AAUAAAGAUGAAAGAGACGAACCUGGCGGCAGCGCAAAAGAUGCGUAAA | GGGCGAUAGCUAUACCUGGAUAGCUAU<br>CGCAUUCUUCUUAUUCUCGUCUCUUUC<br>CACUUUAUUCUUACUUUCACUUUACU | pJK_ToeRepG2_N02_switch | ColA/kanamycin | pJK_ToeRepG2_N02_trigger | CDF/spectinomycin |
| 2 | 64 | 262.8 ± 36.2 | GGGAUGAAAGAUAAGAGAAUAGAUGAGAGAAGUGGGUUUGAAUAUGAUC<br>UAUCUAACAUCAUAAUCAAACCUACUUCUCCCAUCUAUAAACAGAGGAG<br>AAUAGAUAUGAGAAGUAGGAACCUGGCGGCAGCGCAAAAGAUGCGUAAA | GGGACGGGCCUGUAAUUCAACAGGCCC<br>GUCUUAUCAUAUUCAAACCCACUUCUC<br>UCAUCUAUUCUCUUAUCUUUCAUCGC | pJK_ToeRepG2_N64_switch | ColA/kanamycin | pJK_ToeRepG2_N64_trigger | CDF/spectinomycin |
| 3 | 19 | 246.5 ± 78.6 | GGGAUAAAGAAUGAAGUGAAUAGAGUGAUAAAGGCGGUGAUGAUAAAGAC<br>UCAAAGGCUUUAUUAUCACCGUCUUUAUCCCUCUAUUAAACAGAGGAGA<br>AAUAGAAUGAUAAAGACGAACCUGGCGGCAGCGCAAAAGAUGCGUAAA | GGGACGGUUAAGUGAUGCGACUUAACC<br>GUACACUUUAUCAUCACCGCCUUUAUC<br>ACUCUAUUCACUUCAUUCUUUAUCAC | pJK_ToeRepG2_N19_switch | ColA/kanamycin | pJK_ToeRepG2_N19_trigger | CDF/spectinomycin |
| 4 | 72 | 217.5 ± 23.3 | GGGAGAGAAAAUAGAAAGAAUGGAGGUGAAUGGAGGUGUGUGAUGUGUA<br>AUAUAAAGACACAUAACACACCUCCAUUCAGCUCCAUUAAACAGAGGAGA<br>AAUGGAAUGGAAUGGAGGAACCUGGCGGCAGCGCAAAAGAUGCGUAAA | GGGCCCGUGGUGUUGUCUUACACCACG<br>GGCAAACACAUCACACACCUCCAUUCAC<br>CUCCAUUCUUUCUAUUUUCUCUACU | pJK_ToeRepG2_N72_switch | ColA/kanamycin | pJK_ToeRepG2_N72_trigger | CDF/spectinomycin |
| 5 | 11 | 192.2 ± 30.6 | GGGAAAAUAAUGAGAAGAAUAAAGUUGAAGAUAAGGAGUUGUGUAAAGAU<br>CAUACUCCUUUACACAACUCCUUAUCUUCUACUUUAUAAACAGAGGAGA<br>AUAAAGAUGAAGAUAAGGAACCUGGCGGCAGCGCAAAAGAUGCGUAAA | GGGUGGCGCGCUAGAGGUGUAGCGCGC<br>CAAUACUUUACACAACUCCUUAUCUUCA<br>ACUUUAUUCUUCUCAUUAUUUUACU | pJK_ToeRepG2_N11_switch | ColA/kanamycin | pJK_ToeRepG2_N11_trigger | CDF/spectinomycin |
| 6 | 34 | 180.1 ± 40.3 | GGGAUAAGAAGAGUAAGAAUAGAUGUGAAGAAGUGGUGUGAUGAUUUGG<br>AACAUAAACAAAUCAUCACACCACUUCUUCUCAUCUAUAAACAGAGGAGA<br>AUAGAUAUGAAGAAGUGGAACCUGGCGGCAGCGCAAAAGAUGCGUAAA | GGGACAGACGGUAAGCGGUUACCGUCU<br>GUAAACAAAUCAUCACACCACUUCUUCA<br>CAUCUAUUCUUACUCUUCUUAUAUU | pJK_ToeRepG2_N34_switch | ColA/kanamycin | pJK_ToeRepG2_N34_trigger | CDF/spectinomycin |
| 7 | 22 | 117.8 ± 9.4 | GGGAAAGAAGUAAGAAGAAUAAAGUGUGAAUGGGCGUGUGUAAGUAAGAA<br>UACACUACUUACUAACACACGUCCAUUCACACUUUAUAAACAGAGGAGA<br>AUAAAGAUGGAAUGGACGAACCUGGCGGCAGCGCAAAAGAUGCGUAAA | GGGAGUGUGUAGCAUAAAUGCUACACAC<br>UCGACUUACUUACACACGCCCAUUCACA<br>CUUUAUUCUUCUUACUUCUUUAUU | pJK_ToeRepG2_N22_switch | ColA/kanamycin | pJK_ToeRepG2_N22_trigger | CDF/spectinomycin |
| 8 | 96 | 117.7 ± 5.8 | GGGAAUAAAGAUGGAGAAAUAAAGAAUAGAGAGACGGAAAGAAUAGUAAA<br>UCCAUACUACUAUACUUUCCGUCUCUCUAGUCUUUAUAAACAGAGGAGA<br>AUAAAGAUGAGAGAGACGAACCUGGCGGCAGCGCAAAAGAUGCGUAAA | GGGCGUGUCAGUCCCAGCUGACUGACA<br>CGCUAUACUAUUCUUUCCGUCUCUCUA<br>UUCUUUAUUUCUCCAUCUUUAUUACU | pJK_ToeRepG2_N96_switch | ColA/kanamycin | pJK_ToeRepG2_N96_trigger | CDF/spectinomycin |
| 9 | 59 | 87.8 ± 3.8 | GGGAAAGAUAAGAAGGUAAUAAGAAUGAAUGAUGGCUGUGUGUAUAUAA<br>CAGACAACUAUAUAUACACAGCUAUCAUUCUUUCUUAUAAACAGAGGAG<br>AAUAAGAAUGAAUGAUAGCAACCUGGCGGCAGCGCAAAAGAUGCGUAAA | GGGUCAGAUCAAUAACAAGAUUGAUCUG<br>ACAAUAUAUACACACAGCCAUCAUUCAU<br>UCUUAUUACCUUCUUAUCUUUCAC | pJK_ToeRepG2_N59_switch | ColA/kanamycin | pJK_ToeRepG2_N59_trigger | CDF/spectinomycin |
| 10 | 33 | 70.1 ± 10.2 | GGGAAGAUGAUGAAGAAGAUUGGAUUGAAGAUAGGGUUGGUGAGAGUAU<br>UUCAAAUCUACUCUAACCAACCAUAUCUUCGAUCCAAUAAACAGAGGAG<br>AAUUGGAAUGAAGAUAUGGAACCUGGCGGCAGCGCAAAAGAUGCGUAAA | GGGCGACUCCCAUUAUGUCAUGGGAGU<br>CGCACUACUCUCACCAACCCUAUCUUC<br>AAUCCAAUCUUCUUCAUCAUCUUAAU | pJK_ToeRepG2_N33_switch | ColA/kanamycin | pJK_ToeRepG2_N33_trigger | CDF/spectinomycin |
| 11 | 24 | 68.5 ± 12.5 | GGGAGAAGAAUGAGAAGAAUAAAGGAUGAGAGAGCGGAGAAAUGUGUAGA<br>CAAAUUAUACACACUUCUCCGUUCUCUCAGCCUUUAUAAACAGAGGAGA<br>AUAAAGAUGGAGAGAACGAACCUGGCGGCAGCGCAAAAGAUGCGUAAA | GGGCCUUAGCGAAUUAAAUUUCGCUAAG<br>GCAUUACACAUUUCUCCGCUCUCUCAU<br>CCUUUAUUCUUCUCAUUCUUCUAUU | pJK_ToeRepG2_N24_switch | ColA/kanamycin | pJK_ToeRepG2_N24_trigger | CDF/spectinomycin |
| 12 | 18 | 68.1 ± 7.0 | GGGAAGAAAGUAAAUGAGAUAAAGAAUGAGAAGACGGAUAGAGAAGUUAC<br>AAUAAAUAACUUCACUAUCCGUCUUCUCAUUCUUUAUAAACAGAGGAGA<br>AUAAAGAUGGAGAAGACGAACCUGGCGGCAGCGCAAAAGAUGCGUAAA | GGGCCGGAUACCAAAUAUAUGGUAUCCG<br>GCUAAACUUCUCUAUCCGUCUUCUCAU<br>UCUUUAUCUCAUUUACUUUCUUCAC | pJK_ToeRepG2_N18_switch | ColA/kanamycin | pJK_ToeRepG2_N18_trigger | CDF/spectinomycin |
| 13 | 51 | 61.3 ± 3.3 | GGGAAAGAUAAUGGGAAGAUAAUAGGUGGUGAGUGGAGUAAAUGUGAAC<br>CUAAAUUCUUCACACUUACUCCUCUCACCAGCUAUUAUAAACAGAGGAG<br>AAUAAUAAUGGGUGAGAGGAACCUGGCGGCAGCGCAAAAGAUGCGUAAA | GGGCGUACCCUUGCGUAUCCAAGGGUA<br>CGCGAUUCACAUUUACUCCACUCACCA<br>CCUAUUAUCUUCCCAUUAUCUUUAUU | pJK_ToeRepG2_N51_switch | ColA/kanamycin | pJK_ToeRepG2_N51_trigger | CDF/spectinomycin |
| 14 | 79 | 61.2 ± 10.3 | GGGAGAAAUAGUGGAAGAAUAAGAUUGGAGAUGUGGAGGGUUGGUAUAG<br>ACGAACAAUAUACCAACCCUCCUCAUCUCCCAUCUUAUAAACAGAGGAG<br>AAUAAGAAUGGAGAUGAGGAACCUGGCGGCAGCGCAAAAGAUGCGUAAA | GGGCCAAGUAGAGGGCGUGCUCUACUU<br>GGCAAUAUACCAACCCUCCACAUCUCCA<br>AUCUUAUUCUUCCACUAUUUCUAAC | pJK_ToeRepG2_N79_switch | ColA/kanamycin | pJK_ToeRepG2_N79_trigger | CDF/spectinomycin |
| 15 | 17 | 59.9 ± 2.2 | GGGAAAGAUGAAAUGAAGAUAGAAGGUGGAAGUUGGAGUGUGUGUAAGA<br>UUACGAAACUUACAAACACUCCUACUUCCACCUUCUAUAAACAGAGGAG<br>AAUAGAAAUGGGAAGUAGGAACCUGGCGGCAGCGCAAAAGAUGCGUAAA | GGGUCAUUCACCAACACGAUGGUGAAUG<br>ACGACUUACACACACUCCAACUUCCACC<br>UUCUAUCUUCAUUUCAUCUUUACU | pJK_ToeRepG2_N17_switch | ColA/kanamycin | pJK_ToeRepG2_N17_trigger | CDF/spectinomycin |
| 16 | 46 | 58.9 ± 3.1 | GGGAGAGAAGAUAAGAAGAUAGAACGGAAUAAGUGCGAAAGUAGAAAUAA<br>CAUCAAUAUUUCUACUUUCGCUCUUAUUCUGUUCUAUAAACAGAGGAGA<br>AUAGAAAUGAAUAAGAGCAACCUGGCGGCAGCGCAAAAGAUGCGUAAA | GGGUCAGAUGUGAUGCCUCUCACAUCU<br>GAAACAUUUCUACUUUCGCACUUAUUC<br>CGUUCUAUCUUCUUAUCUUCUCUUAU | pJK_ToeRepG2_N46_switch | ColA/kanamycin | pJK_ToeRepG2_N46_trigger | CDF/spectinomycin |
| 17 | 56 | 47.4 ± 3.3 | GGGAUGGAGAAAGAUGAUAUUGAAGGAGAAUGGAGGGUUUGGAGUGUUA<br>CAAACACUAACACUUCAAACCCUCCAUUCUUCUUCAAUAAACAGAGGAG<br>AAUUGAAAUGGAAUGGAGGAACCUGGCGGCAGCGCAAAAGAUGCGUAAA | GGGCGUAACUUGCACCUGGGCAAGUUA<br>CGCAUAACACUCCAAACCCUCCAUUCUC<br>CUUCAAUAUCAUCUUUCUCCAUCAC | pJK_ToeRepG2_N56_switch | ColA/kanamycin | pJK_ToeRepG2_N56_trigger | CDF/spectinomycin |
| 18 | 21 | 47.0 ± 2.1 | GGGAUGAAAGAUGAAGUGAUAGAAGAUGAAGAGGCGAUGAUGUGUAGUU<br>CAAUCUUUACUACAAAUCAUCGUCUCUUCAACUUCUAUAAACAGAGGAG<br>AAUAGAAAUGGAAGAGACGAACCUGGCGGCAGCGCAAAAGAUGCGUAAA | GGGUCACUAGUGGCUGACACCACUAGU<br>GAAUAACUACACAUCAUCGCCUCUUCAU<br>CUUCUAUCACUUCAUCUUUCAUAUU | pJK_ToeRepG2_N21_switch | ColA/kanamycin | pJK_ToeRepG2_N21_trigger | CDF/spectinomycin |
| 19 | 75 | 44.8 ± 3.8 | GGGAUGGAUAAAGAUGAAAUAAGAGGUGAAGAUGGCUGUUGGUAUGUUA<br>AUAAAUCGAACAUAACAACAGCUAUCUUCAUCUCUUAUAAACAGAGGAGA<br>AUAAGAAUGGAAGAUAGCAACCUGGCGGCAGCGCAAAAGAUGCGUAAA | GGGAGUCCCUGGUUUAAUAACCAGGGA<br>CUCAUAACAUACCAACAGCCAUCUUCAC<br>CUCUUAUUUCAUCUUUAUCCAUAUU | pJK_ToeRepG2_N75_switch | ColA/kanamycin | pJK_ToeRepG2_N75_trigger | CDF/spectinomycin |
| 20 | 44 | 43.0 ± 5.3 | GGGAAAGUAAGAAGUGAAAUAAAGAAUGAAGAAGGCGAAGUGUGUAAUCA<br>CAAAUCUAUUACAUACUUCGCUUUCUUCACUCUUUAUAAACAGAGGAGA<br>AUAAAGAUGGAAGAAAGCAACCUGGCGGCAGCGCAAAAGAUGCGUAAA | GGGUCAGUUCCUAGCAUCUUAGGAACU<br>GAUAUAUUACACACUUCGCCUUCUUCA<br>UUCUUUAUUUCACUUCUUACUUUACU | pJK_ToeRepG2_N44_switch | ColA/kanamycin | pJK_ToeRepG2_N44_trigger | CDF/spectinomycin |
| 21 | 88 | 42.2 ± 4.9 | GGGAAUGAAAGAAAGGAUAUAAGGUGGGAAAUGUGGGAGUGUAAGAGAC<br>ACUUCAAUUCUCUUACACUCCCUCAUUUCCGACCUUAUAAACAGAGGAG<br>AAUAAGGAUGGAAAUGAGGAACCUGGCGGCAGCGCAAAAGAUGCGUAAA | GGGCCGAGCCGCCGUUCAGGGCGGCUC<br>GGCAAUCUCUUACACUCCCACAUUUCC<br>CACCUUAUAUCCUUUCUUUCAUUACU | pJK_ToeRepG2_N88_switch | ColA/kanamycin | pJK_ToeRepG2_N88_trigger | CDF/spectinomycin |
| 22 | 77 | 40.6 ± 2.7 | GGGAGAAGAGUAGUGAAGAUUGAAAUUGAAAGAGGCGAAGUUUGUGAUC<br>AAAUAACAAUCACAAACUUCGCUUCUUUCAGUUUCAAUAAACAGAGGAG<br>AAUUGAAAUGGAAAGAAGCAACCUGGCGGCAGCGCAAAAGAUGCGUAAA | GGGACGAGGCGCAGACUUGUGCGCCUC<br>GUCAUAUCACAAACUUCGCCUCUUUCAA<br>UUUCAAUCUUCACUACUCUUCUUUA | pJK_ToeRepG2_N77_switch | ColA/kanamycin | pJK_ToeRepG2_N77_trigger | CDF/spectinomycin |
| 23 | 52 | 40.3 ± 2.8 | GGGAAAGUGGAGAAGAAUAUAGAGGGUGAGAGUGGGUUGAGAUGUUGUA<br>UUAACUUAACAACAACUCAACCUACUCUCAGCCUCUAUAAACAGAGGAG<br>AAUAGAGAUGGAGAGUAGGAACCUGGCGGCAGCGCAAAAGAUGCGUAAA | GGGCGUAGAGGUAACUAUAUACCUCUAC<br>GCAAACAACAUCUCAACCCACUCUCACC<br>CUCUAUAUUCUUCUCCACUUUAAU | pJK_ToeRepG2_N52_switch | ColA/kanamycin | pJK_ToeRepG2_N52_trigger | CDF/spectinomycin |
| 24 | 43 | 39.3 ± 1.9 | GGGAGAAAGAUAAGAAAGAUAAAGAGUGGAAUGAGGUGUGUGAUAGUACA<br>AACUCUUUACUAUAACACACCUCAUUCCAUUCUUUAUAAACAGAGGAGA<br>AUAAAGAUGGGAAUGAGGAACCUGGCGGCAGCGCAAAAGAUGCGUAAA | GGGCGACUCAACCUGCUGAGGUUGAGU<br>CGAACUACUAUCACACACCUCAUUCCAC<br>UCUUUAUCUUUCUUAUCUUUCUACU | pJK_ToeRepG2_N43_switch | ColA/kanamycin | pJK_ToeRepG2_N43_trigger | CDF/spectinomycin |
| 25 | 8 | 37.6 ± 4.1 | GGGAGAGAAAAUAGAGAAAUAGAUAGGAGGAAGUGGAGAGUGAGAGAACU<br>CUCAAAUUUCUCUUACUCUCCACUUCCUCCUAUCUAUAAACAGAGGAGA<br>AUAGAUAUGAGGAAGUGGAACCUGGCGGCAGCGCAAAAGAUGCGUAAA | GGGCCUAACCUUUGGAAUCAAAGGUUAG<br>GCAAUUCUCUCACUCUCCACUUCCUCC<br>UAUCUAUUUCUCUAUUUUCUCUAUU | pJK_ToeRepG2_N08_switch | ColA/kanamycin | pJK_ToeRepG2_N08_trigger | CDF/spectinomycin |
| 26 | 3 | 36.1 ± 12.3 | GGGAGAAAUAUGAAAUGGAUUGAAGGUGUGAAAGGGAGUAGUAGGUUGA<br>UAUUCACGCAACCUACUACUCCAUUUCACAACUUCAAUAAACAGAGGAG<br>AAUUGAAAUGGUGAAAUGGAACCUGGCGGCAGCGCAAAAGAUGCGUAAA | GGGCGACCGACGGCUCGUCCCGUCGGU<br>CGCGGCAACCUACUACUCCCUUUCACA<br>CCUUCAAUCCAUUUCAUAUUUCUAAU | pJK_ToeRepG2_N03_switch | ColA/kanamycin | pJK_ToeRepG2_N03_trigger | CDF/spectinomycin |
| 27 | 89 | 34.5 ± 1.6 | GGGAGAGAAUGGAGAAAGAUAGAAAGGAUGGAAGGGAUAGUGAGAGAUC<br>UCAUUCACAUCUCUAACUAUCCUUUCCAUCCUUUCUAUAAACAGAGGAG<br>AAUAGAAAUGAUGGAAAGGAACCUGGCGGCAGCGCAAAAGAUGCGUAAA | GGGCCUGAUAGGCGCGCUGGCCUAUCA<br>GGCGUAUCUCUCACUAUCCCUUCCAUC<br>CUUUCUAUCUUUCUCCAUUCUCUUAU | pJK_ToeRepG2_N89_switch | ColA/kanamycin | pJK_ToeRepG2_N89_trigger | CDF/spectinomycin |
| 28 | 42 | 34.1 ± 10.6 | GGGGUAAAGAUGAGUGUGAUAAGAGGUAGAGAGGCGUGUGUGAGUGUAG<br>ACAUAAGAUACACUAACACACGUCUCUCUAACUCUUAUAAACAGAGGAG<br>AAUAAGAAUGAGAGAGACGAACCUGGCGGCAGCGCAAAAGAUGCGUAAA | GGGCUUAGUGUGCUAUGCAGCACACUA<br>AGCAUUACACUCACACACGCCUCUCUAC<br>CUCUUAUCACACUCAUCUUUACAUU | pJK_ToeRepG2_N42_switch | ColA/kanamycin | pJK_ToeRepG2_N42_trigger | CDF/spectinomycin |

| | | | | | | | | |
|---|---|---|---|---|---|---|---|---|
| 29 | 15 | 34.1 ± 2.3 | GGGAAGAAAGUAAAUGAGAUAGAAGGAGAUAAGAGGAUGAGAGUAAGAGCUUAUUAUUCUUACACUCAUCCUCUUAUCUUCUUCUAUAAACAGAGGAGAAUAGAAAUGGAUAAGAGGAACCUGGCGGCAGCGCAAAAGAUGCGUAAA | GGGCGCGAGCUAGGAUGGACUAGCUCGCGCAUUCUUACUCUCAUCCUCUUAUCUCCUUCUAUCUCAUUUACUUUCUUACU | pJK_ToeRepG2_N15_switch | ColA/kanamycin | pJK_ToeRepG2_N15_trigger | CDF/spectinomycin |
| 30 | 86 | 33.9 ± 1.9 | GGGGUAAAGAUAAGAGUGAUAAGAGAGGAUGAGUCGUUGUGUGUAAGUUACUUAAACACUUACACACAACGUCUCAUCCUCUCUUAUAAACAGAGGAGAAUAAGAAUGGAUGAGACGAACCUGGCGGCAGCGCAAAAGAUGCGUAAA | GGGCGGAGCAAGGCAUAUUCCUUGCUCCGAAUACUUACACACAACGACUCAUCCUCUCUUAUCACUCUUAUCUUUACAUU | pJK_ToeRepG2_N86_switch | ColA/kanamycin | pJK_ToeRepG2_N86_trigger | CDF/spectinomycin |
| 31 | 1 | 33.2 ± 1.4 | GGGAAGAUAAAGAAAAUGAUAAGAGUGAUGAAAUGGUGUGAUGUAAGUACGGACGAGACUUACAUCACACCAUUUCAUCCCUCUUAUAAACAGAGGAGAAUAAGAAUGAUGAAAUGGAACCUGGCGGCAGCGCAAAAGAUGCGUAAA | GGGAGUAGCCAAACAAGCGUUUGGCUACUAUAACUUACAUCACACCAUUUCAUCACUCUUAUCAUUUUCUUUAUCUUCUU | pJK_ToeRepG2_N01_switch | ColA/kanamycin | pJK_ToeRepG2_N01_trigger | CDF/spectinomycin |
| 32 | 4 | 33.2 ± 9.0 | GGGAAUGAAAUAAUGGAGAUAAGUAAUAAUGAGUCGGUGUGUGCUGUUAAACAGAUCAACAGCACACACCGUCUCAUUAUUACUUAUAAACAGAGGAGAAUAAGUAUGAAUGAGACGAACCUGGCGGCAGCGCAAAAGAUGCGUAAA | GGGCCAAUGGACAACUAUUUGUCCAUUGGAAGAACAGCACACACCGACUCAUUAUUACUUAUCUCCAUUAUUUCAUUUAU | pJK_ToeRepG2_N04_switch | ColA/kanamycin | pJK_ToeRepG2_N04_trigger | CDF/spectinomycin |
| 33 | 41 | 29.3 ± 12.4 | GGGGAUUUGAAAGUGAAUAUUGGCCCUGGACGGUCGAUUGUUGUAGAUACUUCUUCAAUCUACUACAAUCGUCCGUCCAAGGCCAAUAAACAGAGGAGAAUUGGCAUGGGACGGACGAACCUGGCGGCAGCGCAAAAGAUGCGUAAA | GGGUGACCUGUCGAGGGCUCGACAGGUCAUAUAUCUACAACAAUCGACCGUCCAGGGCCAAUAUUCACUUUCAAAUCACA | pJK_ToeRepG2_N41_switch | ColA/kanamycin | pJK_ToeRepG2_N41_trigger | CDF/spectinomycin |
| 34 | 49 | 28.9 ± 1.4 | GGGAAGAUGAUAAGUGAAAUAGAAGGUAGAGAGGCGUGUGUUGAAAGUUUAAUAUCUACUUUCUACACACGUCUCUCUAACUUCUAUAAACAGAGGAGAAUAGAAAUGAGAGAGACGAACCUGGCGGCAGCGCAAAAGAUGCGUAAA | GGGACUGGAUGCAGGACGUUGCAUCCAGUCAUACUUUCAACACACGCCUCUCUACCUUCUAUUUCACUUAUCAUCUUUAU | pJK_ToeRepG2_N49_switch | ColA/kanamycin | pJK_ToeRepG2_N49_trigger | CDF/spectinomycin |
| 35 | 55 | 28.4 ± 15.3 | GGGGUAAAGAAGAUGGAGAUAAAGAGUGAAUGGAGGGUGAAGUGUGUAAUAAACUACUACACAUUUCACCCUCCAUUCAUUCUUUAUAAACAGAGGAGAAUAAAGAUGGAAUGGAGGAACCUGGCGGCAGCGCAAAAGAUGCGUAAA | GGGCUAAGCUUAUGGGUUAAUAAGCUUAGCAAUACACACUUCACCCUCCAUUCACUCUUUAUCUCCAUCUUCUUUACUUA | pJK_ToeRepG2_N55_switch | ColA/kanamycin | pJK_ToeRepG2_N55_trigger | CDF/spectinomycin |
| 36 | 61 | 28.0 ± 1.3 | GGGAGAAGAAUAAGAGUGAUAAAGUGUAGAUGUGCGGUUGUUGAUUUCUAUGAAAUUGAAAUCUACAACCGUACAUCUAAACUUUAUAAACAGAGGAGAAUAAAGAUGAGAUGUACGAACCUGGCGGCAGCGCAAAAGAUGCGUAAA | GGGAGGUAUGUACUACCAAGUACAUACCUAACGAAAUCAACAACCGCACAUCUACACUUUAUCACUCUUAUUCUUCUUUA | pJK_ToeRepG2_N61_switch | ColA/kanamycin | pJK_ToeRepG2_N61_trigger | CDF/spectinomycin |
| 37 | 65 | 26.5 ± 4.3 | GGGAGAAAGUGGAAAGAUAUAGAGUGGAUGGAGAGGUGUGAGUGUGUUAAUACAUACAACACAUUCACACCUCUCCAUCUACUCUAUAAACAGAGGAGAAUAGAGAUGAUGGAGAGGAACCUGGCGGCAGCGCAAAAGAUGCGUAAA | GGGCGACAGUGCGAUAGGACGCACUGUCGCAUAACACACUCACACCUCUCCAUCCACUCUAUAUCUUUCCACUUUCUAAU | pJK_ToeRepG2_N65_switch | ColA/kanamycin | pJK_ToeRepG2_N65_trigger | CDF/spectinomycin |
| 38 | 58 | 26.2 ± 2.6 | GGGAGAAAUAAGAAGAGUAAUGAAAGUGAGAAGUGGUGUGUGAAUAUAAUACAAACUUAUAUUUACACACCACUUCUCACUUUCAUUAAACAGAGGAGAAAUGAAAUGGAGAAGUGGAACCUGGCGGCAGCGCAAAAGAUGCGUAAA | GGGCGAACUAGUGUAGUUCCACUAGUUCGAUAUAUAUUCACACACCACUUCUCACUUUCAUUACUCUUCUUAUUUCUUAU | pJK_ToeRepG2_N58_switch | ColA/kanamycin | pJK_ToeRepG2_N58_trigger | CDF/spectinomycin |
| 39 | 36 | 25.2 ± 1.5 | GGGAAAAUAAUGAGUGGAAUAAAGGGUAAAGAGUCGAGGUGUGUAAGUGAUACAAAGACUUACACACCUCGUCUCUUUAACCUUUAUAAACAGAGGAGAAUAAAGAUGAAAGAGACGAACCUGGCGGCAGCGCAAAAGAUGCGUAAA | GGGCGGUUGCUUAUGGGCGUAAGCAACCGCAAACUUACACACCUCGACUCUUUACCCUUUAUUCCACUCAUUAUUUUCAC | pJK_ToeRepG2_N36_switch | ColA/kanamycin | pJK_ToeRepG2_N36_trigger | CDF/spectinomycin |
| 40 | 50 | 24.6 ± 1.7 | GGGAGAAUAAGAGGAAAUAUAAAGAGUGAGAGUGCGGGAUUGGUUGUAAUCACAAUAUACAACAAAUCCCGUACUCUCAAUCUUUAUAAACAGAGGAGAAUAAAGAUGGAGAGUACGAACCUGGCGGCAGCGCAAAAGAUGCGUAAA | GGGAGACCGGGAAUCCACAUUCCCGGUCUCCGUACAACCAAUCCCGCACUCUCACUCUUUAUAUUUCCUCUUAUUCUUUA | pJK_ToeRepG2_N50_switch | ColA/kanamycin | pJK_ToeRepG2_N50_trigger | CDF/spectinomycin |
| 41 | 76 | 24.5 ± 1.2 | GGGAGUAGAAGAAAUGGAAUAAAGUUUGAGAUGAGGUUGUUGCUUAGUGAAACGAUGACUAAGAAACAACCUCAUCUCACACUUUAUAAACAGAGGAGAAUAAAGAUGGAGAUGAGGAACCUGGCGGCAGCGCAAAAGAUGCGUAAA | GGGCGGUGUGUUCGGUGCCGAACACACCGAUAACUAAGCAACAACCUCAUCUCAAACUUUAUUCCAUUUCUUCUACUAAU | pJK_ToeRepG2_N76_switch | ColA/kanamycin | pJK_ToeRepG2_N76_trigger | CDF/spectinomycin |
| 42 | 73 | 23.4 ± 4.2 | GGGAAGAAUGGAGAGGUAAUAAAGAAUGAGAAGACGUGUGAGAGGAUUGAAAUAAAGAAUCCUAUCACACGUCUUCUCAUUCUUUAUAAACAGAGGAGAAUAAAGAUGGAGAAGACGAACCUGGCGGCAGCGCAAAAGAUGCGUAAA | GGGCGCCAUUAACACCACUGUUAAUGGCGCUAAAUCCUCUCACACGUCUUCUCAUUCUUUAUUACCUCUCCAUUCUUACU | pJK_ToeRepG2_N73_switch | ColA/kanamycin | pJK_ToeRepG2_N73_trigger | CDF/spectinomycin |
| 43 | 91 | 22.5 ± 1.2 | GGGAGCGCUUUAAUUCAUAUAUUUCAGACUAGUACCGUGUUUCUUUGUGAAUAUGAGACAAAGAAACACGGUACUAGUCCGAAAUAUAAACAGAGGAGAAUAUUUAUGACUAGUACCAACCUGGCGGCAGCGCAAAAGAUGCGUAAA | GGGAGUACCACAGCGUUAUCUGUGGUACUGAAACAAAGAAACACGGUACUAGUCUGAAAUAUAUGAAUUAAAGCGCUAAA | pJK_ToeRepG2_N91_switch | ColA/kanamycin | pJK_ToeRepG2_N91_trigger | CDF/spectinomycin |
| 44 | 78 | 21.8 ± 4.8 | GGGAGAAUGAAAGAAGUGAUAAGUGUGAUGAAGUGCGUUGUUAUGUAUCGACUAAACAUACAUAACAACGCUCUUCAUCUCACUUAUAAACAGAGGAGAAUAAGUAUGAUGAAGAGCAACCUGGCGGCAGCGCAAAAGAUGCGUAAA | GGGUCCCUAGGAAACCGUCUUCCUAGGGAUAAAUACAUAACAACGCACUUCAUCACACUUAUCACUUCUUUCAUUCUAUA | pJK_ToeRepG2_N78_switch | ColA/kanamycin | pJK_ToeRepG2_N78_trigger | CDF/spectinomycin |
| 45 | 53 | 21.6 ± 2.3 | GGGAUAAAGAUGGAGAAGAUAAGAUUGGAUGAAGGGUUGAGGGUAUAUAUCGAUCUUAUAUACACUCAACCUUUCAUCCAAUCUUAUAAACAGAGGAGAAUAAGAAUGGAUGAAAGGAACCUGGCGGCAGCGCAAAAGAUGCGUAAA | GGGCGUUCUUGAGAAAUAGCUCAAGAACGCAAAUAUACCCUCAACCCUUCAUCCAAUCUUAUCUUCUCCAUCUUUAUACU | pJK_ToeRepG2_N53_switch | ColA/kanamycin | pJK_ToeRepG2_N53_trigger | CDF/spectinomycin |
| 46 | 83 | 20.3 ± 1.1 | GGGAAGAAUGAAGAAGAUAUAGAAGAAGUAAGAAGGUGUGUAAGUAAUUACAAACACAUUACUUACACACCUUCUUACUUCUUCUAUAAACAGAGGAGAAUAGAAAUGGUAAGAAGGAACCUGGCGGCAGCGCAAAAGAUGCGUAAA | GGGUCCAGUCCUUAAGAACAAGGACUGGAAUUAUUACUUACACACCUUCUUACUUCUUCUAUAUCUUCUUCAUUCUUACU | pJK_ToeRepG2_N83_switch | ColA/kanamycin | pJK_ToeRepG2_N83_trigger | CDF/spectinomycin |
| 47 | 31 | 20.0 ± 2.3 | GGGAGAAGUGAGAAAGAUAUAGAGGGUGAAUGGAGGAAGGUGUGUAAGGAACAAGAACUUACAUACCUUCCUCCAUUCACCCUCUAUAAACAGAGGAGAAUAGAGAUGGAAUGGAGGAACCUGGCGGCAGCGCAAAAGAUGCGUAAA | GGGCAGCCAGGUUUGUUAGAACCUGGCUGCGACUUACACACCUUCCUCCAUUCACCCUCUAUAUCUUUCUCACUUCUAAU | pJK_ToeRepG2_N31_switch | ColA/kanamycin | pJK_ToeRepG2_N31_trigger | CDF/spectinomycin |
| 48 | 48 | 19.9 ± 3.8 | GGGAUAAGAAAGAUGGAGAUUAGAUGGAGAAAGGCCGGUUUAUGUAGAUUCAUCACAUCUACAGAAACCGGUCUUUCUCGAUCUAAUAAACAGAGGAGAAUUAGAAUGAGAAAGACCAACCUGGCGGCAGCGCAAAAGAUGCGUAAA | GGGUGCGGUUUGGUGCCCUCCAAACCGCAAUUUCUACAUAAACCGGCCUUUCUCCAUCUAAUCUCCAUCUUUCUUAUAAC | pJK_ToeRepG2_N48_switch | ColA/kanamycin | pJK_ToeRepG2_N48_trigger | CDF/spectinomycin |
| 49 | 90 | 19.6 ± 1.0 | GGGAGAAAGGUAAAGAAUAUAGAAAGUGUAGAGUGGUGUGUAAUUGUGCAUAACUUUCACAAUAACACACCUCUCUACACUUUCUAUAAACAGAGGAGAAUAGAAAUGGUAGAGAGGAACCUGGCGGCAGCGCAAAAGAUGCGUAAA | GGGCCUAGCUAUAGGAUGAUAUAGCUAGGAAGCACAAUUACACACCACUCUACACUUUCUAUAUUCUUUACCUUUCUACU | pJK_ToeRepG2_N90_switch | ColA/kanamycin | pJK_ToeRepG2_N90_trigger | CDF/spectinomycin |
| 50 | 95 | 19.1 ± 1.0 | GGGAAAGAAUGGAAGAAGAUAGAACUAAAGGAUGGGAUGUGGGAAUGUUAUCUUUAUACAUUCUCACAUCCUAUCCUUUCGUUCUAUAAACAGAGGAGAAUAGAAAUGAAGGAUAGGAACCUGGCGGCAGCGCAAAAGAUGCGUAAA | GGGCGAUUUCACUCCGACAAGUGAAAUCGCAUACAUUCCCACAUCCCAUCCUUUAGUUCUAUCUUCUUCCAUUCUUUAUU | pJK_ToeRepG2_N95_switch | ColA/kanamycin | pJK_ToeRepG2_N95_trigger | CDF/spectinomycin |
| 51 | 16 | 18.4 ± 1.9 | GGGAAAGUAAAUGAAAUGAUAAAGUGUGUGAAUGCGAUGUGUGUAGUUACUAUUCACAACUACACACAUCGUAUUCACAUACUUUAUAAACAGAGGAGAAUAAAGAUGGUGAAUACGAACCUGGCGGCAGCGCAAAAGAUGCGUAAA | GGGAGGCGCUGAAUACUCUUUCAGCGCCUAUUAACUACACACAUCGCAUUCACACACUUUAUCAUUUCAUUUACUUUACU | pJK_ToeRepG2_N16_switch | ColA/kanamycin | pJK_ToeRepG2_N16_trigger | CDF/spectinomycin |
| 52 | 82 | 18.4 ± 1.1 | GGGAGAAGAAUAAGGAAGAUAAAGAAUGAAGGAGGCGAAGGUAUAGUUGACAACUUAAACUAUACCUUCGCUUCCUUCAGUCUUUAUAAACAGAGGAGAAUAAAGAUGGAAGGAAGCAACCUGGCGGCAGCGCAAAAGAUGCGUAAA | GGGUCGGUGUAACACGAACGUUACACCGACAUAACUAUACCUUCGCCUCCUUCAUUCUUUAUCUUCCUUAUUCUUCUACU | pJK_ToeRepG2_N82_switch | ColA/kanamycin | pJK_ToeRepG2_N82_trigger | CDF/spectinomycin |
| 53 | 37 | 18.2 ± 1.6 | GGGAGAAUGAAUAAGAAGAUAAAGUGGAAAGAGGCGGUGAAAGAAGUACCUUAUAACUACUUCAUUCACCGUCUCUUUCCACUUUAUAAACAGAGGAGAAUAAAGAUGAAAGAGACGAACCUGGCGGCAGCGCAAAAGAUGCGUAAA | GGGCCACAAUUUCAGCAUGGAAAUUGUGGCAUUACUUCUUUCACCGCCUCUUUCCACUUUAUCUUCUUAUUCAUUCUCAC | pJK_ToeRepG2_N37_switch | ColA/kanamycin | pJK_ToeRepG2_N37_trigger | CDF/spectinomycin |
| 54 | 63 | 18.2 ± 1.4 | GGGAAAAUAAGAUGGAGAAUAAAGGGAGAUGGAAGGGUGAGAGAGAAACUAACACAAUUUCUCACUCACCCUUCCAUCUUCCUUUAUAAACAGAGGAGAAUAAAGAUGGAUGGAAGGAACCUGGCGGCAGCGCAAAAGAUGCGUAAA | GGGCGAUUCUCUGUUAGGUCAGAGAAUCGCAAUUUCUCUCUCACCCUUCCAUCUCCCUUUAUUCUCCAUCUUAUUUUACU | pJK_ToeRepG2_N63_switch | ColA/kanamycin | pJK_ToeRepG2_N63_trigger | CDF/spectinomycin |
| 55 | 38 | 17.9 ± 5.1 | GGGUUAAGACGCCUGAGAAUAGAAAGAAAUAAGUCGGAAGAAAGUAGUGAUCAAUAUACUACUUUCUUCCGUCUUAUUUCUUUCUAUAAACAGAGGAGAAUAGAAAUGAAUAAGACGAACCUGGCGGCAGCGCAAAAGAUGCGUAAA | GGGUGUGGCUAUCUGGUGCGAUAGCCACAUAUACUACUUUCUUCCGACUUAUUUCUUUCUAUUCUCAGGCGUCUUAAUUA | pJK_ToeRepG2_N38_switch | ColA/kanamycin | pJK_ToeRepG2_N38_trigger | CDF/spectinomycin |
| 56 | 57 | 17.6 ± 1.0 | GGGAGAGAUGAAAGAGUAAUAAAGUGGGAAAGUAGGGUAAGGUGUAUGACUAGAACACAUACACCUUACCCUACUUUCCUACUUUAUAAACAGAGGAGAAUAAAGAUGGAAAGUAGGAACCUGGCGGCAGCGCAAAAGAUGCGUAAA | GGGCCGCCAGCAACGGGAAUUGCUGGCGGCGACAUACACCUUACCCUACUUUCCCACUUUAUUACUCUUUCAUCUCUUUA | pJK_ToeRepG2_N57_switch | ColA/kanamycin | pJK_ToeRepG2_N57_trigger | CDF/spectinomycin |
| 57 | 87 | 16.8 ± 3.1 | GGGAAGAGAGUAAAGAAGAUAAGAGUGAUAAAGUGCUGUGUUGUAGUGCAUAUAAAUCACUACAACACAGCUCUUUAUCCCUCUUAUAAACAGAGGAGAAUAAGAAUGAUAAAGAGCAACCUGGCGGCAGCGCAAAAGAUGCGUAAA | GGGCGGAACUCUUGCGACUAAGAGUUCCGAAACACUACAACACAGCACUUUAUCACUCUUAUCUUCUUUACUCUCUUACU | pJK_ToeRepG2_N87_switch | ColA/kanamycin | pJK_ToeRepG2_N87_trigger | CDF/spectinomycin |
| 58 | 68 | 16.8 ± 1.4 | GGGAGAAGAGUAAGAAGAAUAGAAUGGAAUAGAAGGUUGAUGGUAGUAAAUUACAACUACUACAAUCAACCUUCUAUUCUAUUCUAUAAACAGAGGAGAAUAGAAAUGAAUAGAAGGAACCUGGCGGCAGCGCAAAAGAUGCGUAAA | GGGCGCACGAGGAUAGCAAUCCUCGUGCGAACUACUACCAUCAACCUUCUAUUCCAUUCUAUUCUUCUUACUCUUCUUAU | pJK_ToeRepG2_N68_switch | ColA/kanamycin | pJK_ToeRepG2_N68_trigger | CDF/spectinomycin |

| | | | | | | | | |
|---|---|---|---|---|---|---|---|---|
| 59 | 30 | 16.5 ± 2.1 | GGGAGAGUAAGAAGAUAUAUAGAAGUGCGAAUGAGGUGUGUAAGUAGUA<br>CAAAUAUCACUACUUACACACCUCAUUCGCUCUUCUAUAAACAGAGGAG<br>AAUAGAAAUGCGAAUGAGGAACCUGGCGGCAGCGCAAAAGAUGCGUAAA | GGGCCGUUUGUGCUCGAACGCACAAAC<br>GGAAUACUACUUACACACCUCAUUCGCA<br>CUUCUAUAUAUCUUCUUACUCUACU | pJK_ToeRepG2_N30_switch | ColA/kanamycin | pJK_ToeRepG2_N30_trigger | CDF/spectinomycin |
| 60 | 25 | 16.4 ± 2.4 | GGGAUGGAUAAGAAGUGGAUAAGAUGGAAGAUGACGAGAAUAGGUAUGG<br>ACAAUAAACAUACCUAUUCUCGUCAUCUUCGAUCUUAUAAACAGAGGAG<br>AAUAAGAAUGAAGAUGACGAACCUGGCGGCAGCGCAAAAGAUGCGUAAA | GGGCGGGAAGUCUCUAGGUAGACUUCC<br>CGCAACAUACCUAUUCUCGUCAUCUUC<br>CAUCUUAUCCACUUCUUAUCCAUCUU | pJK_ToeRepG2_N25_switch | ColA/kanamycin | pJK_ToeRepG2_N25_trigger | CDF/spectinomycin |
| 61 | 94 | 15.9 ± 1.1 | GGGAGAGAAAUGAGAAGUAAUGAAGGUGAGAAGGCGAGAAAGAUAAAGGA<br>CUUUAGACUUUAUAUUUCUCGUCUUCUCACCUUCAUUAAACAGAGGAG<br>AAAUGAAAUGGAGAAGACGAACCUGGCGGCAGCGCAAAAGAUGCGUAAA | GGGCUUCCAGGUAGCCGUUUACCUGGA<br>AGCAACUUUAUCUUUCUCGCCUUCUCA<br>CCUUCAUUACUUCUCAUUUCUCUAUA | pJK_ToeRepG2_N94_switch | ColA/kanamycin | pJK_ToeRepG2_N94_trigger | CDF/spectinomycin |
| 62 | 40 | 15.5 ± 0.4 | GGGAGAAUAAGAGAGAUAAUAAGAAAGAGUGAAUGGGUAUGUUGUGUAAA<br>CGGCAACUACACAACAUACCCUUUCACUCCUUCUUAUAAACAGAGGAGA<br>AUAAGAAUGAGUGAAAGGAACCUGGCGGCAGCGCAAAAGAUGCGUAAA | GGGAGCCAAGCCAUCUGAAUGGCUUGG<br>CUCAAUACACAACAUACCCAUUCACUCU<br>UUCUUAUUAUCUCUCUUAUUCUACU | pJK_ToeRepG2_N40_switch | ColA/kanamycin | pJK_ToeRepG2_N40_trigger | CDF/spectinomycin |
| 63 | 92 | 15.3 ± 1.1 | GGGAUAAGAAUGAAGUGGAAUGAAGAUGAAAGAGGCAAGUGUGUAGUUC<br>UUCAUAUCAACUACACACUUGCUUCUUUCAACUUCAUUAAACAGAGGAG<br>AAAUGAAAUGGAAAGAAGCAACCUGGCGGCAGCGCAAAAGAUGCGUAAA | GGGUCAUUUAGAUUAACAAAUCUAAAUG<br>AUUUAACUACACACUUGCCUCUUUCAU<br>CUUCAUUCCACUUCAUUCUUAUUAU | pJK_ToeRepG2_N92_switch | ColA/kanamycin | pJK_ToeRepG2_N92_trigger | CDF/spectinomycin |
| 64 | 66 | 14.8 ± 2.7 | GGGAAAGAGUGGAGAAGUAAUGAAGGUGAGAUGUGGAGUGUGAGUUGUG<br>ACAUAAAUACAACUAACACUCCUCAUCUCAUCUUCAUUAAACAGAGGAG<br>AAAUGAAAUGGAGAUGAGGAACCUGGCGGCAGCGCAAAAGAUGCGUAAA | GGGCCGCAAGCUCAAAUCAGAGCUUGC<br>GGCAAACAACUCACACUCCACAUCUCAC<br>CUUCAUUACUUCUCCACUCUUUAUA | pJK_ToeRepG2_N66_switch | ColA/kanamycin | pJK_ToeRepG2_N66_trigger | CDF/spectinomycin |
| 65 | 35 | 14.3 ± 3.9 | GGGAAAGAUGAAGAGGUAAUAAUGUGUGAAGAGUGCUUGUGUUGUAGAA<br>CAUUUAAUUCUACAACACAAGCUCUCUUCAUACAUUAUAAACAGAGGAG<br>AAUAAUGAUGGAAGAGAGCAACCUGGCGGCAGCGCAAAAGAUGCGUAAA | GGGCCUCUCUGGCCAGGAGGCCAGAGA<br>GGAUAUCUACAACACAAGCACUCUUCAC<br>ACAUUAUUACCUCUUCAUCUUUAAC | pJK_ToeRepG2_N35_switch | ColA/kanamycin | pJK_ToeRepG2_N35_trigger | CDF/spectinomycin |
| 66 | 7 | 13.9 ± 2.0 | GGGAAAGUAAAUGAAGUGAUAGAAGAUAGAAGUGCGAUGUGUGUUAGAU<br>UCAAACAUUCUAACACACAUCGUACUUCUAUCUUCUAUAAACAGAGGAG<br>AAUAGAAAUGAGAAGUACGAACCUGGCGGCAGCGCAAAAGAUGCGUAAA | GGGACCCUAUGCGACUAACCGCAUAGG<br>GUAAAUCUAACACACAUCGCACUUCUAU<br>CUUCUAUCACUUCAUUUACUUUAUU | pJK_ToeRepG2_N07_switch | ColA/kanamycin | pJK_ToeRepG2_N07_trigger | CDF/spectinomycin |
| 67 | 54 | 13.9 ± 1.4 | GGGGUAAGAGAAGUAGUGAUAAGAGGAAUGAGAAGGAUGUUUGUGUAUA<br>CUCGAAAUAUACACAAACAUCCAUCUCAUUGCUCUUAUAAACAGAGGAG<br>AAUAAGAAUGAUGAGAUGGAACCUGGCGGCAGCGCAAAAGAUGCGUAAA | GGGUCUGGUCAUCAAUCAUGAUGACCA<br>GACUAAUACACAAACAUCCUUCUCAUUC<br>CUCUUAUCACUACUUCUCUUACAUU | pJK_ToeRepG2_N54_switch | ColA/kanamycin | pJK_ToeRepG2_N54_trigger | CDF/spectinomycin |
| 68 | 32 | 13.6 ± 1.6 | GGGAUAAGAAAGAGUGGAAUAGAGGUUGAGGAAGGCUUGAGUUGUAGUA<br>CACACAGAACUACAACUCAAGCUUUCCUCAACCUCUAUAAACAGAGGAG<br>AAUAGAGAUGGAGGAAAGCAACCUGGCGGCAGCGCAAAAGAUGCGUAAA | GGGCCAAGCAACGUGUCGGCGUUGCUU<br>GGCAAACUACAACUCAAGCCUUCCUCAA<br>CCUCUAUUCCACUCUUUCUUAUAUU | pJK_ToeRepG2_N32_switch | ColA/kanamycin | pJK_ToeRepG2_N32_trigger | CDF/spectinomycin |
| 69 | 67 | 13.5 ± 3.8 | GGGAGAAGUAAGAAAGUGAUAGAUGUUGAAGAGUGCUGUGUGUAUUGUA<br>CAGAAAGAACAAUAAACACAGCUCUCUUCAUCAUCUAUAAACAGAGGAGA<br>AUAGAUAUGGAAGAGAGCAACCUGGCGGCAGCGCAAAAGAUGCGUAAA | GGGCCUGGGUCUAUGCGGCUAGACCCA<br>GGAUUACAAUACACACAGCACUCUUCAA<br>CAUCUAUCACUUUCUUACUUCUAAU | pJK_ToeRepG2_N67_switch | ColA/kanamycin | pJK_ToeRepG2_N67_trigger | CDF/spectinomycin |
| 70 | 47 | 13.5 ± 1.3 | GGGAAGAUGAAAGAUAGAAUAGAAUGUAGAAGAUGCGAAGUUGAUGUUC<br>AACAAUUCAACAUCAACUUCGCUUCUUCUAGAUUCUAUAAACAGAGGAG<br>AAUAGAAAUGAGAAGAAGCAACCUGGCGGCAGCGCAAAAGAUGCGUAAA | GGGUCUAAGAAUAGGUUUAUAUUCUUAG<br>AAAUAACAUCAACUUCGCAUCUUCUACA<br>UUCUAUUCUAUCUUUCAUCUUACU | pJK_ToeRepG2_N47_switch | ColA/kanamycin | pJK_ToeRepG2_N47_trigger | CDF/spectinomycin |
| 71 | 28 | 12.7 ± 1.6 | GGGAAAAUAAUGGGAGAGAUAAGAGGGAUAAAGAGGGUGAGAAGUAGAAA<br>CUUAACCUCUACUACUCACCCUCUUUAUCCCUCUUAUAAACAGAGGAGA<br>AUAAGAAUGAUAAAGAGGAACCUGGCGGCAGCGCAAAAGAUGCGUAAA | GGGCGAGAGAUGUAUACUAACAUCUCUC<br>GCAAUCUACUUCUCACCCUCUUUAUCC<br>CUCUUAUCUCUCCCAUUAUUUUCAC | pJK_ToeRepG2_N28_switch | ColA/kanamycin | pJK_ToeRepG2_N28_trigger | CDF/spectinomycin |
| 72 | 9 | 12.4 ± 1.6 | GGGAAAGUAAGAAAGAGUAUAAGAGAGAUAGAAAGGUGUGCGCGUAUAC<br>UGACGAGAUAUACGAGCACACCUUUCUAUCUCUCUUAUAAACAGAGGAG<br>AAUAAGAAUGAUAGAAAGGAACCUGGCGGCAGCGCAAAAGAUGCGUAAA | GGGUCCCGGAAACCGAAUCGUUUCCGG<br>GAAUUUAUACGCGCACACCUUUCUAUC<br>UCUCUUAUACUCUUUCUUACUUUAUU | pJK_ToeRepG2_N09_switch | ColA/kanamycin | pJK_ToeRepG2_N09_trigger | CDF/spectinomycin |
| 73 | 29 | 12.3 ± 4.5 | GGGAAAGUGAAUGAGAUUAUUAUUGGAGGUAUUUGGAGAGAAUGUUAUA<br>CGGCACAUAUAACAAUCUCUCCUAAUACCUACAAUAAUAAACAGAGGAGA<br>AUUAUUAUGGGUAUUAGGAACCUGGCGGCAGCGCAAAAGAUGCGUAAA | GGGUCCAGCUCCCAUGAAAGGGAGCUG<br>GACAAAUAACAUUCUCUCCAAAUACCUC<br>CAAUAAUAAUCUCAUUCACUUUAAA | pJK_ToeRepG2_N29_switch | ColA/kanamycin | pJK_ToeRepG2_N29_trigger | CDF/spectinomycin |
| 74 | 80 | 12.1 ± 0.6 | GGGAGAGAUGAUGGAAGAAUAAAGAGUGAGAUGAGGUUGUGUAGUAAGG<br>AGAAAUUACUUACUACACAACCUCAUCUCACUCUUUAUAAACAGAGGAG<br>AAUAAAGAUGGAGAUGAGGAACCUGGCGGCAGCGCAAAAGAUGCGUAAA | GGGCGUGGAGAUCUUAUAAGAUCUCCA<br>CGAAACUUACUACACAACCUCAUCUCAC<br>UCUUUAUUCUUCCAUCAUCUCUUAU | pJK_ToeRepG2_N80_switch | ColA/kanamycin | pJK_ToeRepG2_N80_trigger | CDF/spectinomycin |
| 75 | 13 | 11.7 ± 2.2 | GGGAUGAGAAAUAAGAAGAUAAUAGAAAGAAAUGGCGGUGAAAGAGUAUC<br>AGCCGUAUACUCUAUCACCGCUAUUUCUUUCUAUUAUAAACAGAGGAGA<br>AUAAUAAUGAGAAAUAGCAACCUGGCGGCAGCGCAAAAGAUGCGUAAA | GGGAGACAAGCCAACUCAAUGGCUUGUC<br>UCAUUACUCUUUCACCGCCAUUUCUUU<br>CUAUUAUCUUCUUAUUUCUCAUCAC | pJK_ToeRepG2_N13_switch | ColA/kanamycin | pJK_ToeRepG2_N13_trigger | CDF/spectinomycin |
| 76 | 23 | 11.7 ± 0.8 | GGGAAUGAAAUAGAAAGAAUAAGAAGUGAAAGUGCGAUAGAUAAAGUUCA<br>CUUAAUUAACUUUAUCUAUCGUACUUUCACUUCUUAUAAACAGAGGAGA<br>AUAAGAAUGGAAAGUACGAACCUGGCGGCAGCGCAAAAGAUGCGUAAA | GGGUCGCGUUUCCAGAUUUGGAAACGC<br>GAUUUAACUUUAUCUAUCGCACUUUCA<br>CUUCUUAUUCUUUCUAUUUCAUUAUU | pJK_ToeRepG2_N23_switch | ColA/kanamycin | pJK_ToeRepG2_N23_trigger | CDF/spectinomycin |
| 77 | 12 | 11.6 ± 3.3 | GGGAAAGUGAAAGUAAGUAUAAAGUGUGAUGAGUGGUGUGAAUGUUGUA<br>ACAAACAUACAACAAUCACACCUCUCAUCAUACUUUAUAAACAGAGGAGA<br>AUAAAGAUGGAUGAGAGGAACCUGGCGGCAGCGCAAAAGAUGCGUAAA | GGGCGAUCAUAACCAAUAGGUUAUGAUC<br>GAUAACAACAUUCACACCACUCAUCACA<br>CUUUAUACUUACUUUCACUUUCUU | pJK_ToeRepG2_N12_switch | ColA/kanamycin | pJK_ToeRepG2_N12_trigger | CDF/spectinomycin |
| 78 | 10 | 11.1 ± 1.9 | GGGAGAAGUAAGAAGAUAAUAAAGGUUGAUGAAGGGUGUAAGUGUAGUA<br>UCUACACAACUACACUUACACCUUUCAUCAACCUUUAUAAACAGAGGAG<br>AAUAAAGAUGGAUGAAAGGAACCUGGCGGCAGCGCAAAAGAUGCGUAAA | GGGCGAAUUCAGUUAUGACACUGAAUUC<br>GCGAACUACACUUACACCCUUCAUCAAC<br>CUUUAUUAUCUUCUUACUUCUCUU | pJK_ToeRepG2_N10_switch | ColA/kanamycin | pJK_ToeRepG2_N10_trigger | CDF/spectinomycin |
| 79 | 5 | 10.9 ± 1.0 | GGGAAAGAUAAAGAGAGUAUAAAGAGUGAUGGAGGGUAUGAUAGUUGUA<br>CAAUAAGAACAACUAUCAUACCUUCCAUCAAUCUUUAUAAACAGAGGAGA<br>AUAAAGAUGGAUGGAAGGAACCUGGCGGCAGCGCAAAAGAUGCGUAAA | GGGCGUGUGGUCGGACCUACGACCACA<br>CGACAACAACUAUCAUACCCUCCAUCAC<br>UCUUUAUACUCUCUUUAUCUUUAUU | pJK_ToeRepG2_N05_switch | ColA/kanamycin | pJK_ToeRepG2_N05_trigger | CDF/spectinomycin |
| 80 | 60 | 10.7 ± 1.2 | GGGAGAAAUAUGGAUGAAAUAAAGUGGGAAAUAGGGUGUGUGAUGAAUA<br>AACAUUUGAUUCAUAACACACCUUAUUUCCCACUUUAUAAACAGAGGAG<br>AAUAAAGAUGGAAAUAAGGAACCUGGCGGCAGCGCAAAAGAUGCGUAAA | GGGCUGCACUGCACUCAAUUGCAGUGC<br>AGCAAAUUCAUCACACACCCUAUUUCCC<br>ACUUUAUUUCAUCCAUAUUUCUUAU | pJK_ToeRepG2_N60_switch | ColA/kanamycin | pJK_ToeRepG2_N60_trigger | CDF/spectinomycin |
| 81 | 74 | 10.6 ± 0.9 | GGGAUGAAGAAUGAAGUGAUAAGAAGUAGAAGAACGUGUAAAUGUAAUUC<br>UACAAACAUUACAUUUACACGUUCUUCUACUUCUUAUAAACAGAGGAGA<br>AUAAGAAUGAGAAGAACGAACCUGGCGGCAGCGCAAAAGAUGCGUAAA | GGGUCCGUCAUUAUUUCUAUAAUGACG<br>GAAUUAUUACAUUUACACGUUCUUCUAC<br>UUCUUAUCACUUCAUUCUUCAUUUA | pJK_ToeRepG2_N74_switch | ColA/kanamycin | pJK_ToeRepG2_N74_trigger | CDF/spectinomycin |
| 82 | 69 | 9.9 ± 1.8 | GGGAGAAUGGAGAGAGUAAUAAGAUGGAAGAUAAGGACAGAAUGCAAAGU<br>GUUUAAAUUUGCAUUCUGUCCUUAUCUUCCAUCUUAUAAACAGAGGAGA<br>AUAAGAAUGAAGAUAAGGAACCUGGCGGCAGCGCAAAAGAUGCGUAAA | GGGUUGAAGCUUUCCACGGAAAGCUUC<br>AACUGUUUGCAUUCUGUCCUUAUCUUC<br>CAUCUUAUUACUCUCUCCAUUCUACU | pJK_ToeRepG2_N69_switch | ColA/kanamycin | pJK_ToeRepG2_N69_trigger | CDF/spectinomycin |
| 83 | 20 | 9.4 ± 1.8 | GGGAAGAUGAAGAGAGAUAUAAGAGAUGAAGAGAGCGUGUGUUGUAGAG<br>UUAAUUUAUCUACAACACACGCUCUCUUCAGCUCUUAUAAACAGAGGAG<br>AAUAAGAAUGGAAGAGAGCAACCUGGCGGCAGCGCAAAAGAUGCGUAAA | GGGUCAGUCGAGUUGUUGGACUCGACU<br>GAAUAUCUACAACACACGCUCUCUUCAU<br>CUCUUAUAUCUCUCUUCAUCUUAAU | pJK_ToeRepG2_N20_switch | ColA/kanamycin | pJK_ToeRepG2_N20_trigger | CDF/spectinomycin |
| 84 | 26 | 9.2 ± 2.7 | GGGAAGGUAAGAGAAGUGAUAAAUGUGAAGAUAAGGUGUAGAUGUGUAG<br>GAUAUAAAUACACAACUACACCUUAUCUUCUCAUUUAUAAACAGAGGAG<br>AAUAAAUAUGAAGAUAAGGAACCUGGCGGCAGCGCAAAAGAUGCGUAAA | GGGUCCAGGAUAAGAAUAUUUAUCCUGG<br>AAAUUACACAUCUACACCUUAUCUUCAC<br>AUUUAUCACUUCUCUUACCUUAUU | pJK_ToeRepG2_N26_switch | ColA/kanamycin | pJK_ToeRepG2_N26_trigger | CDF/spectinomycin |
| 85 | 45 | 8.6 ± 1.1 | GGGAUUAAUGGGAUGUGGAUUGUAGUUAAGAAGAGGUGCGUAGCUAGU<br>UCUCAACAUACUAGCCACGCACCACUUCUUAUCUACAAUAAACAGAGGA<br>GAAUUGUAAUGAAGAAGUGGAACCUGGCGGCAGCGCAAAAGAUGCGUAA<br>A | GGGUGGCACUAGGCGUUACCCUAGUGC<br>CAUAUACUAGCUACGCACCUCUUCUUAA<br>CUACAAUCCACAUCCCAUUAAUAAG | pJK_ToeRepG2_N45_switch | ColA/kanamycin | pJK_ToeRepG2_N45_trigger | CDF/spectinomycin |
| 86 | 71 | 7.2 ± 10.9 | GGGGUAACUAAGAAUACUAUAAAGGGCACGGAAAGGAUUAGGAAAUUGAA<br>UUUAAUACAAUUUACUAAUCCUUUCCGUGGCCUUUAUAAACAGAGGAGA<br>AUAAAGAUGACGGAAAGGAACCUGGCGGCAGCGCAAAAGAUGCGUAAA | GGGAGUCGCCAAUCCGCACAUUGGCGA<br>CUUGGCAAUUUCCUAAUCCUUUCCGUG<br>CCCUUUAUAGUAUUCUUAGUUACACA | pJK_ToeRepG2_N71_switch | ColA/kanamycin | pJK_ToeRepG2_N71_trigger | CDF/spectinomycin |
| 87 | 6 | 6.3 ± 0.7 | GGGAAAGUGAAAGUGAAGAUAAAUGAAGUGAAGAGGUGUGUUGCUGUAA<br>CAACACAGUACAGCGACACACCUCUUCACUUCAUUUAUAAACAGAGGAG<br>AAUAAAUAUGGUGAAGAGGAACCUGGCGGCAGCGCAAAAGAUGCGUAAA | GGGUCCCUCAAGAAACAGAUCUUGAGGG<br>AAUAUACAGCAACACACCUCUUCACUUC<br>AUUUAUCUUCACUUUCACUUUACU | pJK_ToeRepG2_N06_switch | ColA/kanamycin | pJK_ToeRepG2_N06_trigger | CDF/spectinomycin |
| 88 | 62 | 5.5 ± 0.7 | GGGAGGAAGAGUAAGUGGAUAGUGGAAGAAGUGUGGUAUGUAAGUUGUA<br>ACAAAGCGACAACUCACAUACCUCACUUCUUCCACUAUAAACAGAGGAG<br>AAUAGUGAUGGAAGUGAGGAACCUGGCGGCAGCGCAAAAGAUGCGUAAA | GGGCGGUACCCGUACGUGAACGGGUAC<br>CGCAAACAACUUACAUACCACACUUCUU<br>CCACUAUCCACUUACUCUUCCUAAU | pJK_ToeRepG2_N62_switch | ColA/kanamycin | pJK_ToeRepG2_N62_trigger | CDF/spectinomycin |

| | | | | | | | | |
|---|---|---|---|---|---|---|---|---|
| 89 | 81 | 5.3 ± 0.7 | GGGAGAGAUGAAGAGAAUAUUGGAGAGGAAUGGAGGUUGAUGAGUUGUA<br>CUUCAGACACAACUAAUCAACCUCCAUUCCUCUCCAAUAAACAGAGGAG<br>AAUUGGAAUGGAAUGGAGGAACCUGGCGGCAGCGCAAAAGAUGCGUAAA | GGGUCGCAACUCAUCAUGCUGAGUUGC<br>GACAAACAACUCAUCAACCUCCAUUCCU<br>CUCCAAUAUUCUCUUCAUCUCUACU | pJK_ToeRepG2_N81_switch | ColA/kanamycin | pJK_ToeRepG2_N81_trigger | CDF/spectinomycin |
| 90 | 84 | 4.9 ± 0.6 | GGGAUGGAAAGUGAUAAGAUAAAGUGUAAAGAGUCGUGUGUUGUAAGUC<br>AAUACAAUACUUACAACACACGUCUCUUUAUACUUUAUAAACAGAGGAG<br>AAUAAAGAUGAAAGAGACGAACCUGGCGGCAGCGCAAAAGAUGCGUAAA | GGGCCAGCGCCUAGACACAUAGGCGCU<br>GGACAACUUACAACACACGACUCUUUAC<br>ACUUUAUCUUAUCACUUUCCAUUAU | pJK_ToeRepG2_N84_switch | ColA/kanamycin | pJK_ToeRepG2_N84_trigger | CDF/spectinomycin |
| 91 | 14 | 4.0 ± 0.7 | GGGAAGAAAGUAAGUGAAAUAAAGUGUGAAGAGGCGUGUGAUUGUAGUC<br>AAGAAACAACUACAAUCACACGUCUCUUCAGACUUUAUAAACAGAGGAGA<br>AUAAAGAUGGAAGAGACGAACCUGGCGGCAGCGCAAAAGAUGCGUAAA | GGGUCAUCUUAUCGCUCAGGAUAAGAU<br>GAAUAACUACAAUCACACGCCUCUUCAC<br>ACUUUAUUUCACUUACUUUCUUUAU | pJK_ToeRepG2_N14_switch | ColA/kanamycin | pJK_ToeRepG2_N14_trigger | CDF/spectinomycin |
| 92 | 70 | 3.8 ± 4.2 | GGGAGAGAAUUGAGAUAUAUUAGAUGGAGGAGAUCGUUUGUUAUUAGUU<br>CUCUACUUACUAAUAACAAACGUUCUCCUCCAUCUAAUAAACAGAGGAG<br>AAUUAGAAUGAGGAGAACGAACCUGGCGGCAGCGCAAAAGAUGCGUAAA | GGGUGCCCGCGCAUAAGCGUGCGCGGG<br>CAAUAACUAAUAACAAACGAUCUCCUCC<br>AUCUAAUAUAUCUCAAUUCUCUAAC | pJK_ToeRepG2_N70_switch | ColA/kanamycin | pJK_ToeRepG2_N70_trigger | CDF/spectinomycin |
| 93 | 39 | 3.7 ± 0.2 | GGGAAAGAUAAGAAAGUGAUAAAGUGUGAAGAAGGCGUGUGUAGUAUAA<br>GAACACAAUAUACUACACACGCUUUCUUCAAACUUUAUAAACAGAGGAG<br>AAUAAAGAUGGAAGAAAGCAACCUGGCGGCAGCGCAAAAGAUGCGUAAA | GGGUCUAGAUCGGUCGCUCCCGAUCUA<br>GAAAUUAUACUACACACGCCUUCUUCAC<br>ACUUUAUCACUUUCUUAUCUUUACU | pJK_ToeRepG2_N39_switch | ColA/kanamycin | pJK_ToeRepG2_N39_trigger | CDF/spectinomycin |
| 94 | 27 | 2.9 ± 0.3 | GGGAACUUGAUUGGUGAAAUUGGUUGUCUUUAUUCGUAUAUUCGUUUC<br>GAAACAAAUGAAACGAAUAUACGUAUAAAGAUAACCAAUAAACAGAGGAG<br>AAUUGGUAUGCUUUAUACGAACCUGGCGGCAGCGCAAAAGAUGCGUAAA | GGGCGUUAAACAGCGCAUCCUGUUUAA<br>CGAAAGAAACGAAUAUACGAAUAAAGACA<br>ACCAAUUUCACCAAUCAAGUUAUA | pJK_ToeRepG2_N27_switch | ColA/kanamycin | pJK_ToeRepG2_N27_trigger | CDF/spectinomycin |
| 95 | 85 | 2.0 ± 0.3 | GGGAGAAUAAAUGUGGAGAUAAUGUGUGAAAGAUCGUGUGUUUAUGUUA<br>CAAACACUAACAUAAACACACGUUCUUUCAGACAUUAUAAACAGAGGAGA<br>AUAAUGAUGGAAAGAACGAACCUGGCGGCAGCGCAAAAGAUGCGUAAA | GGGCGCGGUCCGAAUCCUUUCGGACCG<br>CGGACAACAUAAACACACGAUCUUUCAC<br>ACAUUAUCUCCACAUUUAUUCUAAU | pJK_ToeRepG2_N85_switch | ColA/kanamycin | pJK_ToeRepG2_N85_trigger | CDF/spectinomycin |
| 96 | 93 | 1.7 ± 0.1 | GGGAGAAGUAAGAAAGUGAUAAGAUGUAGAAAUGCCGUGAUGUGUUAUC<br>AGCAAGAAAUAACAAAUCACGGUAUUUCUAUAUCUUAUAAACAGAGGAGA<br>AUAAGAAUGAGAAAUACCAACCUGGCGGCAGCGCAAAAGAUGCGUAAA | GGGUCUGCUAUUUACACUCAAAUAGCAG<br>AAUUAUAACACAUCACGGCAUUUCUACA<br>UCUUAUCACUUUCUUACUUCUUAU | pJK_ToeRepG2_N93_switch | ColA/kanamycin | pJK_ToeRepG2_N93_trigger | CDF/spectinomycin |

**Table S5. Shortened Triggers for SHAPE-Seq Study of 3WJ Repressors 21 and 13**

**NOTE:**
Trigger RNA sequences include the T7 terminator sequence

| Trigger name | Shortened 3WJ Repressor Trigger Sequence |
|---|---|
| Trig7_len18 | GGGUGCUCGCGUUAGAGCACCCAGUGUCGGAUAAGUGAAUUAGCAUAACCCCUUGGGGCCUCUAAACGGGUCUUGAGGGGUUUUUUG |
| Trig7_len20 | GGGUGCUCGCGUUAGAGCACCCUAGUGUCGGAUAAGUGAAUAUAGCAUAACCCCUUGGGGCCUCUAAACGGGUCUUGAGGGGUUUUUUG |
| Trig7_len25 | GGGUGCUCGCGUUAGAGCACCCGUAGUGUCGGAUAAGUGAAUAGUAGCAUAACCCCUUGGGGCCUCUAAACGGGUCUUGAGGGGUUUUUUG |
| Trig8_len18 | GGGCCUGCGGCAGAGCAGGCCCUGUAUGUGAUUUGUAUUUUAGCAUAACCCCUUGGGGCCUCUAAACGGGUCUUGAGGGGUUUUUUG |
| Trig8_len20 | GGGUGCUCGCGUUAGAGCACCCUUGUAUGUGAUUUGUAUUUGUAGCAUAACCCCUUGGGGCCUCUAAACGGGUCUUGAGGGGUUUUUUG |

**Table S6. Indices for Othogonal Sets of Toehold and 3WJ Repressors**

**A. Sets of orthogonal toehold repressors with different crosstalk levels**

***Toehold Repressor Libraries Selected After 3-hr Induction***

| Library Size | Toehold Repressor Indices | Library Dynamic Range (3 hr) |
|---|---|---|
| 14 | 3,4,16,30,36,41,42,49,65,76,78,86,91,95 | 1.5 |
| 13 | 3,4,16,30,36,41,42,49,65,78,86,91,95 | 1.6 |
| 12 | 3,4,16,30,36,42,49,59,65,78,91,95 | 1.7 |
| 11 | 3,16,30,36,41,42,49,65,76,78,86 | 2.5 |
| 10 | 3,16,30,36,41,42,65,76,78,86 | 2.8 |
| 9 | 3,4,16,30,36,49,86,91,95 | 4.9 |
| 8 | 16,30,36,42,76,78,86,91 | 7.0 |
| 7 | 16,30,42,78,86,91,95 | 7.3 |
| 6 | 3,16,36,49,78,86 | 7.5 |
| 5 | 16,30,49,78,86 | 8.7 |
| 4 | 16,30,78,86 | 12.0 |
| 3 | 30,36,78 | 15.8 |
| 2 | 59,77 | 24.4 |

**B. Sets of orthogonal 3WJ repressors with different crosstalk levels**

***3WJ Repressor Libraries Selected After 3-hr Induction***

| Library Size | 3WJ Repressor Indices | Library Dynamic Range (3 hr) | Library Dynamic Range (4 hr) | Library Dynamic Range (5 hr) |
|---|---|---|---|---|
| 16 | 2,3,7,10,11,12,13,15,19,20,21,23,24,32,34,40 | 3.1 | 6.5 | 11.6 |
| 15 | 2,3,7,10,11,12,13,15,20,21,23,24,32,34,40 | 18.2 | 24.9 | 23.9 |
| 14 | 2,3,7,10,11,13,15,20,21,23,24,32,34,40 | 18.5 | 24.9 | 23.9 |
| 13 | 2,3,7,10,11,13,15,20,21,23,24,32,34 | 20.1 | 24.9 | 23.9 |
| 12 | 2,3,10,11,13,15,20,21,23,24,32,34 | 23.9 | 28.6 | 23.9 |
| 11 | 2,3,10,11,13,15,20,21,24,32,34 | 24.2 | 28.6 | 23.9 |
| 10 | 2,3,10,13,20,21,23,24,32,34 | 25.6 | 28.6 | 23.9 |
| 9 | 2,3,10,13,20,21,23,24,32 | 28.4 | 28.6 | 23.9 |
| 8 | 2,3,10,13,20,21,23,32 | 29.2 | 57.4 | 42.1 |
| 7 | 3,10,13,20,23,24,32 | 30.8 | 48.0 | 41.3 |
| 6 | 10,20,21,23,24,32 | 40.0 | 47.4 | 42.1 |
| 5 | 10,20,23,24,32 | 43.5 | 51.1 | 42.1 |
| 4 | 19,20,21,32 | 47.7 | 62.6 | 50.6 |
| 3 | 19,20,32 | 63.8 | 118.7 | 100.6 |
| 2 | 20,32 | 87.1 | 145.0 | 122.5 |

***3WJ Repressor Libraries Selected After 4-hr Induction***

| Library Size | 3WJ Repressor Indices | Library Dynamic Range (3 hr) | Library Dynamic Range (4 hr) | Library Dynamic Range (5 hr) |
|---|---|---|---|---|
| 16 | 2,3,7,10,11,12,13,15,19,20,21,23,24,32,34,40 | 3.1 | 6.5 | 11.6 |
| 15 | 2,3,7,10,11,12,13,15,20,21,23,24,32,34,40 | 18.2 | 24.9 | 23.9 |
| 14 | 2,3,10,11,12,13,15,20,21,23,24,32,34,40 | 18.2 | 28.6 | 23.9 |
| 13 | 3,10,11,12,13,15,20,21,23,24,32,34,40 | 18.2 | 32.7 | 32.9 |
| 12 | 3,10,11,13,15,20,21,23,24,32,34,40 | 18.5 | 39.3 | 32.9 |
| 11 | 2,3,10,13,15,20,21,23,32,34,40 | 18.5 | 40.0 | 42.1 |
| 10 | 2,3,10,13,15,20,21,23,32,40 | 18.5 | 41.7 | 42.1 |
| 9 | 2,3,10,13,15,20,21,23,32 | 23.9 | 50.9 | 42.1 |
| 8 | 2,3,10,13,20,21,23,32 | 29.2 | 57.4 | 42.1 |
| 7 | 2,3,10,13,20,21,32 | 29.2 | 61.3 | 50.6 |
| 6 | 2,3,10,13,20,32 | 33.6 | 70.3 | 66.7 |
| 5 | 3,10,13,20,32 | 36.2 | 75.8 | 66.7 |
| 4 | 10,13,20,32 | 40.0 | 90.2 | 66.7 |
| 3 | 2,20,21 | 42.4 | 126.5 | 131.0 |
| 2 | 2,20 | 67.8 | 197.5 | 237.2 |

***3WJ Repressor Libraries Selected After 5-hr Induction***

| Library Size | 3WJ Repressor Indices | Library Dynamic Range (3 hr) | Library Dynamic Range (4 hr) | Library Dynamic Range (5 hr) |
|---|---|---|---|---|
| 16 | 2,3,7,10,11,12,13,15,19,20,21,23,24,32,34,40 | 3.1 | 6.5 | 11.6 |
| 15 | 2,3,7,10,11,12,13,15,19,20,21,23,24,32,34 | 14.0 | 23.9 | 23.9 |
| 14 | 2,3,7,10,11,12,13,15,19,20,21,23,32,34 | 14.0 | 23.9 | 38.0 |
| 13 | 2,3,10,11,12,13,15,20,21,23,32,34,40 | 18.2 | 32.7 | 42.1 |
| 12 | 2,3,10,11,12,13,15,19,20,21,23,34 | 14.0 | 32.7 | 55.1 |
| 11 | 2,3,10,11,12,13,15,20,21,34,40 | 18.4 | 34.4 | 61.1 |
| 10 | 2,3,10,12,13,15,19,20,21,34 | 14.0 | 33.3 | 64.3 |
| 9 | 2,3,10,12,13,15,20,21,40 | 18.4 | 34.4 | 69.0 |
| 8 | 2,3,10,13,15,20,21,40 | 18.5 | 41.7 | 72.7 |
| 7 | 2,3,10,13,15,20,21 | 24.3 | 52.3 | 86.2 |
| 6 | 2,10,13,15,20,21 | 24.3 | 52.3 | 118.4 |
| 5 | 2,10,13,15,20 | 27.0 | 63.6 | 145.9 |
| 4 | 2,10,13,20 | 33.6 | 75.1 | 164.6 |
| 3 | 10,13,20 | 40.0 | 90.2 | 196.6 |
| 2 | 2,20 | 67.8 | 197.5 | 237.2 |

**Table S7. mRNA Sensors and Trigger Sequences**

**A. Toehold Repressor mRNA Sensors**

| mRNA Sensor | Sensor Sequence | mRNA Target Subsequence | Full mRNA Target Sequence | Switch Plasmid |
|---|---|---|---|---|
| kanR sensor | GGGUUAUUCAUAUCAGGAUUAUCAAUACCAUAUUUUUGAAAAA<br>GCCGUUUCUGUAAUGAAGGACUCCACAUAUCCUUCGUUACAGA<br>UACGGCUUGUUCAAAAAAACAGAGGAGAUUUUGAAUGAGCCGU<br>AUCAACCUGGCGGCAGCGCAAAAGAUGCGUAAA | UCCUUCAUUACAGAAACGGCUUUUUCAAAAAU<br>AUGGUAUUGAUAAUCCUGAUAUGAAUAA | AUGAGCCAUAUUCAACGGGAAACGUCUUGCUCUAGGCCGCGAUUAAAUUCCAACAUGGAUGCUGAUUUAUAUGGGUAUAAAUGGGCUCGCGAUAAUGUCGGGCAA<br>UCAGGUGCGACAAUCUAUCGAUUGUAUGGGAAGCCCGAUGCGCCAGAGUUGUUUCUGAAACAUGGCAAAGGUAGCGUUGCCAAUGAUGUUACAGAUGAGAUGGU<br>CAGACUAAACUGGCUGACGGAAUUUAUGCCUCUUCCGACCAUCAAGCAUUUUAUCCGUACUCCUGAUGAUGCAUGGUUACUCACCACUGCGAUCCCCGGGAAAA<br>CAGCAUUCCAGGUAUUAGAAGAAUAUCCUGAUUCAGGUGAAAAUAUUGUUGAUGCGCUGGCAGUGUUCCUGCGCCGGUUGCAUUCGAUUCCUGUUUGUAAUUGU<br>CCUUUUAACAGCGAUCGCGUAUUUCGUCUCGCUCAGGCGCAAUCACGAAUGAAUAACGGUUUGGUUGAUGCGAGUGAUUUUGAUGACGAGCGUAAUGGCUGGCC<br>UGUUGAACAAGUCUGGAAAGAAAUGCAUAAACUUUUGCCAUUCUCACCGGAUUCAGUCGUCACUCAUGGUGAUUUCUCACUUGAUAACCUUAUUUUUGACGAGG<br>GGAAAUUAAUAGGUUGUAUUGAUGUUGGACGAGUCGGAAUCGCAGACCGAUACCAGGAUCUUGCCAUCCUAUGGAACUGCCUCGGUGAGUUUUCUCCUUCAUUA<br>CAGAAACGGCUUUUUCAAAAAUAUGGUAUUGAUAAUCCUGAUAUGAAUAAAUUGCAGUUUCAUUUGAUGCUCGAUGAGUUUUUCUAA | pJK_ToeRep_kanR_sensor |
| bla sensor | GGGAAACAGGAAGGCAAAAUGCCGCAAAAAAGGGAAUAAGGGCG<br>ACACGGAAAUGUUGAAUACAAAUCAACAGUAUUCUACAUUUCC<br>GUGUCGCACUUAUUCAAACAGAGGAGAGAAUAAAUGCGACACG<br>GAAACCUGGCGGCAGCGCAAAAGAUGCGUAAA | GUAUUCAACAUUUCCGUGUCGCCCUUAUUCC<br>CUUUUUUGCGGCAUUUUGCCUUCCUGUUU | AUGAGUAUUCAACAUUUCCGUGUCGCCCUUAUUCCCUUUUUUGCGGCAUUUUGCCUUCCUGUUUUUGCUCACCCAGAAACGCUGGUGAAAGUAAAAGAUGCUGA<br>AGAUCAGUUGGGUGCACGAGUGGGUUACAUCGAACUGGAUCUCAACAGCGGUAAGAUCCUUGAGAGUUUUCGCCCCGAAGAACGUUUUCCAAUGAUGAGCACUU<br>UUAAAGUUCUGCUAUGUGGCGCGGUAUUAUCCCGUGUUGACGCCGGGCAAGAGCAACUCGGUCGCCGCAUACACUAUUCUCAGAAUGACUUGGUUGAGUACUCA<br>CCAGUCACAGAAAAGCAUCUUACGGAUGGCAUGACAGUAAGAGAAUUAUGCAGUGCUGCCAUAACCAUGAGUGAUAACACUGCGGCCAACUUACUUCUGACAACG<br>AUCGGAGGACCGAAGGAGCUAACCGCUUUUUUGCACAACAUGGGGGAUCAUGUAACUCGCCUUGAUCGUUGGGAACCGGAGCUGAAUGAAGCCAUACCAAACGAC<br>GAGCGUGACACCACGAUGCCUGCAGCAAUGGCAACAACGUUGCGCAAACUAUUAACUGGCGAACUACUUACUCUAGCUUCCCGGCAACAAUUAAUAGACUGGAUG<br>GAGGCGGAUAAAGUUGCAGGACCACUUCUGCGCUCGGCCCUUCCGGCUGGCUGGUUUAUUGCUGAUAAAUCUGGAGCCGGUGAGCGUGGGUCUCGCGGUAUCA<br>UUGCAGCACUGGGGCCAGAUGGUAAGCCCUCCCGUAUCGUAGUUAUCUACACGACGGGGAGUCAGGCAACUAUGGAUGAACGAAAUAGACAGAUCGCUGAGAUAG<br>GUGCCUCACUGAUUAAGCAUUGGUAA | pJK_ToeRep_bla_sensor |

**B. 3WJ Repressor mRNA Sensors**

| mRNA Sensor | Sensor Sequence | mRNA Target Subsequence | Full mRNA Target Sequence | Switch Plasmid |
|---|---|---|---|---|
| KanR sensor | GGGAGGACAAUUACAAACUUGUUAUAGUUAUGAACAGAGGAGA<br>CAUAACAUGAACAAAGGAAUCGAAUGGAAUAAGAACCUGGCGGC<br>AGCGCAAAAGAUGCGUAAA | CAUUCGAUUCCUGUUUGUAAUUGUCCU | AUGAGCCAUAUUCAACGGGAAACGUCUUGCUCUAGGCCGCGAUUAAAUUCCAACAUGGAUGCUGAUUUAUAUGGGUAUAAAUGGGCUCGCGAUAAUGUCGGGCAA<br>UCAGGUGCGACAAUCUAUCGAUUGUAUGGGAAGCCCGAUGCGCCAGAGUUGUUUCUGAAACAUGGCAAAGGUAGCGUUGCCAAUGAUGUUACAGAUGAGAUGGU<br>CAGACUAAACUGGCUGACGGAAUUUAUGCCUCUUCCGACCAUCAAGCAUUUUAUCCGUACUCCUGAUGAUGCAUGGUUACUCACCACUGCGAUCCCCGGGAAAA<br>CAGCAUUCCAGGUAUUAGAAGAAUAUCCUGAUUCAGGUGAAAAUAUUGUUGAUGCGCUGGCAGUGUUCCUGCGCCGGUUGCAUUCGAUUCCUGUUUGUAAUUGU<br>CCUUUUAACAGCGAUCGCGUAUUUCGUCUCGCUCAGGCGCAAUCACGAAUGAAUAACGGUUUGGUUGAUGCGAGUGAUUUUGAUGACGAGCGUAAUGGCUGGCC<br>UGUUGAACAAGUCUGGAAAGAAAUGCAUAAACUUUUGCCAUUCUCACCGGAUUCAGUCGUCACUCAUGGUGAUUUCUCACUUGAUAACCUUAUUUUUGACGAGG<br>GGAAAUUAAUAGGUUGUAUUGAUGUUGGACGAGUCGGAAUCGCAGACCGAUACCAGGAUCUUGCCAUCCUAUGGAACUGCCUCGGUGAGUUUUCUCCUUCAUUA<br>CAGAAACGGCUUUUUCAAAAAUAUGGUAUUGAUAAUCCUGAUAUGAAUAAAUUGCAGUUUCAUUUGAUGCUCGAUGAGUUUUUCUAA | pYZ_3WJrep_kanR_sensor |
| aadA sensor | GGGUUAGCGCCUCAAAUAUUGUUAUAGUUAUGAACAGAGGAGA<br>CAUAACAUGAACAAGAUCCUGUUCAGCACAUUCAACCUGGCGG<br>CAGCGCAAAAGAUGCGUAAA | CUGAACAGGAUCUAUUUGAGGCGCUAA | AUGAGGGAAGCGGUGAUCGCCGAAGUAUCGACUCAACUAUCAGAGGUAGUUGGCGUCAUCGAGCGCCAUCUCGAACCGACGUUGCUGGCCGUACAUUUGUACGG<br>CUCCGCAGUGGAUGGCGGCCUGAAGCCACACAGUGAUAUUGAUUUGCUGGUUACGGUGACCGUAAGGCUUGAUGAAACAACGCGGCGAGCUUUGAUCAACGACC<br>UUUUGGAAACUUCGGCUUCCCCUGGAGAGAGCGAGAUUCUCCGCGCUGUAGAAGUCACCAUUGUUGUGCACGACGACAUCAUUCCGUGGCGUUAUCCAGCUAAG<br>CGCGAACUGCAAUUUGGAGAAUGGCAGCGCAAUGACAUUCUUGCAGGUAUCUUCGAGCCAGCCACGAUCGACAUUGAUCUGGCUAUCUUGCUGACAAAAGCAAGA<br>GAACAUAGCGUUGCCUUGGUAGGUCCAGCGGCGGAGGAACUCUUUGAUCCGGUUCCUGAACAGGAUCUAUUUGAGGCGCUAAAUGAAACCUUAACGCUAUGGAA<br>CUCGCCGCCCGACUGGGCUGGCGAUGAGCGAAAUGUAGUGCUUACGUUGUCCCGCAUUUGGUACAGCGCAGUAACCGGCAAAAUCGCGCCGAAGGAUGUCGCU<br>GCCGACUGGGCAAUGGAGCGCCUGCCGGCCCAGUAUCAGCCCGUCAUACUUGAAGCUAGACAGGCUUAUCUUGGACAAGAAGAAGAUCGCUUGGCCUCGCGCGC<br>AGAUCAGUUGGAAGAAUUUGUCCACUACGUGAAAGGCGAGAUCACCAAGGUAGUCGGCAAAUAA | pYZ_3WJrep_aadA_sensor |

**Table S8. NAND Gate and NOR Gate Circuit RNA Sequences**

**NOTES:**

1. Input or trigger RNA sequences are listed up to the base immediately before the T7 terminator used to terminate transcription.
2. Gate RNA sequences are listed up to the 30th base (inclusive) following the end of their trigger binding domain. This 30-nt sequence contains the 21-nt linker sequence and the first 9-nts of GFPmut3b.

**A. 2-input Toehold Repressor NAND Gate Circuit**

| Gate RNA | Gate Plasmid | Gate Plasmid Origin/Resistance | Input A1 Sequence | Input A2 Sequence | Non-Cognate RNA C Sequence | Non-Cognate RNA D Sequence | Triggers for Input 11 | Triggers for Input 10 | Triggers for Input 01 | Triggers for Input 00 |
|---|---|---|---|---|---|---|---|---|---|---|
| GGGAAGAUGAAAGAAGAUGGUAUGGUGAAGAGGUGUAAAGAUACACAAUACAACAA<br>CUAUCUUUCCACCUCUCCACCAUACAAACAGAGGAGAUGGAGAAUGGGAAAGAUAA<br>ACCUGGCGGCAGCGCAAAAGAUGCGUAAA | pJK_NNAND1_RR07 | ColA/kanamycin | GGGCCUAUGGUGAU<br>UGUGCUCACCAUAGG<br>GACUAUCUUUACACC<br>UCUUCACCAUACACG<br>AUAAACGACGAUACA<br>AAUGACUA | GGGUCAUGCCAGC<br>UUCAUUGCUGGCA<br>UGAAUUUCAUUUG<br>UAUCUUCGUUUAU<br>CGUCAUCUUCUUU<br>CAUCUUCAC | GGGCCAAGAACCGGUU<br>UCACGGUUCUUGGACU<br>UAUCUUCACACCAUUC<br>UACAACUAAACUAAGC<br>UUGCGCGUCUCAUA | GGGUCUCACGCCCUC<br>AGCUGGGCGUGAGAU<br>GAGCCUCGUCUCCAG<br>AUGACGAGGCAACGUA<br>GGAUCUGACUGAUCC<br>UACUAU | A1, A2 | A1, D | C, A2 | C, D |

**B. 2-input 3WJ Repressor NAND Gate Circuit**

| Gate RNA | 3WJ Repressor Switch Indices in Gate | Gate Plasmid | Gate Plasmid Origin/Resistance | 3WJ Repressor Trigger Indices for Input 11 | 3WJ Repressor Trigger Indices for Input 10 | 3WJ Repressor Trigger Indices for Input 01 | 3WJ Repressor Trigger Indices for Input 00 | Input Plasmid Origin/Resistance |
|---|---|---|---|---|---|---|---|---|
| GGGACUAAUCAGAUCUACUUGUUAUAGUUAUGAACAGAGGAG<br>ACAUAACAUGAACAAGCACCUAACAAGACUAAUCAUAUACAAU<br>ACCUCCAAAUCAAUCAAUUCUACUUGUUAUAGUUAUGAACAG<br>AGGAGACAUAACAUGAACAAUCACAUAAACCUAAGAACUAACC<br>UGGCGGCAGCGCAAAAGAUGCGUAAA | 19, 11 | pYZ_NAND2_L17_S19_S11 | ColA/kanamycin | 19, 11 | 19, 12 | 11, 12 | 12, 12 | ColE1/ampicillin,<br>CDF/spectinomycin |

**C. 2-Input 3WJ Repressor NOR Gate Circuit**

| Gate RNA | 3WJ Repressor Switch Index in Gate | Gate Plasmid | Gate Plasmid Origin/Resistance | Trigger A Sequence | Trigger B Sequence | Non-Cognate RNA C Sequence | Non-Cognate RNA D Sequence | Triggers for Input 11 | Triggers for Input 10 | Triggers for Input 01 | Triggers for Input 00 |
|---|---|---|---|---|---|---|---|---|---|---|---|
| GGGAGUAAAGUAUAGAACGCUUGAUGUUGAUGUUAGCGACAC<br>UAAUAGUGUAGUGUCGGAUAAGUGAAUAUAGUCUAACAUCAA<br>CAUCAAGCGCCUCAAUUCGAGGCAAGACUAAGACUAAAUGAU<br>GUAUAGAUUGAAGAUUGCGACACUAAUAGUGUAGUGUCGGAU<br>AAGUGAAUAGUAGUCAAUCUUCAAUCUAUACAGCCUCAAUUC<br>GAGGCAAGACUACUAUUCACUUAUUGUUAUAGUUAUGAACAG<br>AGGAGACAUAACAUGAACAAUCCGACACUACAAUUUCGAAACC<br>UGGCGGCAGCGCAAAAACCUGGCGGCAGCGCAAAAGAUGCGU<br>AAA | 21 | pYZ_NOR2_HpEF | ColA/kanamycin | GGGUGACAUAUCCAA<br>GAGCGGAUAUGUCAC<br>UAACAUCAACAUCAA<br>GCGUUCUAUACUUUA<br>CU | GGGACGUGCGCAG<br>CUCCUACUGCGCA<br>CGUCAAUCUUCAA<br>UCUAUACAUCAUU<br>UAGUCUUAGU | GGGACGAUAUACGGGU<br>CUACGUAUAUCGUACA<br>AGAACCAUACAAGACAA<br>CGACGACACUAGA | GGGACAUCAGAGACUA<br>GUAUCUCUGAUGUAU<br>AAGACAAGACAUAAGA<br>ACAGAAAUACACAGA | A, B | A, C | B, C | C, D |

**D. 3-Input 3WJ Repressor NAND Gate Circuit**

| Gate RNA | 3WJ Repressor Switch Indices | Gate Plasmid | Gate Plasmid Origin/Resistance | 3WJ Repressor Trigger Indices for Input 111 | 3WJ Repressor Trigger Indices for Input 011 | 3WJ Repressor Trigger Indices for Input 101 | 3WJ Repressor Trigger Indices for Input 110 | 3WJ Repressor Trigger Indices for Input 100 | 3WJ Repressor Trigger Indices for Input 010 | 3WJ Repressor Trigger Indices for Input 001 | 3WJ Repressor Trigger Indices for Input 000 |
|---|---|---|---|---|---|---|---|---|---|---|---|
| GGGAUCAAUCAAUUCUACUUGUUAUAGUUAUGAACAGAGGAG<br>ACAUAACAUGAACAAUCACAUAAACCUAAGAACUCAAUAUCGC<br>AUACAAUCAAAUACAAAUUGUUAUAGUUAUGAACAGAGGAGAC<br>AUAACAUGAACAAUCACAUACAAAGCAAACGACAAAUACAAAUA<br>CUAAUCAGAUCUACUUGUUAUAGUUAUGAACAGAGGAGACAU<br>AACAUGAACAAGCACCUAACAAGACUAAUCAACCUGGCGGCAG<br>CGCAAAAGAUGCGUAAAGGAGAAGAACUUUUCACUGGAACCU<br>GGCGGCAGCGCAAAAGAUGCGUAAA | 11, 13, 19 | pYZ_NAND3_L11_S11_S13_S19 | ColA/kanamycin | (11, 19), 13 | (13, 12), 19 | 11, (19, 12) | 11, (13, 12) | (11, 12), 12 | (12, 12), 19 | (12, 12), 13 | (12, 12), 12 |

**E. 4-Input 3WJ Repressor NAND Gate Circuit**

| Gate RNA | 3WJ Repressor Switch Indices | Gate Plasmid | Gate Plasmid Origin/Resistance | 3WJ Repressor Trigger Indices for Input 1111 | 3WJ Repressor Trigger Indices for Input 0111 | 3WJ Repressor Trigger Indices for Input 1011 | 3WJ Repressor Trigger Indices for Input 1101 | 3WJ Repressor Trigger Indices for Input 1110 | 3WJ Repressor Trigger Indices for Input 1100 | 3WJ Repressor Trigger Indices for Input 1010 | 3WJ Repressor Trigger Indices for Input 1001 |
|---|---|---|---|---|---|---|---|---|---|---|---|
| GGGACUAAUCAGAUCUACUUGUUAUAGUUAUGAACAGAGGAG<br>ACAUAACAUGAACAAGCACCUAACAAGACUAAUCAACUACAAA<br>UACAAUAACUCCUAUCACUUUACUUGUUAUAGUUAUGAACAG<br>AGGAGACAUAACAUGAACAAGCACACUAACUACAAAUUCACAA<br>UUACCGACGCAAUACAAUCAAAUACAAAUUGUUAUAGUUAUGA<br>ACAGAGGAGACAUAACAUGAACAAUCACAUACAAAGCAAACGA<br>AAUACCGCAAAUACCAUAUCAAUCAAUUCUACUUGUUAUAGU<br>UAUGAACAGAGGAGACAUAACAUGAACAAUCACAUAAACCUAA<br>GAACUAACCUGGCGGCAGCGCAAAACCUGGCGGCAGCGCAAA<br>AGAUGCGUAAA | 24, 11, 19, 13 | pYZ_NAND4_L17_S4_S11_S19_S13 | ColA/kanamycin | (11, 13), (19, 24) | 12, (13, 19, 24) | 12, (11, 19, 24) | 12, (11, 13, 24) | 12, (11, 13, 19) | (12, 12), (11, 13) | (12, 12), (11, 19) | (12, 12), (11, 24) |

**F. NOT ((A1 AND A2) OR (B1 AND B2)) Circuit**

| Gate RNA | Gate Plasmid | Gate Plasmid Origin/Resistance | Input A1 Sequence | Input A2 Sequence | Input B1 Sequence | Input B2 Sequence | Non-Cognate RNA C Sequence | Non-Cognate RNA D Sequence | Triggers for Input 11 | Triggers for Input 10 |
|---|---|---|---|---|---|---|---|---|---|---|
| GGGAAGAUGAAAGAAGAUGGUAUGGUGAAGAGGUGUAAAGAUACACAAUACAACAA<br>CUAUCUUUCCACCUCUCCACCAUACAAACAGAGGAGAUGGAGAAUGGGAAAGAUAA<br>ACCUGGCGGCAGCGCAAAAGAUGCGUAAA | pJK_NNAND1_RR07 | ColA/kanamycin | GGGCCUAUGGUGAU<br>UGUGCUCACCAUAGG<br>GACUAUCUUUACACC<br>UCUUCACCAUACACG<br>AUAAACGACGAUACA<br>AAUGACUA | GGGUCAUGCCAGC<br>UUCAUUGCUGGCA<br>UGAAUUUCAUUUG<br>UAUCUUCGUUUAU<br>CGUCAUCUUCUUU<br>CAUCUUCAC | GGGAGCAGAGGUUCGU<br>ACUAACCUCUGCUACU<br>UAUCUUUACACCUCUU<br>CACCUGAUUAACGCGC<br>ACUAUGAUUGACAC | GGGACAUGAUAAGACA<br>UGCCUUAUCAUGUCA<br>UUCAAUCAUAGUACGC<br>GUUAAUCAAUACCAUC<br>UUCUUUCAUCUUAUC | GGGCCAAGAACCG<br>GUUUCACGGUUCU<br>UGGACUUAUCUUC<br>ACACCAUUCUACAA<br>CUAAACUAAGCUU<br>GCGCGUCUCAUA | GGGUCUCACGCC<br>CUCAGCUGGGCG<br>UGAGAUGAGCCU<br>CGUCUCCAGAUG<br>ACGAGGCAACGUA<br>GGAUCUGACUGA<br>UCCUACUAU | A1, A2 | A1, D |

| Trigger Plasmid Origin/Resistance |
| --- |
| ColE1/ampicillin, CDF/spectinomycin |

| Trigger Plasmid Origin/Resistance |
| --- |
| ColE1/ampicillin, CDF/spectinomycin |

| Input Plasmid Origin/Resistance |
| --- |
| ColE1/ampicillin, CDF/spectinomycin |

| 3WJ Repressor Trigger Indices for Input 0110 | 3WJ Repressor Trigger Indices for Input 0101 | 3WJ Repressor Trigger Indices for Input 0011 | 3WJ Repressor Trigger Indices for Input 1000 | 3WJ Repressor Trigger Indices for Input 0100 | 3WJ Repressor Trigger Indices for Input 0010 | 3WJ Repressor Trigger Indices for Input 0001 | 3WJ Repressor Trigger Indices for Input 0000 | Input Plasmid Origin/Resistance |
| --- | --- | --- | --- | --- | --- | --- | --- | --- |
| (12, 12), (13, 19) | (12, 12), (13, 24) | (12, 12), (19, 24) | (12, 12), (11, 12) | (12, 12), (13, 12) | (12, 12), (19, 12) | (12, 12), (24, 12) | (12, 12), (12, 12) | ColE1/ampicillin, CDF/spectinomycin |

| Triggers for Input 01 | Triggers for Input 00 | Triggers for Input 0011 | Triggers for Input 0001 | Triggers for Input 0010 | Triggers for Input 1100 | Triggers for Input 0100 | Triggers for Input 1000 | Triggers for Input 1010 | Triggers for Input 0110 | Triggers for Input 1001 | Triggers for Input 0101 | Triggers for Input 0000 | Trigger Plasmid Origin/Resistance |
| --- | --- | --- | --- | --- | --- | --- | --- | --- | --- | --- | --- | --- | --- |
| C, A2 | C, D | B1, B2 | C, B2 | B1, D | A1, A2 | C, A2 | A1, C | A1, B1 | A2, B1 | A1, B2 | A2, B2 | C, D | ColE1/ampicillin, CDF/spectinomycin |

**Table S9. Descriptions of Thermodynamic Parameters Used for Automated Forward Engineering**

| Group | Parameter Name | Parameter Description |
|---|---|---|
| **MFE of RNA strands and critical RNA subsequences** | deltaG_hpin | The minimum free energy (MFE) of the switch (or hairpin) RNA running from the 5' end to 29th base of the GFP coding sequence. |
| | deltaG_targ | The MFE of the trigger (or target) RNA from the 5' end to the end of the terminator sequence. |
| | deltaG_comp | The MFE of the complex formed between the switch RNA sequence defined above and the full trigger RNA. |
| | deltaG_min_hpin | The MFE of the minimal switch RNA sequence from the 5' end to the base immediately before the start of the 21-nt linker sequence. |
| | deltaG_min_targ | The MFE of the 45-nt minimal trigger RNA sequence that is programmed to bind to the switch RNA. |
| | deltaG_min_comp | The MFE of the complex formed between the minimal switch RNA sequence and the minimal trigger RNA sequence. |
| | deltaG_min_stem_recon | The MFE of the hairpin secondary structure that forms to repress translation once the trigger RNA binds; this sequence comprises the 18-nt repressing stem along with the RBS and start codon. |
| **Measures of binding for critical RNA domains** | deltaG_toeh_binding | The free energy of the 15-nt toehold sequence of the repressor when base pairing perfectly to its reverse complement. |
| | deltaG_targ_binding | The free energy of the 45-nt minimal trigger RNA sequence when base pairing perfectly to its reverse complement sequence |
| | deltaG_toeh_binding_actual | The MFE of the 15-nt toehold sequence of the repressor bound to its reverse complement sequence; this MFE structure may have unpaired bases at the ends of the duplex. |
| | deltaG_targ_binding_actual | The MFE of the 45-nt minimal trigger RNA sequence bound to its reverse complement sequence; this MFE structure may have unpaired bases at the ends of the duplex. |
| **Net reaction free energies** | net_deltaG_comp | The net free energy change upon formation of the trigger/switch RNA complex; it is equal to deltaG_comp - deltaG_hpin - deltaG_targ. |
| | net_deltaG_min_comp | The net free energy change upon formation of the trigger/switch RNA complex based on the minimal trigger and switch sequences; it is equal to deltaG_min_comp - deltaG_min_hpin - deltaG_min_targ. |
| **Influence of coding sequence secondary structure on translation in active state** | dup2mRNA_deltaG | The MFE of the RNA sequence starting from the first base after the switch RNA stem (nucleotide 88) through to the 29th base of the GFP coding sequence |
| | dup2link_deltaG | The MFE of the RNA sequence starting from the first base after the switch RNA stem (nucleotide 88) through to the end of the 21-nt linker sequence (nucleotide 138, just before the begininning of the GFP sequence). |
| | dup2pos03_deltaG | The MFE of the RNA sequence starting from the first base after the switch RNA stem (nucleotide 88) through to the Nth base of the GFP coding sequence, where N = 3, 6, 9, etc. |
| | dup2pos06_deltaG | |
| | dup2pos09_deltaG | |
| | dup2pos12_deltaG | |
| | dup2pos15_deltaG | |
| | dup2pos18_deltaG | |
| | dup2pos21_deltaG | |
| | dup2pos24_deltaG | |
| | dup2pos27_deltaG | |
| | AUG2mRNA_deltaG | The MFE of the RNA sequence starting from the start codon of the switch RNA (nucleotide 106) through to the 29th base of the GFP coding sequence. |
| | dup2AUG_deltaG | The MFE of the RNA sequence starting from the first base after the switch RNA stem (nucleotide 88) through to the end of the switch RNA start codon (nucleotide 108). |
| **Influence of coding sequence secondary structure on translation in inactive state** | recon_dup2mRNA_deltaG | The MFE of the RNA sequence starting from the first base after the trigger RNA binding site on the switch RNA (nucleotide 49) through to the 29th base of the GFP coding sequence. |
| | recon_dup2link_deltaG | The MFE of the RNA sequence starting from the first base after the trigger RNA binding site on the switch RNA (nucleotide 49) through to the end of the 21-nt linker sequence (nucleotide 138, just before the begininning of the GFP sequence). |
| | recon_dup2pos03_deltaG | The MFE of the RNA sequence starting from the first base after the trigger RNA binding site on the switch RNA (nucleotide 49) through to the Nth base of the GFP coding sequence, where N = 3, 6, 9, etc. |
| | recon_dup2pos06_deltaG | |
| | recon_dup2pos09_deltaG | |
| | recon_dup2pos12_deltaG | |
| | recon_dup2pos15_deltaG | |
| | recon_dup2pos18_deltaG | |
| | recon_dup2pos21_deltaG | |
| | recon_dup2pos24_deltaG | |
| | recon_dup2pos27_deltaG | |
| **Deviations of actual sequence from the ideal, design-specified secondary structure** | dev_deltaG_hpin | The difference in energy obtained by subtracting deltaG_hpin from the free energy of the switch RNA sequence when it is folded in the ideal, design-specified secondary structure. |
| | dev_deltaG_targ | The difference in energy obtained by subtracting deltaG_targ from the free energy of the trigger RNA sequence when it is folded in the ideal, design-specified secondary structure. |
| | dev_deltaG_comp | The difference in energy obtained by subtracting deltaG_comp from the free energy of the trigger/switch RNA complex when it is folded in the ideal, design-specified secondary structure. |
| | dev_deltaG_min_hpin | The difference in energy obtained by subtracting deltaG_min_hpin from the free energy of the minimal switch RNA sequence when it is folded in the ideal, design-specified secondary structure. |
| | dev_deltaG_min_targ | The difference in energy obtained by subtracting deltaG_min_targ from the free energy of the minimal trigger RNA sequence when it is folded in the ideal, design-specified secondary structure. |

| | | |
|---|---|---|
| | dev_deltaG_min_comp | The difference in energy obtained by subtracting deltaG_min_comp from the free energy of the trigger/switch RNA complex generated from the minimal sequences when it is folded in the ideal, design-specified secondary structure. |
| | dev_recon_dup2mRNA_deltaG | The difference in energy obtained by subtracting recon_dup2mRNA_deltaG from the free energy of that sequence (i.e. the one used for calculating recon_dup2mRNA_deltaG) in the ideal, design-specified secondary structure, which contains a translation-repressing hairpin structure. |
| | dev_recon_dup2link_deltaG | The difference in energy obtained by subtracting recon_dup2link_deltaG from the free energy of that sequence in the ideal, design-specified secondary structure. |
| | dev_recon_dup2pos03_deltaG | The difference in energy obtained by subtracting recon_dup2posN_deltaG from the free energy of that sequence in the ideal, design-specified secondary structure, where N = 3, 6, 9, etc. |
| | dev_recon_dup2pos06_deltaG | |
| | dev_recon_dup2pos09_deltaG | |
| | dev_recon_dup2pos12_deltaG | |
| | dev_recon_dup2pos15_deltaG | |
| | dev_recon_dup2pos18_deltaG | |
| | dev_recon_dup2pos21_deltaG | |
| | dev_recon_dup2pos24_deltaG | |
| | dev_recon_dup2pos27_deltaG | |
| **Measures of switch RNA stem sequence and secondary structure** | deltaG_stem_01 | This set of parameters was calculated starting from the 69-nt sequence of the main switch RNA hairpin, which consisted of a 30-nt stem and a 9-nt loop. The main switch RNA hairpin was then analyzed for different subsequences corresponding to different stem lengths. deltaG_stem_01 is the MFE of nucleotides 1 to 69, deltaG_stem_02 is the MFE of nucleotides 2 to 68, etc. |
| | deltaG_stem_02 | |
| | deltaG_stem_03 | |
| | deltaG_stem_04 | |
| | deltaG_stem_05 | |
| | deltaG_stem_06 | |
| | deltaG_stem_07 | |
| | deltaG_stem_08 | |
| | deltaG_stem_09 | |
| | deltaG_stem_10 | |
| | deltaG_stem_11 | |
| | deltaG_stem_12 | |
| | deltaG_stem_13 | |
| | deltaG_stem_14 | |
| | deltaG_stem_15 | |
| | deltaG_stem_16 | |
| | deltaG_stem_17 | |
| | deltaG_stem_18 | |
| | deltaG_stem_19 | |
| | deltaG_stem_20 | |
| | deltaG_stem_21 | |
| | deltaG_stem_22 | |
| | deltaG_stem_23 | |
| | deltaG_stem_24 | |
| | deltaG_stem_25 | |
| | deltaG_stem_26 | |
| | deltaG_stem_27 | |
| | deltaG_stem_28 | |
| | deltaG_stem_29 | |
| | deltaG_stem_30 | |
| | min_bot_deltaG_01 | This set of parameters was calculated using the 5' and 3' arms of the main switch RNA hairpin. The 5' and 3' arms comprised nucleotides 19 to 48 and 58 to 87 of the switch RNA sequence, respectively. For the purposes of the calculation, subsequences from the 5' and 3' arms were joined using the sequence AAAAAAAAAA and their MFE computed. min_bot_deltaG_02, for instance, has the 1st through 29th bases of the 5' arm, the poly-A loop, and the 2nd through 30th bases of the 3' arm. |
| | min_bot_deltaG_02 | |
| | min_bot_deltaG_03 | |
| | min_bot_deltaG_04 | |
| | min_bot_deltaG_05 | |
| | min_bot_deltaG_06 | |
| | min_bot_deltaG_07 | |
| | min_bot_deltaG_08 | |
| | min_bot_deltaG_09 | |
| | min_bot_deltaG_10 | |
| | min_bot_deltaG_11 | |
| | min_bot_deltaG_12 | |
| | min_bot_deltaG_13 | |
| | min_bot_deltaG_14 | |
| | min_bot_deltaG_15 | |
| | min_bot_deltaG_16 | |
| | min_bot_deltaG_17 | |
| | min_bot_deltaG_18 | |
| | min_bot_deltaG_19 | |
| | min_bot_deltaG_20 | |
| | min_bot_deltaG_21 | |
| | min_bot_deltaG_22 | |
| | min_bot_deltaG_23 | |
| | min_bot_deltaG_24 | |
| | min_bot_deltaG_25 | |
| | min_bot_deltaG_26 | |
| | min_bot_deltaG_27 | |
| | min_bot_deltaG_28 | |
| | min_bot_deltaG_29 | |
| | min_bot_deltaG_30 | |
